# Time-resolved effects of cold atmospheric plasma on *E. coli* GW-AmxH19 transcriptome and proteome in an emulated wastewater environment

**DOI:** 10.1101/2025.05.08.652849

**Authors:** Jörg Bernhardt, Dominik Schneider, Julia Rielicke, Susanne Sievers, Veronika Hahn, Mechthild Bömeke, Anja Poehlein, Dirk Albrecht, Juergen F. Kolb, Katharina Riedel, Rolf Daniel, Daniela Zühlke

## Abstract

Cold atmospheric plasma (CAP) has been shown to be effective against a variety of microorganisms. In this study, we described effects on an *Escherichia coli* strain isolated from hospital wastewater caused by a treatment with physical plasma. *E. coli* GW-AmxH19 was incubated in artificial wastewater and treated for 15 minutes with CAP. Transcriptomes and proteomes were monitored at different timepoints within a 24 h period to differentiate between immediate physiological responses and adaptations in the recovery phase. Reduction of viable cells was on average at 90%. The short-term response of the surviving cells to physical plasma aims at repairing and protecting cellular structures from plasma-induced damages, mainly provoked by reactive oxygen and nitrogen species. Notably, CAP induced a temporary transcription of genes from a conjugative plasmid carrying antibiotic resistance determinants. The late response during recovery phase is dominated by a massive activation of two prophages turning cold plasma treatment into a novel possible strategy to induce the lytic cycle of prophages. This study is the first report on the combined analysis of transcriptional and translational effects of CAP on an environmental bacterial isolate in a time-resolved manner. Chances and risks of considering CAP as an additional purification step in wastewater treatment plants are depicted and discussed.

## 1 Introduction

Antimicrobial resistance (AMR) is one of the top global public health threats. Infectious diseases caused by (multi-)resistant bacteria become increasingly difficult to treat effectively, and sometimes even antibiotics of last resort may become useless. In 2019, nearly five million deaths were associated with bacterial AMR and one million deaths were directly attributed with AMR (Murray et al., 2022); estimates suggest ten million AMR-related deaths by the year 2050 (O’Neill, 2014). Thus, preventing the emergence and spread of antibiotic-resistant bacteria is of global importance. One hot spot for the emergence and dissemination of antibiotic-resistant bacteria are wastewater treatment plants, receiving sewage from private households and hospitals (Rizzo et al., 2013). Besides harmless microbes these sewage waters contain potentially pathogenic and/or antibiotic-resistant bacteria as well as antibiotic residues in varying concentrations. Nutrient-rich conditions and high bacterial density due to microbial organization in free biofilm entities (bacterial flocs) in wastewater treatment plants favour horizontal gene transfer (HGT) and thus might also promote exchange of antibiotic resistance genes (Stalder et al., 2012). In addition, residues of antibiotics present in wastewater (Coutu et al., 2013; Bengtsson-Palme et al., 2016; Wang et al., 2020; Zhang et al., 2024) might favour selection for these resistant bacteria even in sub-inhibitory concentrations und thus outcompete sensitive bacteria (Gullberg et al., 2011). Although the bacterial load is generally reduced during the treatment process, “problematic” microbes may survive the process and are released with the effluent into the receiving waters and thus spread in the aquatic environment (Czekalski et al., 2012; Marti et al., 2014; Bengtsson-Palme et al., 2016; Narciso-da-Rocha et al., 2018; Alexander et al., 2020). Hence, pathogenic and antibiotic-resistant bacteria are potentially (re-)transmitted to humans and animals (Zhou et al., 2018). To further reduce the bacterial load in treated wastewater and in order to reduce the risk of spreading resistant bacteria additional treatment strategies are investigated or already implemented in wastewater treatment plants. Besides filtration and chlorination, so-called advanced oxidation processes (AOP) like ozone or UV irradiation are explored. These approaches have been proven to reduce the load of antibiotic-resistant bacteria in wastewater (Guo et al., 2013; Czekalski et al., 2016; Zheng et al., 2017; Hembach et al., 2019; Rodríguez-Chueca et al., 2019). Additionally, there is evidence that sub-lethal UV treatment leads to a reduction in antibiotic resistance genes (Zheng et al., 2017). As an AOP, cold atmospheric plasma (CAP) can also be successfully applied for the degradation of microbial and chemical contaminants in wastewater. Thus, plasma treatment has been demonstrated effective in the inactivation of bacteria and fungi (Hahn et al., 2019; Handorf et al., 2019) including antibiotic-resistant microorganisms (Daeschlein et al., 2014; Li et al., 2021). Furthermore, the degradation of pharmaceuticals like antibiotics (Terefinko et al., 2023) or recalcitrant pollutants such as perfluorooctanoic acid (PFOA) (Buckstöver et al., 2025) has also been shown with plasma. Physical plasma can generate highly reactive but short-lived species (especially hydroxyl radicals), which allow for a destruction of particularly persistent substances, such as X-ray contrast agents and resilient bacteria. The approach is therefore often more efficient than other AOP like ozonation and UV irradiation (Bruggeman et al., 2016; Banaschik et al., 2017, 2018). Large-scale wastewater treatment with plasma is in principle possible. There are already systems available for the treatment of several hundred liters of water per day. However, so far little is known about the potential and possible risks of plasma-based wastewater decontamination. Plasma is generated only by the supply of energy; no chemicals or additives are needed. The inactivation of microbes by plasma originates from a plethora of reactive oxygen and nitrogen species (RONS), such as singlet oxygen, hydroxyl radicals, superoxide anion, nitric oxide, and the compounds formed from these RONS, e.g. peroxynitrous acid (Lukes et al., 2014; Balazinski et al., 2021). This results in the oxidation of biomolecules like lipids, lipopolysaccharides, proteins, DNA, or RNA and thus is detrimental for microorganisms. The effects of physical plasma on Gram-positive and -negative model bacteria and their cellular responses were analyzed in several studies (Sharma et al., 2009; Winter et al., 2011, 2013; Yost and Joshi, 2015; Pan et al., 2022; Liew et al., 2023). As a protection mechanism to the toxic effects of RONS bacteria express plasma-protective genes that lead to the survival of longer plasma exposure (Krewing et al., 2019). However, so far, no specific resistance mechanisms against the fundamental physical and oxidative action of plasma has been observed.

Recently, we investigated short-term physiological changes in the proteome of the hospital wastewater-borne *Escherichia coli* strain GW-AmxH19 after cold atmospheric plasma treatment (Hahn et al., 2024). A response to oxidative stress provoked by ROS and RNS and pH-balancing reactions were the key proteomic signatures observed after plasma treatment. To mimic the conditions of a real plasma treatment of wastewater, we here performed experiments in artificial wastewater, also including the recovery phase of *E. coli* GW-AmxH19 by studying long-termed effects after re-growth of bacteria surviving the treatment. To obtain a comprehensive perspective on the physiological changes and adaptation mechanisms, we analyzed transcriptome (RNA abundance) and proteome (protein abundance) and collected data at certain intervals over 24 h after treatment which advanced our mechanistic understanding of the action of physical plasma on bacteria.

## 2 Materials and Methods

### 2.1 Hospital effluent derived isolate *Escherichia coli* GW-AmxH19

The *E. coli* strain GW-AmxH19 was isolated from wastewater of the hospital in Greifswald (Germany) in March, 2017. The Gram-negative bacterium is mesophilic and has a genome consisting of two replicons, a circular chromosome (5.03 Mbp) and a circular plasmid (126.96 kbp) with in total 4,795 protein-coding genes (Schneider et al., 2020). The isolate is closely related to uropathogenic *E. coli* and contains genes conferring potential resistance to antibiotics, such as resistance to aminoglycosides, beta-lactams, fluoroquinolones, macrolides, polymyxin. Furthermore, PHASTEST analysis (Wishart et al., 2023) revealed the presence of two complete and two incomplete prophage regions in the genome of this strain (Supplementary Figure S6).

### 2.2 Precultivation of *E. coli* GW-AmxH19

*E. coli* GW-AmxH19 was inoculated in 20 mL LB medium (20 g/L) and incubated overnight with shaking at 37°C for 20 hours. The optical density (OD) was determined at 500 nm. The entire culture was centrifuged at room temperature at 4,700 rpm for 5 minutes, the supernatant was discarded, and the pellet was resuspended with 20 mL of 0.85% NaCl solution. This washing procedure was repeated twice.

### 2.3 Experimental setup and treatment of *E. coli* GW-AmxH19

To emulate the conditions of a wastewater treatment plant the washed bacterial cell pellet was used to inoculate synthetic wastewater. The composition of synthetic wastewater is proposed by the OECD guidelines and consists of 160 mg/L pepton, 110 mg/L yeast extract, 30 mg/L urea, 28 mg/L K_2_HPO_4_, 7 mg/L NaCl, 3.44 mg/L CaCl_2_ and 2 mg/L MgSO_4_, giving a mean DOC (dissolved organic carbon) concentration of about 100 mg/L (OECD, 2001). Therefore, 10 mL of the bacterial suspension were transferred to Schott flasks (control) and to beakers (plasma treatment) containing 500 mL artificial wastewater. Cultivation was performed by continuous shaking. Optical density of the culture throughout the experiment was determined at 500 nm against 0.85% NaCl solution. The plasma source was a pin-to-liquid discharge developed at the Leibniz Institute of Plasma Science and Technology (Schmidt et al., 2017). For detailed descriptions of the plasma source used for similar experiments please refer to (Hahn et al., 2024). In short, it consists of four stainless steel electrodes placed 3 mm above the liquid (Figure 1B) and operates at atmospheric pressure and ambient air. AC voltage (17 kHz) is provided by a high voltage source. Plasma treatment was applied for 15 min.

**Figure 1:**
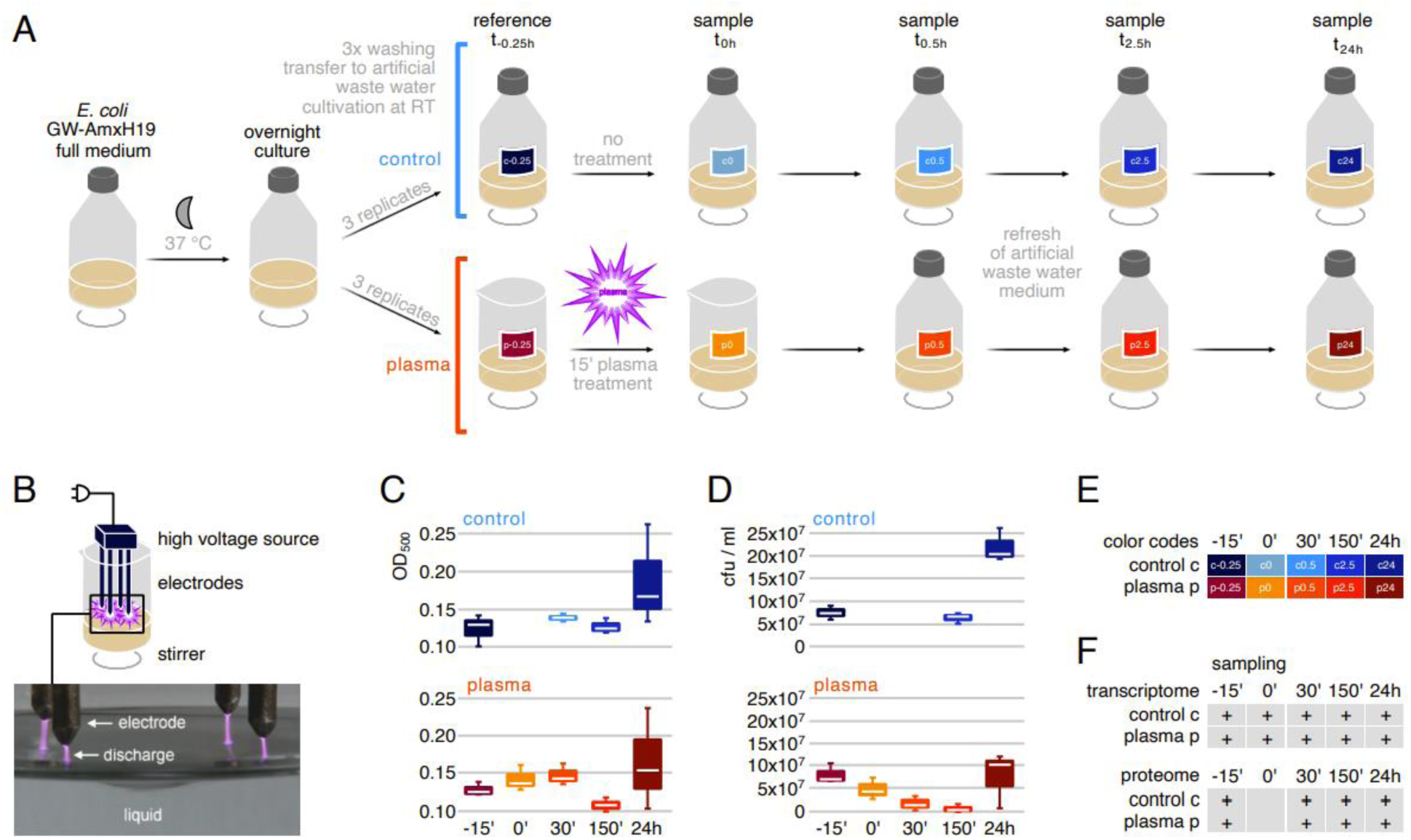
Experimental setup, sampling scheme and metadata of plasma treatment of *E. coli* GW-AmxH19. **A)** Overview of the experimental design, including sampling time points (t0.25 h, t0 h, t0.5 h, t2.5 h, t24 h) and sample allocation. The transcriptome-only sample at t0 h (immediately after plasma exposure) was included to capture early transcriptional responses. The artificial wastewater medium was refreshed at t0.5 h (30 minutes post-treatment). **B)** Technical schematic of the plasma source used in this study, as described by (Schmidt et al., 2017) and photograph of plasma discharge over a stirred liquid (Schmidt et al., 2019). **C)** OD_500nm_ measured at each sampling point. **D)** Colony forming units (CFU) determined at each sampling point. **E)** Color codes representing sampling points in figures presented in manuscript. **F)** Overview of samples collected for transcriptome and proteome analyses.

Samples of control and plasma-treated *E. coli* GW-AmxH19 cultures were taken at different time points (before treatment = t_-0.25h_, directly after treatment = t_0_, 30 min after treatment = t_0.5h_, 150 min after treatment = t_2.5h_, 24h after treatment = t_24h_) for further analysis. This comprised samples for transcriptome analysis (10 mL), proteome analysis (35 mL) and live cell counting (1 mL) (Figure 1).

The plasma-treated culture was transferred directly after treatment to a 1 L Schott flask and shaken together with the control sample at room temperature (100 rpm). Thirty minutes after plasma treatment, additional samples were taken from the control and plasma-treated culture. Subsequently, medium was exchanged by pelleting 200 mL of the culture, washing it twice with 50 mL of artificial wastewater and adding it to a new 1 L Schott flasks wit 150 mL artificial wastewater (Figure 1A). This step was necessary to enable re-grow of bacterial cells after CAP treatment. For comparison this medium exchange was also applied to the untreated control at the same time point. Further samples for transcriptome and proteome were taken 150 min and 24 h after plasma treatment (Figure 1F).

### 2.4 Determination of live cell counts

Dilution series were prepared from selected control and plasma-treated cultures (undiluted to 10^−3^ for plasma-treated sample and to 10^−5^ for control and 24 h after plasma treatment) and 50 μL of each dilution was applied to LB plates using a spiral plater (Eddyjet, I&L Biosystems GmbH, Germany). Plates were incubated overnight at 37 °C and analyzed the next day using a Flash & Go automated colony counter (I&L Biosystems GmbH, Germany) and Flash & Go software (version 1.1.1104.0).

### 2.5 Preparation of total RNA and transcriptome sequencing

Samples (10 mL) were centrifuged at 4,700 rpm for 5 minutes and the supernatant was discarded. Equal to the volume of the pellet (approximately 20-30 μL), RNAprotect® (Qiagen, Hilden, Germany) was added, and the samples were stored overnight at 4 °C. The next day, they were frozen at −70 °C for permanent storage.

Cells were resuspended in 800 µL RLT buffer (RNeasy Mini Kit, Qiagen, Hilden, Germany) with β-Mercaptoethanol (10 µL/mL) and lysis was performed using a laboratory ball mill. Subsequently, 400 µL RLT buffer (RNeasy Mini Kit, Qiagen, Hilden, Germany) with β-Mercaptoethanol (10 µL/mL) and 1,200 µL 96 % [v/v] ethanol were added. RNA isolation was performed with the RNeasy Mini Kit (Qiagen, Hilden, Germany) as recommended by the manufacturer, but instead of RW1 buffer the RWT buffer (Qiagen, Hilden, Germany) was used to isolate RNAs smaller 200 bp. To determine the RNA integrity number (RIN) the isolated RNA was run on an Agilent Bioanalyzer 2100 using an Agilent RNA 6000 Nano Kit as recommended by the manufacturer (Agilent Technologies, Waldbronn, Germany). Potential leftovers of genomic DNA were removed by TURBO DNase (Invitrogen, ThermoFisher Scientific, Paisley, United Kingdom) digestion. The Illumina Ribo-Zero plus rRNA Depletion Kit (Illumina Inc., San Diego, CA, USA) was used to remove ribosomal RNA. Strand-specific cDNA libraries were constructed with NEBNext Ultra II directional RNA library preparation kit for Illumina and the NEBNext Multiplex Oligos for Illumina (96) (New England BioLabs, Frankfurt am Main, Germany). Quality and size of the libraries were assessed with an Agilent Bioanalyzer 2100 using an Agilent High Sensitivity DNA Kit as recommended by the manufacturer (Agilent Technologies, Waldbronn, Germany). The cDNA concentration of the libraries was determined using the Qubit dsDNA HS Assay Kit as recommended by the manufacturer (Life Technologies GmbH, Darmstadt, Germany). Sequencing was performed on the NovaSeq 6000 instrument (Illumina Inc., San Diego, CA, USA) using NovaSeq 6000 SP Reagent Kit v1.5 and the NovaSeq XP 2-Lane Kit v1.5 for sequencing in the paired-end mode and running 2x 61 cycles.

### 2.6 Processing of RNA sequence data

Where possible programs and commands were parallelized with GNU parallel v20190322 (Tange, 2018). The raw sequence data was quality-filtered with fastp v0.20.0 (Chen et al., 2018). This comprised overlap correction, minimum quality threshold of 20, sequence clipping with a sliding window of 4, removal of sequences shorter 42 bp, and adapter removal. Afterwards, potential adapter leftovers were removed with cutadapt v3.2 (Martin, 2011) followed by an additional size filter with vsearch v2.15 (Rognes et al., 2016) excluding reads shorter than 42 bp. Ribosomal RNA sequences were removed with sortmerna v2.1 (Kopylova et al., 2012). Reads were then re-paired with BBMap’s v38.89 repair.sh script (BBMap, Bushnell B., sourceforge.net/projects/bbmap/). Finally, the reference genome of *E. coli* GW-AmxH19 (chromosome CP048647.1, plasmid CP048648.1) was annotated with Prokka v1.14.5 (Seemann, 2014) (including additional HMM databases: TIGRFAMs v15.0, PGAP v6.0, Pfam35, pVOG v20171012) and indexed with salmon v1.5.2 (Patro et al., 2017) (options: -k 21 -keepDuplicates) and all quality-filtered, rRNA-free sequences were mapped against it (-l A [automatically detected as ISR]--validateMappings -- seqBias --gcBias --recoverOrphans --numBootstraps 9999). All genes were additionally assigned to cluster of orthologous genes with COGclassifier v1.0.5 (Shimoyama, 2022). Nucleic acid sequences, amino acid sequences and Prokka annotation are deposited at figshare (https://doi.org/10.6084/m9.figshare.28582277).

### 2.7 Sample preparation and mass spectrometry analysis of cytosolic proteins

The samples (35 mL) were centrifuged at 8,500 rpm for 10 minutes and 4°C and the supernatant was discarded. Cell pellets were washed twice in 1 mL of TE buffer (10 mM Tris, 1 mM EDTA; pH 7.5) and finally resolved in 1 mL TE buffer. Cell suspensions were added to 500 µL glass beads (0.1 mm diameter, Satorius Stedim Biotech, Göttingen, Germany) in screw cap tubes and cells were mechanically disrupted using FastPrep (three cycles of 30 s at 6,5 m/s; MP Biomedicals, Irvine, USA). Cell debris and glass beads were removed by centrifugation (first centrifugation step 5 minutes 13,000 rpm, 4°C; second centrifugation step 30 minutes 13,000 rpm, 4°C). Protein concentration was determined using Roti-Nanoquant (Carl Roth GmbH, Karlsruhe, Germany). 50 µg of protein extract per sample were subjected to tryptic digestion using S-Trap columns (ProtiFi, Huntington, USA) according to the manufacturer’s instructions as described in detail by (Brauer et al., 2022). Purification of tryptic peptides, resolved in 300 µL of 0.1 % fluoroacetic acid, by basic pH reversed-phase fractionation was achieved using self-packed and equilibrated C18 columns (Reprosil Gold 300 C18, 5 µm; Dr. Maisch GmbH, Ammerbruch-Entringen, Germany) as described (Mücke et al., 2020). Mass spectrometry analyses were done on an EASY-nLC II coupled to an LTQ Orbitrap mass spectrometer (Thermo Fisher Scientific, Waltham, USA). Peptides were directly loaded onto a self-packed analytical column (C18 material 3.6 µm particle size, 20 cm length, inner diameter 0.1 mm) and separated by an 80 min binary gradient from 2 % buffer A (0.1 % acetic acid) to 30 % buffer B (0.1 % acetic acid in acetonitrile) and a constant flow rate of 300 nL/min. Survey scans in the orbitrap were recorded in a mass range of 300 – 2000 m/z and a resolution of 30,000. The five most intense peptide ions per scan cycle were selected for CID fragmentation in the LTQ. Single charged precursor ions as well as ions with unknown charge state were rejected from fragmentation. Dynamic exclusion for 30 s was enabled, as well as lock mass correction (lock mass 445.120025).

### 2.8 Protein identification

Raw data were searched against an *E. coli* GW-AmxH19 protein sequence database (GenBank accession numbers CP048647 (chromosome) and CP048648 (plasmid)), including reverse sequences and common laboratory contaminants (4,795 entries) using MaxQuant software (v1.6.17.0) (Tyanova et al., 2016) and the following parameters: cleavage enzyme trypsin, two missed cleavages, oxidation of methionine as variable modification, carbamidomethylation of cysteine as fixed modification. Match between runs was enabled. Unique and razor peptides were considered for label-free quantification (LFQ) with a minimum ratio count of 2.

### 2.9 Data analysis and visualization

#### Gene Functional Assignment

Functional gene assignments were performed by analyzing FASTA nucleotide and protein sequences using eggNOG-mapper (Huerta-Cepas et al., 2017), Operon Mapper (Taboada et al., 2018), and HMMScan (Finn et al., 2011). Hidden Markov model libraries from TIGRFAMs (Haft et al., 2003) and the Comprehensive Antibiotic Resistance Database (CARD) (Jia et al., 2017) were employed to assign genes and proteins to COGs, KEGG pathways, TIGRFams, antimicrobial resistance markers, and operon structures.

#### Software

Data preparation, management, analysis, and visualization were carried out partly using Microsoft Excel and predominantly with R in a Jupyter Notebook environment, utilizing a variety of packages (see below). All analyses were ultimately performed in R. For data visualization, ggplot2 was primarily employed (Wickham, 2016), supplemented by ggrepel, ggforce, ggalt, concaveman, gridExtra, ggpubr, scales, and ggnewscale to create, annotate, and arrange publication-quality plots. The stringr package was used for text manipulation, and pracma provided advanced mathematical functions. Missing values in proteomic data were imputed with imputeLCMD v2.0 (Lazar, 2015), while compositions (van den Boogaart and Tolosana-Delgado, 2008) facilitated centered log-ratio transformations. Correspondence analysis was performed with the ca package, intersections visualized using ComplexUpset (Lex et al., 2014) and eulerR, and heatmaps generated with ComplexHeatmap (Gu et al., 2016). Differential expression was assessed via DESeq2 (Love et al., 2014) for transcriptomic data and DEqMS (Zhu et al., 2020) for proteomic data, while base R functions (e.g., cmdscale from the stats package) supported multidimensional scaling.

#### Transcriptome

For transcriptome analysis, RNA counts were imported and organized into a table with genes as rows and samples as columns. Genes with very weak expression (average mRNA count < 4) were excluded prior to further processing. Normalization and differential expression analysis were conducted using DESeq2 (Love et al., 2014), supported by an experimental design table incorporating time, condition, and replicate as factors. Regularized logarithm transformation (rlog) – was applied for normalization. Sample classification was performed via ordination analysis using multidimensional scaling (MDS) based on Euclidean distances computed from rlog-transformed data. MDS was implemented using the base R function cmdscale from the stats package. Co-expression patterns among operon/regulon members were explored using correspondence analysis with the ca package, and biplots displaying gene and sample loadings— with highlighted operon/regulon members—illustrated plasma treatment–related regulation. Pairwise comparisons between plasma-treated samples and untreated controls were extracted from the DESeq2 results, taking into account mRNA expression ratios, p-values, and BH adjusted p-values, and visualized using volcano plots (per time point). Induced genes at each time point were filtered according to one of the following criteria: a p/c log₂ fold change ≥ 3; a p/c log₂ fold change > 0 coupled with an adjusted p-value < 0.01; or a p/c log₂ fold change > 2 paired with an adjusted p-value of 0.05. These genes were further analyzed and visualized using upset plots (ComplexUpset package, (Lex et al., 2014) and Euler charts (eulerR package) to depict intersections among induced gene. Finally, the ComplexHeatmap package (Gu et al., 2016) was used for gene expression profile visualization (row wise z-scores of ppm normalized data), with profiles ordered based on their temporal centroid - the weighted average of time point ranks calculated using only positive log₂ ratios as weights.

#### Proteome

For proteome data analysis, LFQ (label-free quantification) values were organized in a table with proteins as rows and samples as columns. Proteins with more than 66% missing values across samples were discarded. The remaining data were logarithmized, and missing values were imputed via the QRILC (quantile regression imputation of left-censored data) method using the imputeLCMD package v2.0 (Lazar, 2015). The imputed data were then exponentiated (delogarithmized). For normalization, either a centered log-ratio (CLR) transformation (using the compositions package, (van den Boogaart and Tolosana-Delgado, 2008) or parts-per-million (ppm) normalization was applied. Multidimensional scaling (MDS) plots were generated with the base R function cmdscale, using Euclidean distances computed from the CLR-transformed data. Further statistical analysis was carried out with DEqMS (Zhu et al., 2020), which incorporates the number of identified peptides (pep counts) per protein to refine variance estimation, thereby improving the detection of differentially abundant proteins. The log-transformed, median-centered ppm data were used as input, along with an experimental design file (treatment, time point, replicate) structured similarly to the transcriptome analysis. Pairwise comparisons were visualized in volcano plots, and overall data representation was performed using the ComplexHeatmap package (Gu et al., 2016), following the same procedures described in the transcriptome analysis section. Induced proteins at each time point were filtered according to one of the following criteria: a p/c log₂ fold change ≥ 2; a p/c log₂ fold change > 0 coupled with an adjusted p-value < 0.01; or a p/c log₂ fold change > 1 paired with an adjusted p-value of 0.05.

### 2.10 Data availability

The transcriptome raw data has been deposited at GenBank under BioProject accession number PRJNA524094. The mass spectrometry proteomics data have been deposited to the ProteomeXchange Consortium via the PRIDE (Perez-Riverol et al., 2022) partner repository with the dataset identifier PXD056785.

## 3 Results

### 3.1 Impact of plasma treatment on cell growth and viability

To analyze the effect of CAP treatment on *E. coli* GW-AmxH19 cell density and number of viable cells were determined (Figure 1C,D). Pilot experiments showed that medium exchange is necessary as otherwise no re-growth of the plasma-treated cells was possible (due to complete bacterial inactivation; data not shown). Therefore, after 30 minutes of incubation in artificial wastewater after the plasma treatment the medium was exchanged. The measurement of the optical density (OD) showed that the cell density only varied slightly between control and plasma treated samples, with higher final OD in the control samples (Figure 1C). However, in treated samples the average number of viable cells was clearly decreased by 90.16% (t_0.5h_, control vs plasma CFU, Figure 1D).

### 3.2 Transcriptomic responses to plasma treatment

Our dataset comprises 30 transcriptome samples including three biological replicates, consisting of 15 control samples and 15 plasma-treated samples derived from clonal cultures of the *E. coli* GW-AmxH19 isolate. In total, 286 million sequences (5.5–12.7 million forward and reverse reads per sample) went into analysis after quality control and ribosomal RNA filtering.

The transcriptional pattern based on the annotated genome of *E. coli* GW-AmxH19 (based on Transcripts Per Million) is depicted in a MDS plot (Figure 2A). Despite notable variation among the biological replicates, the overall timeline pattern remains highly consistent. For timepoints t_0_ – t_2.5h_ the transcriptome of plasma treated samples evolves in different pattern than that of control samples and converges to the control pattern after 24 h (t_24h_), (Figure 2A).

**Figure 2:**
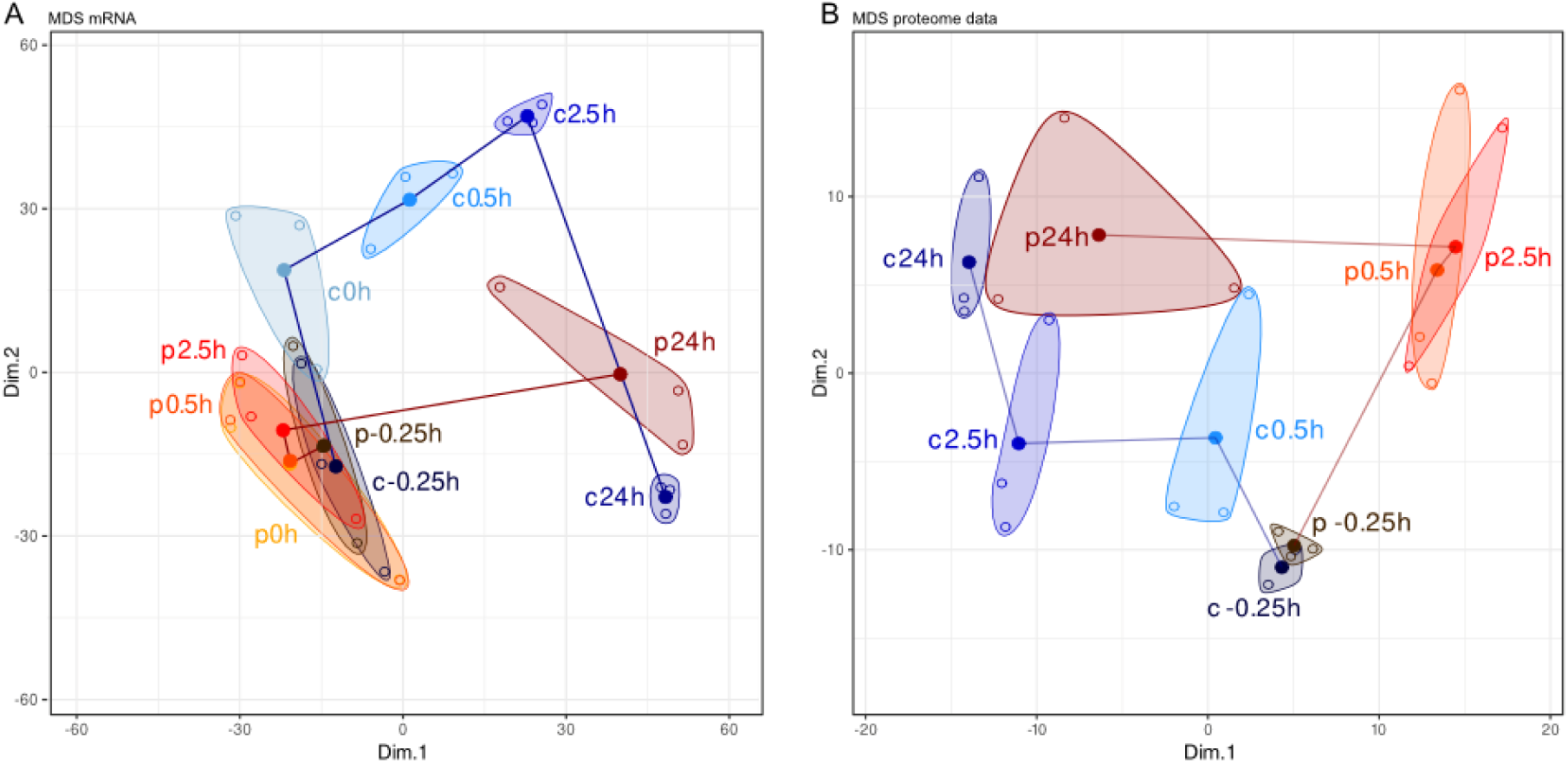
Multidimensional scaling (MDS) plots illustrating pairwise dissimilarities between samples based on global transcriptomic **(A)** and proteomic **(B)** expression profiles in *Escherichia coli* GW-AmxH19 across experimental time points. Each point represents a biological replicate, color-coded by experimental group: plasma-treatment series (red) and untreated control series (blue). Convex hulls group replicates from identical time points within each series. Samples were collected at −0.25h (prior to plasma exposure), 0h (immediately following plasma exposure), and at 0.5h, 2.5h, and 24h thereafter. Both series were sampled identically over time; however, only the treatment series received plasma exposure immediately before the 0h time point. Sample identifiers carry a prefix denoting group affiliation: “p” for plasma-treatment series (e.g., p0.5h) and “c” for control series (e.g., c0.5h). Replicates with insufficient data were excluded from the analysis. Note: For the proteomic dataset (B), no sample was collected at time point p0h, as immediate post-treatment changes in protein levels were not expected due to delayed protein accumulation dynamics. The trajectories across time reflect progressive divergence in expression profiles between treated and control series, especially at later time points.

Time-resolved analysis of transcripts in response to plasma treatment revealed a total of 1363 genes significantly differing in their expression levels as compared to the untreated control (Supplementary Table S4, Supplementary Figure S2). The response can be clustered into an early response (t_0_, and overlap to t_0.5h_) comprising 161 genes, a delayed response (t_2.5h_ and overlap to t_0.5h_), comprising 772 genes, and a late response at 24 h after treatment (t_24h_; 273 genes) (Figure 3A,B).

**Figure 3.**
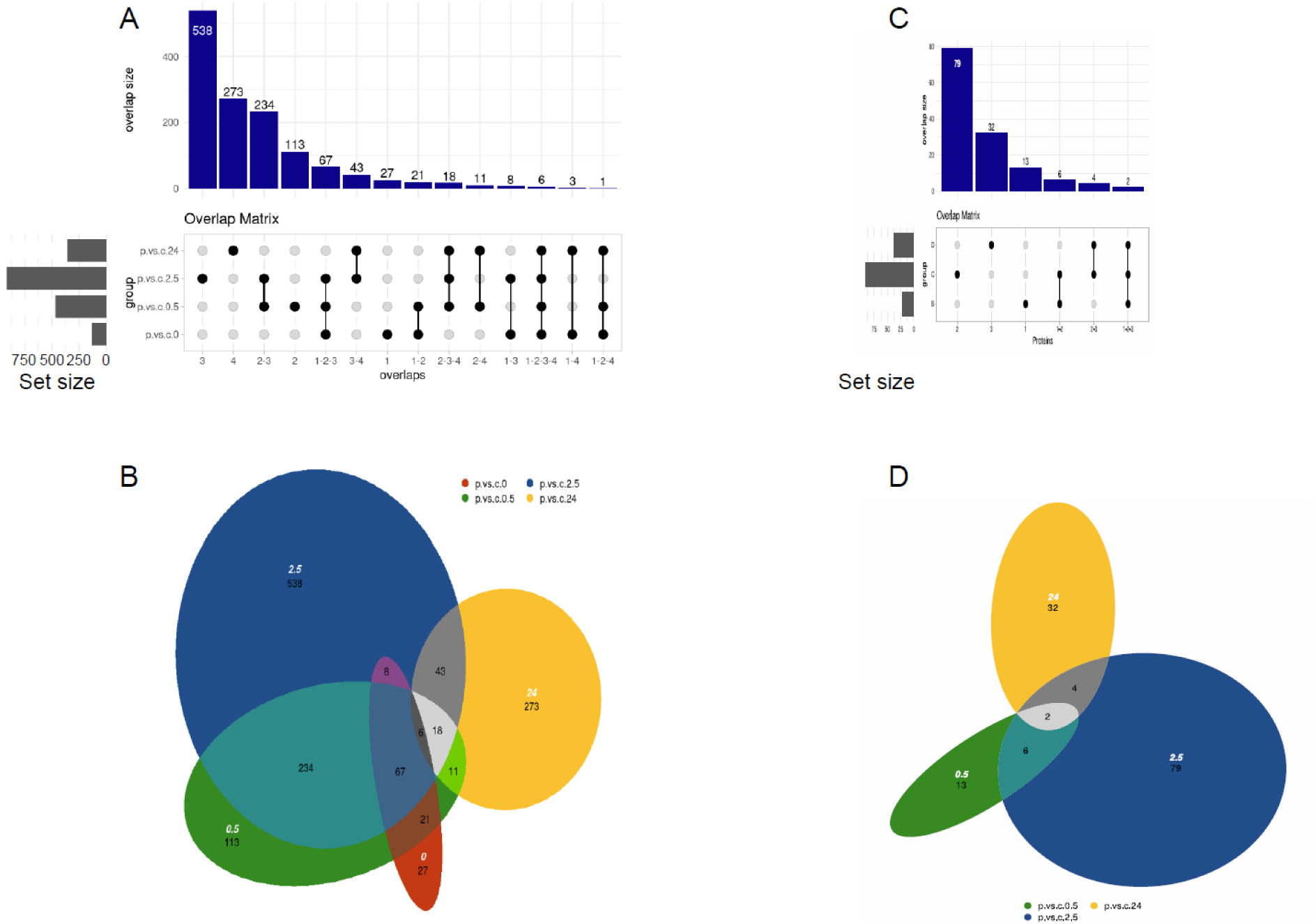
UpSet plots (**A, C**) and Euler diagrams (**B, D**) visualizing sets of genes (**A, B**) and proteins (**C, D**) with increased expression or abundance in Escherichia coli GW-AmxH19 following plasma treatment. Each horizontal bar in the UpSet plots represents the total number of elements (genes or proteins) that passed predefined filter criteria in plasma-versus-control comparisons at a given time point. Vertical bars indicate the size of intersections between time points, as specified by the connected dots in the matrix below. Euler diagrams illustrate the degree of overlap between induced elements across time points, with area sizes proportional to the number of transcripts (B) or proteins (D) meeting the respective thresholds. Filter criteria: Transcriptomic data (A, B): Strict: log₂ fold change (log₂FC) > 3 (no p-value required); Moderate: log₂FC > 2 with Benjamini–Hochberg (BH) adjusted p-value < 0.05; Statistical: BH-adjusted p-value < 0.01 (no fold-change threshold). Proteomic data (C, D): Strict: log₂FC > 2 (no p-value required); Moderate: log₂FC > 1 with SCA.padj < 0.05; Statistical: SCA.padj < 0.01 (no fold-change threshold). For transcriptomic data, p-values were corrected for multiple testing using the Benjamini–Hochberg (BH) method. For proteomic data, p-value adjustment was performed using DEqMS, which models peptide-level variance and applies BH correction; the resulting SCA.padj values thus reflect both spectral count– dependent variance and multiple testing adjustment. The term “significantly increased” is avoided here, as element inclusion was based on combined fold-change and statistical filtering criteria.

#### 3.2.1 Early response to physical plasma (time points p0h and p0.5h)

Treatment of *E. coli* GW-AmxH19 with physical plasma leads to an immediate induction of genes involved in DNA protection and (mismatch) repair, respectively (*dps and xthA, cho, uvrC, umuC_2, umuD_2, mutH*), heme biosynthesis (*hemA*, *hemF*, *hemH*), iron-storage (*bfr*) and uptake of iron and manganese (*fepA_1*, *fepD, fepE, mntH*). Genes encoding transcriptional regulators to cope with superoxide and nitric oxide (*soxS*), ferric uptake regulator *fur*, nitric oxide (*nsrR*) are rapidly induced as well. Additionally, expression of several genes encoding proteins involved in detoxification of reactive oxygen species (ROS) was induced immediately after plasma treatment (*ahpC*, *ahpF, trxC*, *katG*, *grxA, gor*, *rclA, rclC*, *azoR, yaaA* [03909]). *yhaK* gene (putatively involved in stress response and iron homeostasis) was strongly induced after plasma treatment. Furthermore, cold shock protein-encoding genes *cspA*, *cspD* and *cspG* were higher expressed immediately after plasma treatment. We also detected fast induction of expression of genes from putative type II secretion system *gsp* (*gspMEDC*). Transcription of two stress-responsive small outer membrane proteins (*bhsA_2*, *bhsA_3*) was also induced.

In contrast, plasma treatment caused a strong and immediate of the *nuo* operon encoding the components of the membrane-bound NADH oxidoreductase (*nuoB-C-E-F-G-H-I-J-K-L-M-N*). Furthermore, expression of genes and operons that encode proteins involved in biosynthesis of the enterobacterial common antigen (ECA) (genes *04671-04682*), O-antigen biosynthesis (genes *01726-01731*) and other surface structures (genes *00799-00813*) was rapidly reduced compared to the untreated control sample (Supplementary Figure S1).

#### 3.2.2 Delayed response to physical plasma (time point p2.5h)

This time point comprises the most genes with significantly induced expression compared to the control (Figure 4 and Supplementary Figure S1). Most of the highly transcribed genes from the fast response that encode proteins involved in detoxification processes or DNA repair are still expressed in high levels (e.g. *grxA*, *bhsA*_*2*, *bhsA*_*3*, *azoR*, *trxC*, *katG*, *ahpC*-*ahpF*; *umuD*, *cho*). Additionally, transcription of phage shock proteins was massively induced (*pspD-C-B_1-A*, *pspE*, *pspG*), as was the neighbouring *ompG* operon (*ycjM-02484-ycjO-ycjP-tdh_3-iolE-iolG-kojP-ycjU-ugpC_2-ompG*). *torZY*, encoding periplasmic oxidoreductase and membrane-anchored cytochrome (*torC*) were highly induced as well. An increased transcription of the cytochrome bd-II ubiquinol oxidase genes (*appB*, *appC*) was also observed. Expression of several genes encoding proteins conferring multi-drug resistances are induced at this time point, including components of efflux pumps and transporters (*emrD*, *acrF-E*, *emrA-B_1*, *acrB_1*, *mdtA-B-C-D*, *mdtK*, *mexB-mepC, mdtG*, *02848*, *yheI*-*mdlB*). Genes encoding components of a putative type II secretion system show higher expression levels (*epsCDE_1F_1F_2L*). Elevated transcription of genes involved in iron acquisition via siderophore enterobactin or other transport systems and receptors were also detected as well (*entS*, *fepD-E-G, fes_2, fes_1*, *fepA_1*, *fepA_2-entD*, *yusV-yfiZ-feuC*, *fhuE*, *hemR_1*, *hem_R_2*, *hemR_3*). Furthermore, higher mRNA levels of manganese transporter genes were observed (*hpf_2-02644-mntB_1-B_2*). Expression of a potassium ABC transport system showed also higher expression levels after plasma treatment (*kdpA-B-C-D*). The transcription of an operon encoding putative chaperone-usher fimbriae is exclusively induced at this time point (*yfcV-papC_2-yfcS_2-yfcR-yfcQ-yfcP-yfcO*). Several sulfoxide reductases showed increased transcription 2.5 h after plasma treatment (*ynfE-ynfF-dsmB_1-C_2-D*, *dsmA-B_2-C_2, msrP-Q, yedZ_1-Y_1*) just as genes encoding enzyme of the thiamine biosynthetic (*tcdA*, *thiM-D*, *thiK*, *thiI*, *thiL*, *thiC-E-F-S*). Moreover, most of the plasmid encoded genes (ranging from accessions ecoli_04756 to ecoli_04904) are highly expressed at this time point, among them genes encoding for the conjugal apparatus enabling transfer of the plasmid and antibiotic resistance genes coding for a TEM beta-lactamase (*bla*) and a tetracycline resistance gene (*tetA*).

**Figure 4.**
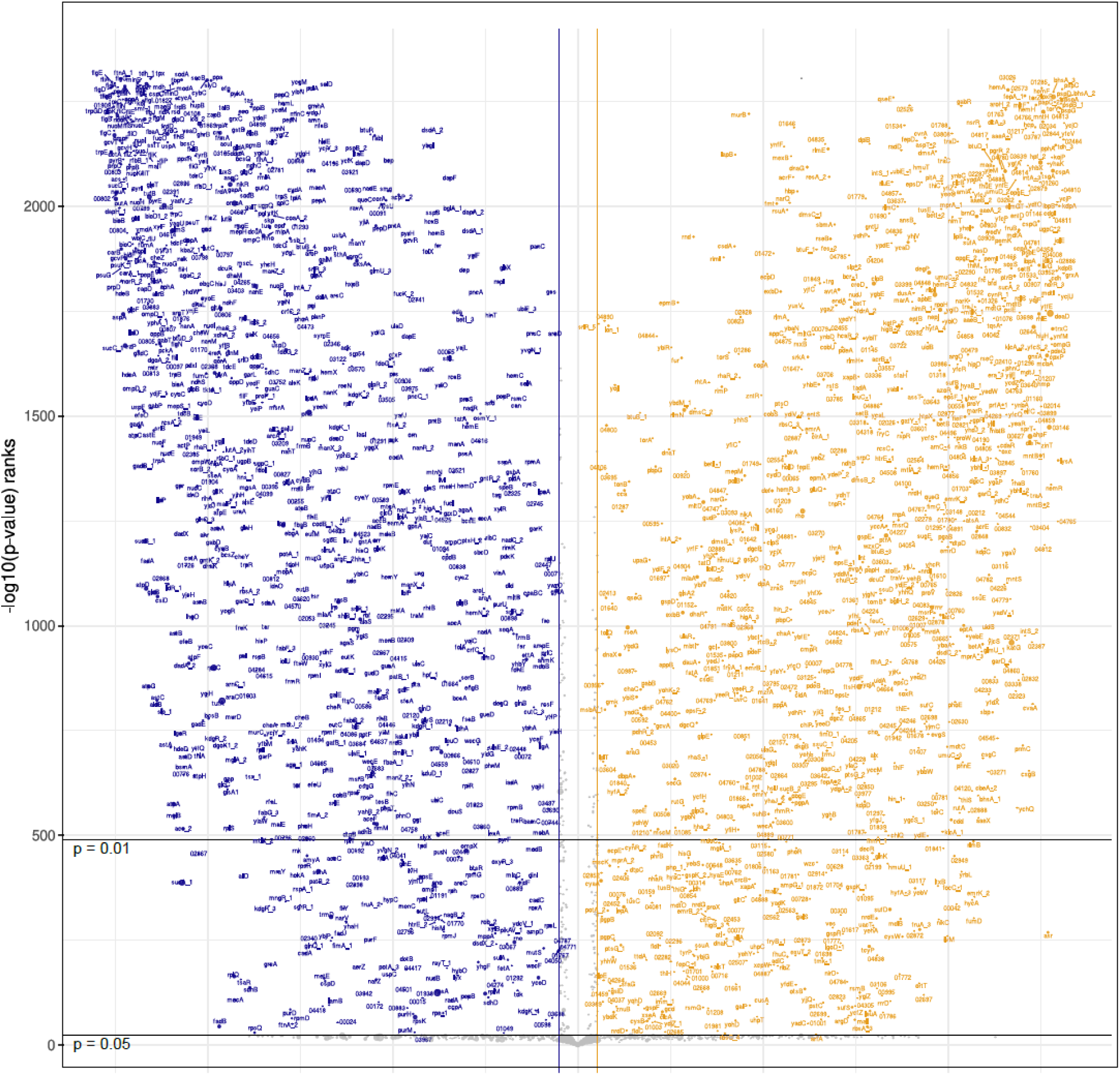
Volcano plot visualizing transcriptomic changes in *E. coli* GW-AmxH19, comparing plasma-treated and control samples at 2.5 hours post-treatment. Each point represents a gene; threshold lines indicate log₂FC = +1 (FC = 2) and log₂FC = –1 (FC = 0.5) as well as adjusted p-value cutoffs at 0.05 and 0.01. Genes with log₂FC > +1 are highlighted in orange, and genes with log₂FC < –1 in blue. This plot focuses on the time point with the strongest transcriptomic response; results for all time points are provided in Supplementary Figure S1. Note on visualization method: Axes do not reflect raw log₂FC or p-values, but instead represent sorted positions (ranks) of genes by effect size and significance. This means the original value distributions are first converted into their ranked order. To enhance readability, the ranked axis positions are then plotted using a two-zone scaling: Genes with fold change or p-values outside the defined thresholds are plotted with uniform spacing (1 axis unit per rank position). Genes within the thresholds are additionally compressed, using a scaling factor (e.g., 1 axis unit per 20 ranks). This hybrid visualization compresses the dense central region of the plot while retaining full resolution for genes of biological interest, allowing comprehensive labeling and interpretation of all relevant features — even in datasets with high transcript counts.

The expression of the aforementioned NADH oxidoreductase *nuo* operon is still downregulated as well as the expression of the potential ECA and O-antigen operons. Notably, operons encoding components of the flagellar apparatus are drastically downregulated as well at this time point in comparison to the untreated control (*flgB-C-D-E-F-G-H-I-J-K-L-M*, *flgA-M-N*, *fliY-Z-A-C, fliD-S-T, fliE, fliF-G-H-I-J-K-01909-fliM-N-O-P-01904-01903*).

#### 3.2.3 Late response to physical plasma (time point p24h)

24 hours post plasma treatment, the genes encoding phage shock proteins (*pspD-C-B_1-A, pspE*) and the *ompG* operon (*ycjM-02484-ycjO-ycjP-tdh_3-iolE-iolG-kojP-ycjU-ugpC_2-ompG*) are still highly induced compared to the untreated control (Supplementary Figure S1). The expression of many genes of the colanic acid biosynthesis operon (*ecoli_01698*, *01704*, *01705*, *01706*, *01707*, *01708*, *01708*, *01710*, *01711*, *01712*, *01714*) is induced only at this time point. Notably, the expression of genes probably encoding for the synthesis of O-specific lipopolysaccharides, that were repressed in time points 0.5h and 2.5h now show higher expression levels in comparison to the control (genes *00799-00813*). Several genes encoding sugar transport systems were induced during this phase, e.g. mannose-specific phosphotransferase system (PTS) (*manX_4-Y-Z_4*) and maltose transport (*malE-F-G*, *malK-lamB*). The genes encoding ATPase subunits are higher expressed compared to the control sample (*atpBEFHAGD*) as well as the cytochrome bo_3_ oxidase operon (*cyoABCDE*). The operon encoding components of the respiratory nitrate reductase (*narZYW*) showed also increased transcription compared to the untreated control. Elevated levels of transcription were also observed for the *cadBA* operon, encoding a lysine decarboxylase and lysine-cadaverine antiporter, and the manganese exporter gene *mntP*. SOS responsive genes *recN*, *recA* and *dinI* were also transcribed at higher levels as compared to the untreated control. Expression of genes *hupA* and *hupB,* coding for histone-like proteins, that showed decreased expression rates at the earlier time points was now also highly induced in response to plasma treatment. Remarkably, the genes encoding two complete prophages from the Siphoviridae family (encoded in region ecoli_02649 to ecoli_02712) and Myoviridae family (encoded in region ecoli_03090 to ecoli_03135), respectively (Supplementary Figure S2), are highly expressed compared to the control.

The *prp* operon encoding proteins involved in methylcitrate cycle revealed strong downregulation at transcriptional level after plasma treatment. Transcription of several tRNA encoding genes was downregulated as well (*e.g.* tRNA-Glu, tRNA-Ser, tRNA-Leu, tRNA-Cys, tRNA-Gln). An operon located on the plasmid possibly involved in iron uptake showed also lower transcription rates compared to the control (genes 04800 – 04793). Expression of a stress-responsive small outer membrane protein (*bhsA*_1, *bhsA*_3) was strongly repressed in response to plasma treatment. Several genes encoding regulators (*bssS*, *soxS*, *csrA*) were expressed at lower levels compared to the control (Supplementary Figure S1).

### 3.3 Regulon analysis using density ordination biplots

Density plots were created to detect regulons showing a positively correlated expression pattern with plasma treatment (Figure 5). Such positive correlation was observed for regulators BglJ, LeuO and StpA, that control genes and operons coding for transport and utilization of aromatic glycosides (BglJ: *bglB, bglF, ynbC, ynbD, ynbA, ygiZ,* gene *00558*), and other specific beta-glycosides (LeuO: *bglB*, *bglF*, *bglJ*, *yjjQ*) and StpA (*bglB, bglF, ygfA*). Various genes of the EvgA regulon, providing resistance to survive unfavorable conditions, showed higher transcription levels correlating with plasma treatment (*emrK*_2, *emrY*, *ydeQ*, *ydeP*_1, *yfd*, *yfdX*, *frc*, *oxc*, gene *01407*). Expression of regulons encoding genes to cope with oxidative stress and anerobic conditions were positively correlated with plasma treatment es well, e.g. OxyR (*grxA*, *hemH*, *sufABCDS*, *trxC*, *hcr*, *hcp*, *mntH*, *znuC*, *zinT*, genes *00212*, *03952*), NsrR (*sufABCDS*, *hcp*, *hcr*, *cdd*, *nor*_3, *ndhB*, *hyfA*_3, *hycA*, *yeaR*, *ytfE*, *pgpC*, gene *01095*), Zur (*zirT*, *rpmE*, *znuC*). We also observed plasma-dependent induction of genes belonging to regulons SlyA, involved in biofilm formation, *(gspCDEFGHLMO, rmf, pagP,* gene *03979)* and RcsB controlling capsule biosynthesis (*ynbC*, *ynbD*, *bglB*, *bglF*, *adiC*, *chuR*_2, *wzc*, *ygiZ*, *ydeO*, *ydeP*_1, genes *01701*, *02410*, *04400*, *00558*). Several genes belonging to the CsrA regulon, which probably provides acid stress resistance, showed also expression pattern that positively correlates with plasma treatment (*pgaABC*, *dgcT*, *glgS*, *nhaR*, gene *01348*). Three genes encoding nickel transport proteins, that are controlled by the nickel-responsive regulator NikR, showed also positive correlation (*nikABC*) (Supplementary Figure S3, Supplementary Table S3).

**Figure 5:**
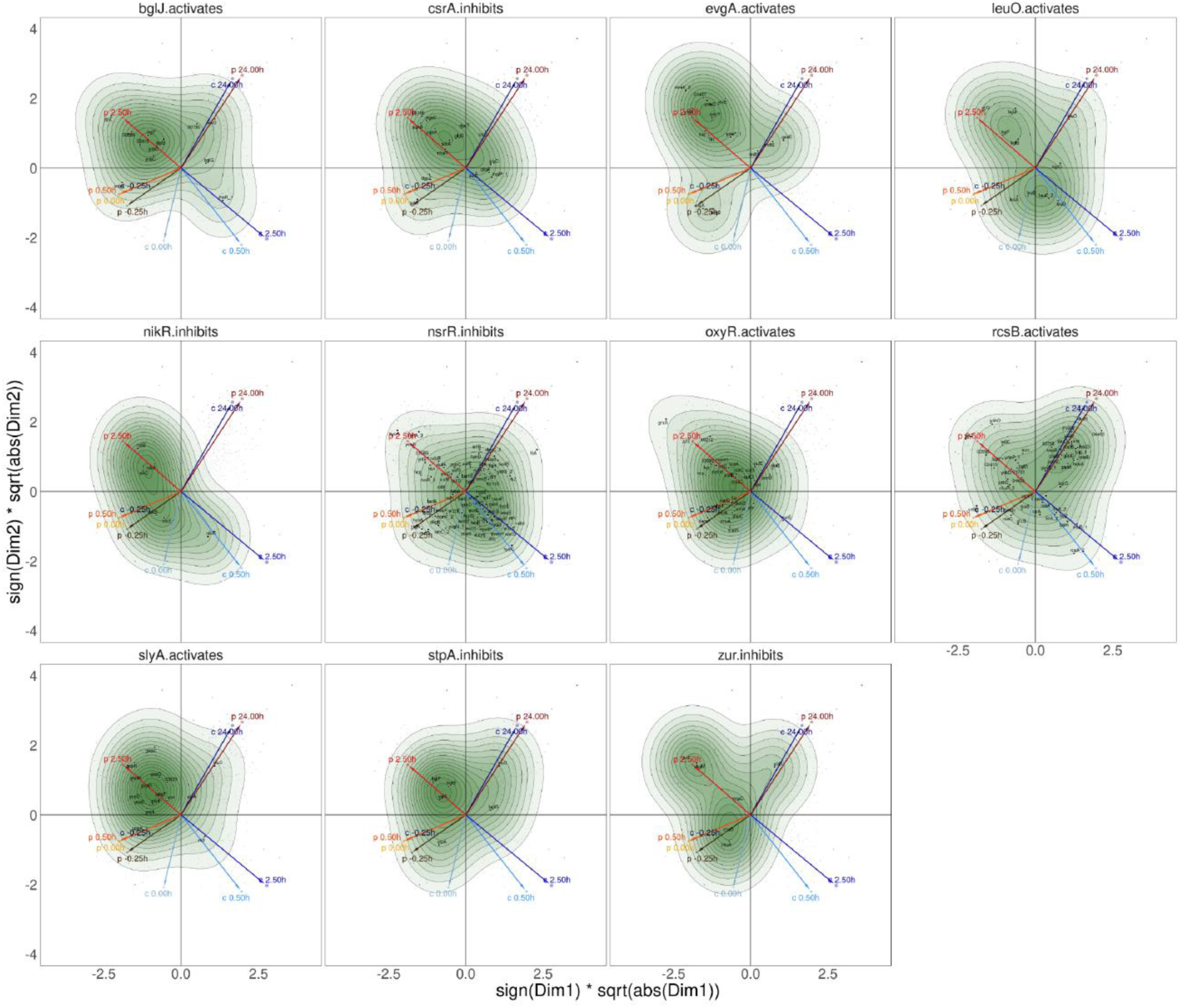
Correspondence analysis (CA) density biplots of gene expression in *Escherichia coli* GW-AmxH19 following plasma treatment. The ordination is based on the distribution of gene expression values across all samples; the first two CA dimensions (Dim1 and Dim2) capture the major structure in gene– sample relationships. All subplots show the same underlying ordination but differ in which transcriptional regulons are highlighted. Each arrow represents a biological sample from a specific time point, with red arrows indicating samples from the plasma-treated culture and blue arrows from the untreated control (see Figure 1). The red arrows at 0.5 h and 2.5 h post-treatment indicate the most pronounced transcriptomic shifts. Each point represents a gene, positioned according to its expression pattern across all samples. Labeled genes are members of the regulon of the transcriptional regulator named in the respective subplot. Green contour lines indicate the density of gene distributions in the CA space, highlighting expression pattern clusters. Only transcription factors whose regulons exhibit pronounced expression changes following plasma treatment are shown. These include activators (e.g., EvgA, LeuO, RcsB) and repressors (e.g., CsrA, NikR, StpA). Note on interpretation: In CA, distances in the plot reflect similarities in expression profiles across samples, not direct correlations or absolute expression levels. CA is based on relative abundance patterns and emphasizes co-occurrence structure, making it particularly suited to highlight sample-driven expression gradients.

### 3.4 Proteomic responses to plasma treatment

The proteome dataset comprises 24 samples including three biological replicates, composed of 12 control samples and 12 plasma-treated samples of *E. coli* GW-AmxH19. A total of 1061 proteins were identified. As already observed for the transcriptomic data the MDS plot shows a notable variation among the biological replicates, while the overall timeline pattern remains consistent. For the timepoints t_0.5_ and t_2.5h_ the proteome of the plasma treated samples evolves in different pattern than that of the controls and converges to the control pattern after 24 h (t_24h_) (Figure 2B). Similar to transcriptome level most changes on protein level were again observed 2.5 h after plasma treatment with 92 proteins showing significantly different levels compared to the untreated control time point (Figure 3 C,D, Supplementary Figure S5).

#### 3.4.1 Early response to physical plasma (time point p0.5h)

For proteomic analysis the sampling point immediately after plasma treatment was omitted; to allow cells to accumulate proteins in response to plasma treatment, the first sample was taken 30 min after plasma treatment. The highest induction rates at this early time point were observed for pyridoxine 5’-phosphate oxidase PdxH and NADH-quinone oxidoreductase subunit NuoI, involved in vitamin B_6_ biosynthesis and part of the respiratory chain, respectively. Pyridoxine 5’-phosphate synthase PdxJ showed weaker induction. Subunit NuoC was also induced, albeit to a lesser extent. Subunit O of the formate dehydrogenase complex (FdoG_2) was also present in higher amounts. Proteins possibly involved in protection against and repair of oxidative damage are found in higher amounts after plasma treatment, e.g. universal stress protein UspD, thioredoxin TrxA and thioredoxin reductase TrxB, methionine-R-sulfoxide reductase MsrC, and disulfide-bond oxidoreductase YfcG. Iron-sulfur cluster containing proteins SdhB (succinate dehydrogenase subunit) and FrdB (fumarate reductase subunit) showed elevated levels as well. The membrane-bound protease FtsH and its modulator proteins HflC and HflK were present in slightly higher amounts compared to the control sample. Impact on sugar transport is indicated by elevated levels of glucose specific PTS component PtsG_1 and phosphorcarrier protein HPr (PtsH_2). Increased amounts were also observed for Exonuclease SbcB and cold shock protein CspC, both implicated in DNA metabolism. Copper transporting ATPase CopA exporting this potentially toxic metal was also detected.

Decreased abundance due to plasma treatment was detected for proteins involved in DNA metabolism: ATP-dependent RNA helicases DeaD and RhlE, as well as cold shock protein CspA. Furthermore, several proteins involved in cell wall biogenesis (*e.g.* DdlB, GlmU, MurD, MurE, MurG, MurC) and biosynthesis of lipopolysaccharides (*e.g.* RfbB_2, RfbB_1, RfbC_2, RfbC_1, LpxC, LpxD) were present in lower amounts in plasma treated cells (Supplementary Figure S4).

#### 3.4.2 Delayed response to physical plasma (time point p2.5h)

Elevated levels of cytochrome oxidases subunits (CyoAB, CydAB) and quinol monooxygenase YgiN were observed 150 min after plasma treatment (Figure 6). In addition, the NarZ component of the respiratory nitrate reductase was present in higher amounts in plasma-treated cells. Furthermore, proteins indicative for an acid stress response were induced, *e.g.* chaperones HdeA and HdeB and glutamate/gamma-aminobutyrate transporter GadC. Efflux pump components MdtE and MdtF, also part of the GAD gene cluster important for acid resistance, showed higher amounts in plasma treated cells. The cellular response to oxidative stress caused by plasma treatment and detoxification of ROS was still active, but mediated by different proteins than observed for the first time point: glutaredoxin 3 GrxC and flavodoxin/ferredoxin-NADP reductase Fpr were found at higher levels. Two flavoproteins, oxygen-insensitive nitroreductases NfsA and NfsB, were also induced in response to plasma treatment. While universal stress protein UspD was only induced in the early time point, other universal stress proteins (UspA, UspF, UspG) were found in higher amounts at this time point. Protein YqjH which enhances siderophore utilization was also present in elevated levels compared to the untreated control. HemH, encoding ferrochelatase for heme biosynthesis, was also found in higher concentrations. YqjD, a protein possibly involved in ribosome hibernation and YqjE of unknown function, encoded by the same operon, showed also increased amounts in plasma-treated cells.

**Figure 6:**
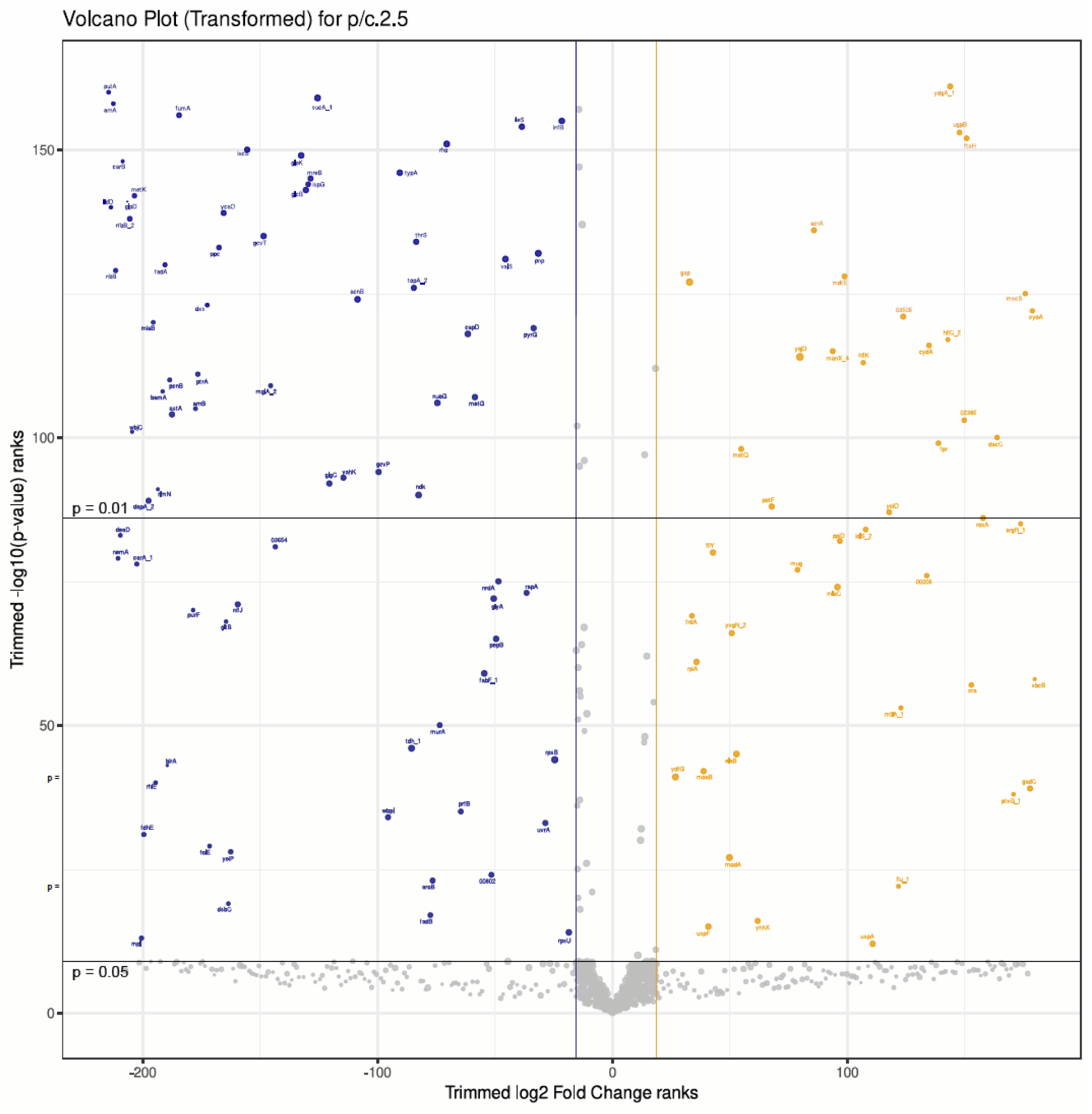
Volcano plot visualizing proteomic changes in *E. coli* GW-AmxH19, comparing plasma-treated and control samples at 2.5 hours post-treatment. The plot follows the same visualization concept as described in Figure 4, using ranked values for fold change and adjusted p-values, with compressed axis scaling within the defined thresholds. Threshold lines correspond to log₂FC = 1 (FC = 2) and log₂FC = 1 (FC = 0.5), and to adjusted p-value cutoffs of 0.05 and 0.01. Proteins with log₂FC > 1 are highlighted in orange; those with log₂FC < –1 in blue. P-values were adjusted using DEqMS with spectral count–aware modeling; adjusted values are referred to as SCA.padj.

Some proteins already present in elevated amounts in the early time point after plasma treatment, were still induced compared to the control, *e.g.* FdoG, SbcB, PtsG_1, CopA, FtsH and its modulators HflK and HflC.

Several enzymes involved in cell wall biosynthesis (*e.g.* DdlB, Mpl, MurC, MurD) and lipopolysaccharide biosynthesis (*e.g.* RfbB_2, RfbC_1, WbjC) were continuously down regulated on protein level in plasma treated cells. In addition, we observed decreased amounts for enzymes participating in biosynthesis of different cofactors, *e.g.* menaquinone/ubiquinone (UbiE_3, protein 03654), riboflavin/FAD/FMN (RibA, RibB), folic acid (FolE), others (IscA and IscS catalyzing iron-sulfur cluster biogenesis).

#### 3.4.3 Late response to physical plasma (time point p24h)

The protein with highest inductions 24 h after plasma treatment in comparison to the untreated control was the phage shock protein PspA (Supplementary Figure S4). Several proteins involved in ROS detoxification showed higher amounts in plasma-treated cells (AhpC and AhpF, KatG, GrxB, FldA, YqhD); a putative arsenate reductase YffB was also present in slightly higher levels. Furthermore, the NADH dehydrogenase Ndh was present in higher concentrations compared to the untreated control. Nitroreductase NsfB and cytochrome oxidase subunit CyoA showed still higher levels in plasma-treated cells.

Also, enzymes involved in energy metabolism and utilization of amino acids were present in higher amounts, *e.g.* AstB and AstD from arginine succinyltransferase pathway, tryptophanase TnaA, asparaginase AnsB, aspartate ammonia-lyase AspA. NADP-dependent glutamate dehydrogenase GdhA was highly induced as well. Moreover, all three key enzymes of the methylcitrate pathway metabolizing propionate (PrpC, PrpD, PrpB) were found in higher levels in plasma-treated cells. Indications for gluconeogenesis were observed by increased levels of PckA.

Compared to the earlier time point, the ATP-dependent RNA helicase DeaD is now highly induced in cells after plasma treatment. Recombination protein RecA was also found in elevated levels. The same accounts for NMN amidohydrolase PncC_1 involved in NAD recycling showing higher amounts in plasma-treated cells (Supplementary Figure S4).

Decreased amounts were still observed for some proteins involved in lipopolysaccharide biosynthesis (LpxC, MurD); RNA helicase RhlE, Rmf ribosome hibernation factor, some ribosomal proteins (RplW, RpsQ, RpsL) and translation factors (InfA, RsgA) were present in lower levels compared to the untreated control sample (Supplementary Figure S4).

## 4 Discussion

Additional treatment steps of urban wastewater could help to prevent the spread of antibiotic resistances that are released from wastewater treatment plants into the environment. Cold atmospheric plasma (CAP) with its antimicrobial efficacy and its antibiotic-degrading capacity could be such a technique implemented in wastewater treatment plants. The herein analyzed response of the wastewater isolate *E. coli* GW-AmxH19 to CAP can be divided in two stages: a fast response providing protection to and detoxification of reactive species, which was already reported in previous studies, and a long-term adaptation and recovery phase which is characterized by multiple physiological rearrangements that have not been described before. In this work, the time-resolved response of a wastewater isolate of *E. coli* was characterized in-depth. Changes in gene expression and protein synthesis are extensive and the obtained datasets demonstrate a valuable source of information in the field of antimicrobial plasma application. Only a few interesting effects could be selected and further discussed at this point.

### 4.1 First line defenses to plasma treatment

The antimicrobial effect of CAP results mainly from the formation of a variety of RONS, that cause damage to cellular macromolecules like lipids, nucleic acids, amino acids and proteins (Juan et al., 2021). Thus, immediate mechanisms are induced by *E. coli* GW-AmxH19 upon plasma-treatment to prevent and repair these damages to maintain cellular homeostasis and to detoxify reactive species. This includes expression of genes coding for ROS detoxifying enzymes such as alkylhydroperoxide reductase, catalase, glutaredoxin, and glutathione reductase (Farr and Kogoma, 1991). Reductases RclA and RclC have been shown to provide protection against reactive chlorine species (Parker et al., 2013; Sultana et al., 2022), indicating that besides RONS this reactive species might be also of relevance in plasma-treated bacteria. Interestingly, on protein level the immediate defense to the deleterious effects of CAP is mediated by different enzymes, indicating that post-transcriptional regulation in addition to transcriptional upregulation takes place to enable the cell to respond much faster to harmful conditions. Thus, the combined analysis of transcriptomic and proteomic responses to CAP enabled us to detect different levels of regulation of gene expression. This post-transcriptional regulation leads to immediate increased amounts of the universal stress protein UspD, which provides protection against oxidative stress and is involved in iron scavenging (Nachin et al., 2005). Furthermore, rapid synthesis of thioredoxin system (TrxA and TrxB) can help to reduce cysteine disulfide bonds in proteins (Anjou et al., 2024) while free methionine-sulfoxide is reduced by MsrC (Lin et al., 2007). Post-transcriptional upregulation of these enzymes might therefore be the first line of defense against oxidative damage before newly transcribed proteins accumulate and take over.

Oxidative stress results in numerous types of DNA lesions like strand breaks, base modifications or crosslink-formation (Farr and Kogoma, 1991). As a consequence, genes coding for repair mechanisms like exonucleases, the Uvr endonuclease complex or mismatch repair were found to be rapidly induced after plasma treatment. Continued replication across DNA lesions is ensured by UmuD’_2_C complex (Tang et al., 1999, 2000), while the dual role of Dps shields DNA from further damage and also protects the cell from Fenton-mediated oxidative damage (Zhao et al., 2002; Bellapadrona et al., 2010; Karas et al., 2015).

Ferrous iron is an essential cofactor in many metabolic pathways. Upon oxidation to its ferric form iron is released from iron-dependent enzymes, like mononuclear iron enzymes, iron-sulfur- or heme-containing enzymes, rendering them inactive (Imlay, 2014). In addition, ferric iron also generates free radicals that further damage lipids, nucleic acids or proteins. Manganese is much less reactive and can either substitute for iron or exclusively manganese-dependent enzymes like coproporhyrinogen III oxidase HemF are synthesized (Breckau et al., 2003). After plasma treatment we observed an expression of manganese importer MntH which helps to restore replication enzymes after oxidative stress (Wang et al., 2023). Furthermore, *E. coli* GW-AmxH19 replenishes the intracellular iron pool by rapidly expressing genes for siderophore enterobactin transport after treatment with CAP. The observed downregulation of the *nuo* genes encoding the NADH dehydrogenase as well as the increased occurrence of Nuo proteins in the cytoplasm directly after CAP exposure might be a consequence of disintegration of Fe-S clusters of this protein complex. Other Fe-S cluster containing proteins are probably inactivated as well, e.g. fumarate reductase FrdB and succinate dehydrogenase subunit SdhB, but the effect on Nuo complex might be particularly striking because of its high number of Fe-S-clusters.

An immediate downregulation of biosynthetic operons for outer membrane-located polysaccharides (o-antigen and enterobacterial common antigens (ECA)) was observed after plasma treatment. This might be a consequence of the disturbance of outer membrane (OM) integrity, as it can be permeabilized by CAP-derived RONS in synergy with acidification or lead to lipid oxidation (Xie et al., 2024). CAP has also been shown to affect the structure of the cytoplasmic membrane (Joshi et al., 2011; Xie et al., 2024).

### 4.2 Long-term adaptation to plasma treatment and recovery

Half an hour after plasma treatment cells were transferred to new medium. This medium exchange resembles a real scenario allowed to study the effects on microorganisms when plasma-generated RONS are diluted through the inflow to surface waters. Despite medium exchange after plasma treatment *E. coli* GW-AmxH19 still suffers from and adapts to the effects of different reactive species as indicated by the ongoing upregulated expression of genes encoding diverse detoxification mechanisms at timepoint 2.5 h. After 24 h the cells are already recovering from CAP-induced stress and start re-growth as indicated by higher cell counts and higher amount of several metabolic enzymes. Plasma-treated cells do not only suffer from RONS but also encounter acid stress by CAP (Hahn et al., 2024; Xie et al., 2024). Accordingly, we detected increased transcription of genes belonging to CsrA and EvgS/EvgA regulons, providing acid stress resistance (Itou et al., 2009; Gorelik et al., 2024). In line with this observation is the increased amount of chaperones and transporters encoded by the Gad gene cluster (Yamanaka et al., 2022) enabling survival at extreme acidification (reviewed in (Li et al., 2024)). Upregulation of *cadAB*, encoding a lysine decarboxylase and lysine-cadaverine antiporter, hints at milder acidic conditions in the recovery phase (Watson et al., 1992). Intracellular acidification interferes with flagellar motors (Minamino et al., 2003) and increased activity of the Gad system could lead to the observed decreased flagellar gene expression (Yamanaka et al., 2022). *E. coli* GW-AmxH19 apparently switches from planktonic to adherent state by production of adhesive fimbriae and the synthesis of colonic acid which is a major component of biofilm architecture in *E. coli* strains (Prigent-Combaret et al., 2000) and which expression is controlled by RcsB (Stout and Gottesman, 1990). Consistent with this is the positive correlation of expression of genes controlled by regulator SlyA enabling expression of different surface components (Curran et al., 2017). Therefore, the production of capsular polysaccharides and biofilm might be a delayed response after plasma treatment and a protective mechanism that acts as a physical barrier against further external stressors. In the recovery phase *E. coli* GW-AmxH19 resumes biosynthesis of O antigen, that was repressed before as a possible consequence of CAP-derived outer membrane disturbances. Nevertheless, the higher expression and presence of membrane-stabilizing phage shock proteins in later phases after treatment indicates that the bacteria still cope with envelope stress (reviewed in (Flores-Kim and Darwin, 2016)).

The adaptation to CAP was also characterized by an increased transcription of several high affinity iron transport systems, probably in a Fur-dependent manner (McHugh et al., 2003). Some of the multidrug efflux pumps, that were also induced, are involved in exporting these siderophores (Horiyama and Nishino, 2014a). Furthermore, manganese is still taken up to replace oxidized iron in enzymes; only in the recovery phase the cells start to export manganese to lower its intracellular concentration and to prevent mismetallation of iron-dependent enzymes when iron is probably available again. Additionally, we observed different effects of RONS on the expression of terminal oxidases of. *E. coli* GW-AmxH19. While transcription of cytochrome *bo*_3_ and *bd*-I oxidases was decreased after CAP treatment, *bd*-II transcription was induced at 2.5 h. Furthermore, on proteome level increased amount of Cyd (Cyt *bd*-I) and Cyo (Cyt *bo*_3_) subunits was detected which might hint at disintegration of these complexes from the membrane. While not much is known so far on the physiological role of Cyt *bd*-II in *E. coli*, available data reveal H_2_O_2_ scavenging activity (Forte et al., 2022) and a possible involvement of this complex in catalase-assisted aerobic respiration in the presence of ROS (Chanin et al., 2020). Conversely, in the recovery phase increased expression of cytochrome *bo*_3_ oxidase, which has full activity under aerobic conditions (Cotter et al., 1990), and increased expression of membrane-bound F-ATPase was detected, indicating “back to normal” conditions at time point 24 h.

### 4.3 Opportunities and challenges of CAP applications on wastewater

CAP can efficiently inactivate microorganisms – independent of any antibiotic resistances (ABR) – and also degrade nucleic acids including antibiotic resistance genes (ARG) (Daeschlein et al., 2014; Liao et al., 2018, 2019; Yang et al., 2020). Despite this high efficiency, incomplete degradation caused by shielding matrix effects and the potential for increased ARG transfer due to DNA leakage following plasma-induced membrane damage was already noted. Thus, a further optimization of plasma processes for wastewater treatments is required (Liao et al., 2018). In our study we stopped the plasma treatment after 15 min to gain an average reduction of 90 percent of living cells. However, the surviving cells showed responses on transcriptome and proteome level that could have implications with regard to antibiotic resistance development or even spread of resistances. These points should be considered and addressed in more detail in future studies.

We observed an increased transcription of antibiotic resistance genes, mainly efflux pumps and transporters, 2.5 h post CAP treatment, which could promote at least temporal increased resistance to certain antibiotics. In a previous study analyzing the short-term reaction of *E. coli* GW-AmxH19 to CAP we did not observe changed resistance profiles despite the detection of proteins potentially involved in mediating antibiotic resistance (Hahn et al., 2024). However, exposure to sublethal concentrations of hydrogen peroxide and provoking an oxidative stress response can play a role in development of ABR via increased expression of efflux pumps (Shirshikova et al., 2021) or increased mutation rates as part of the SOS response (Poole, 2012). Nevertheless, the multidrug efflux systems AcrB, AcrD and MdtABC are also involved in the export of the siderophore enterobactin (Horiyama and Nishino, 2014a). Thus, the transporters and efflux pumps transcribed in higher amounts after CAP treatment may be used for the acquisition of e.g. iron-chelating compounds to maintain iron homeostasis in the cell. In general, such functions of efflux pumps were also described for other species e.g. *Serratia marcesens*. In this case the pump is accompanied with biofilm formation (Shirshikova et al., 2021).

Our transcriptome data indicate a mobilization of the plasmid harbored by *E. coli* GW-AmxH19 since the whole conjugative apparatus is increasingly transcribed 2.5 h after CAP treatment. As this plasmid encodes tetracycline resistance and a beta lactamase (Schneider et al., 2020), mobilization and conjugative transfer could spread these resistances. In contrast to the transcriptome, the proteome data showed no higher amounts of proteins involved in these resistances (e.g. higher beta lactamase amounts) or conjugation. Conjugation is the dominant mode of HGT triggered by various factors including natural factors, contaminants from human activities, but also water treatment like chlorination and UV (Jiang et al., 2022). Particularly, oxidative stress and the SOS response caused by these agents were shown to promote plasmid conjugation (Lu et al., 2018; Wang et al., 2019). On the other hand, a transiently higher transcription of genes which are involved in conjugation may not absolutely result in a higher conjugation frequency. Thus, Li et al. (2021) described a decreased conjugative transfer frequency of ARG (of an antibiotic-resistant *E. coli*) after plasma treatment.

Our transcriptome data also revealed that treatment with physical plasma and the resulting cellular SOS response triggers activation of two prophages that are integrated in the genome of *E. coli* GW-AmxH19. Phage proteins were not detected in this study, since only cytosolic proteins were analyzed, but we could identify phage proteins in the extracellular proteome of *E. coli* GW-AmxH19 (unpublished data). The induction of prophages and the subsequent cell lysis might be a positive side effect of plasma treatment: Bacteria that are not destroyed by CAP and survive the process may eventually be killed by their lysogenic phages. The idea of using prophage activation by plasma to eradicate otherwise inaccessible bacteria was already proposed by Gu and co-workers who demonstrated that prophage activation might be a strategy to disrupt bacterial biofilms from the inside (Gu et al., 2022). However, also in this context further research is needed. Thus, HGT through imprecise excision of the phage genome is a well-known mechanism for transfer of resistance genes, also in wastewater treatment plants (Wang et al., 2025). So far, it is not quite clear if intact plasmids are released upon phage lysis, but Keen and colleagues reported lytic “superspreader” phages that promote HGT by transformation (Keen et al., 2017) thereby potentiating the risk of HGT.

Regardless which wastewater treatment procedure is used a risk for the transfer of ARG or the dissemination of primary (naturally) or secondary (acquired) antibiotic resistant microorganisms remains. This is valid for municipal wastewater treatment plants with antibiotic residues causing a selection pressure and also for methods like UV, chlorine or plasma. Moreover, risk assessments about the maximal abundance of ABR or ARG in wastewater which would bare a safety reason for humans, animals and the environment in general are missing (Rizzo et al. 2013). Thus, the comparison of different methods according to this concern is limited.

In case of plasma an insufficient inactivation can be countered through an adaption of the treatment process, e.g. a longer treatment time or a higher energy input to the plasma enabling a more comprehensive effect up to mineralization.

### 4.4 Conclusion

CAP is a promising technology to improve conventional wastewater treatment. Besides its bactericidal effect it also degrades antibiotics and other drugs. However, more research is needed to ensure a safe application of plasma processes in wastewater remediation particularly to elucidate the possible role of plasma in the transmission of ABR determinants into the environment via mobilized plasmids and phages. Additionally, consequences of a plasma treatment of wastewater for the receiving water bodies and their ecosystems should be investigated.

## Data availability

Data generated or analyzed during this study are available from the corresponding author upon reasonable request.

## Acknowledgements

We thank Melanie Heinemann for technical assistance.

This study was partly supported by the Bundesministerium für Bildung und Forschung (BMBF) in the ANTIRES2.0 project (BMBF grant 03ZZ0815A/B to KR and RD). The funders had no role in study design, data collection, and interpretation, or the decision to submit the work for publication.

J.B.: Formal analysis, Data curation, Validation, Visualization, Software, Writing – review & editing. D.S.: Formal analysis, Data curation, Validation, Software, Writing – review & editing. J.R.: Investigation. S.S.: Writing – review & editing. V.H.: Conceptualization, Supervision, Methodology, Writing – review & editing. M.B.: Methodology. A.P.: Methodology. D.A.: Methodology. J.F.K.: Resources, Writing – review & editing. K.R.: Resources, Funding acquisition. R.D.: Supervision, Resources, Funding acquisition, Writing – review & editing. D.Z.: Writing - original draft, Conceptualization, Supervision, Methodology, Validation, Formal analysis. All authors read and approved the final manuscript.

The authors declare no conflict of interest.

## Supplementary Material

Overview of genes and gene products responding to plasma treatment

**Supplementary Table 3:** Regulons and corresponding genes correlating with plasma treatment in general.

| **inductor / regulator** | **genes** |
| --- | --- |
| **aromatic glycosides** |  |
| BglJ | *bglB, bglF, ynbC, ynbD, ynbA, ygiZ, gene 00558* |
| **other beta-glycosides** |  |
| LeuO | *bglB, bglF, bglJ, yjjQ* |
| StpA | *bglB, bglF, ygfA* |
| **resistance to survive unfavorable conditions** |  |
| EvgA | *emrK_2, emrY, ydeQ, ydeP_1, yfd, yfdX, frc, oxc, gene 01407* |
| **oxidative stress and anerobic conditions** |  |
| OxyR | *grxA, hemH, sufABCDS, trxC, hcr, hcp, mntH, znuC, zinT, genes 00212, 03952* |
| NsrR | *sufABCDS, hcp, hcr, cdd, nor_3, ndhB, hyfA_3, hycA, yeaR, ytfE, pgpC, gene 01095* |
| Zur | *zirT, rpmE, znuC* |
| **biofilm formation** |  |
| SlyA | *gspCDEFGHLMO, rmf, pagP, gene 03979* |
| **capsule biosynthesis** |  |
| RcsB | *ynbC, ynbD, bglB, bglF, adiC, chuR_2, wzc, ygiZ, ydeO, ydeP_1, genes 01701, 02410, 04400, 00558* |
| **acid stress resistance** |  |
| CsrA | *pgaABC, dgcT, glgS, nhaR, gene 01348* |
| **nickel transport** |  |
| NikR | *nikABC* |

**Supplementary Table S4:**
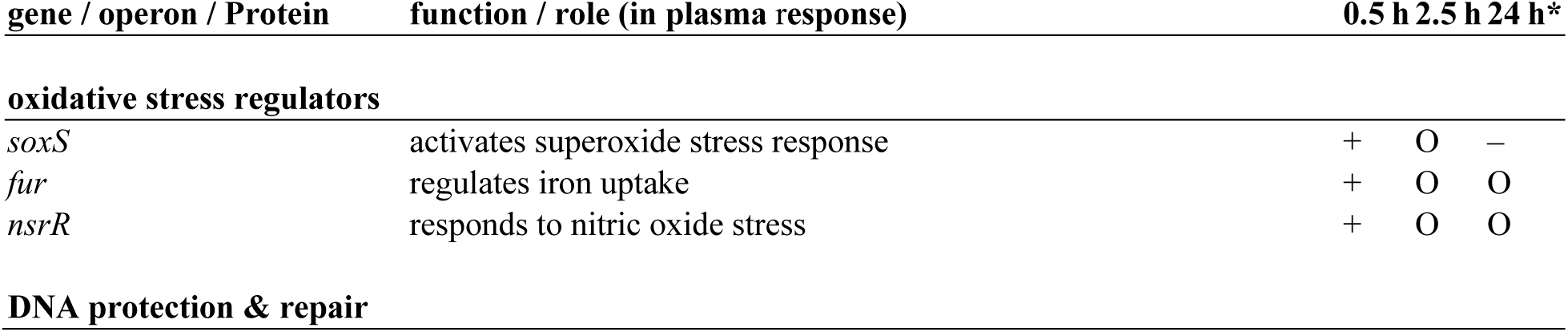
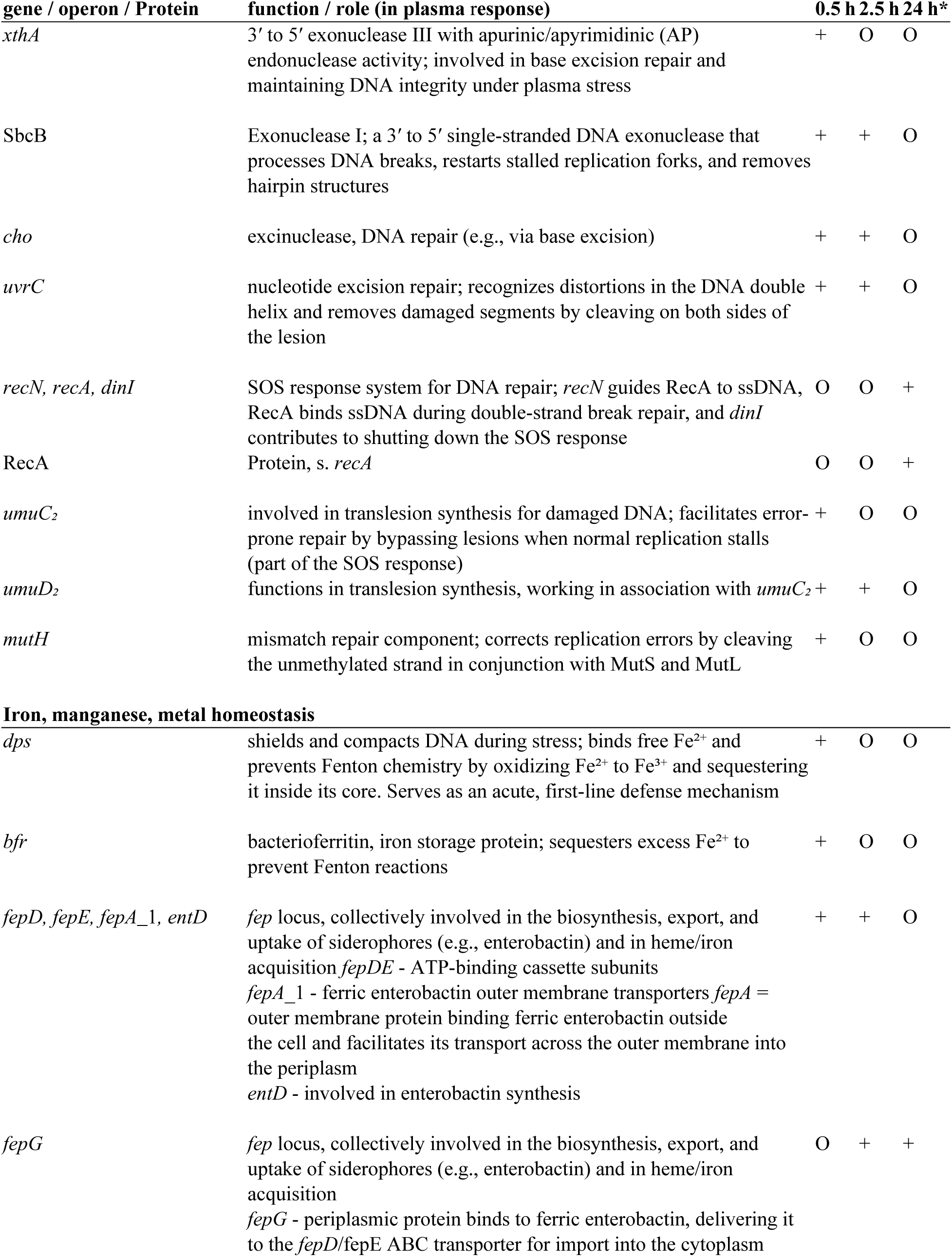
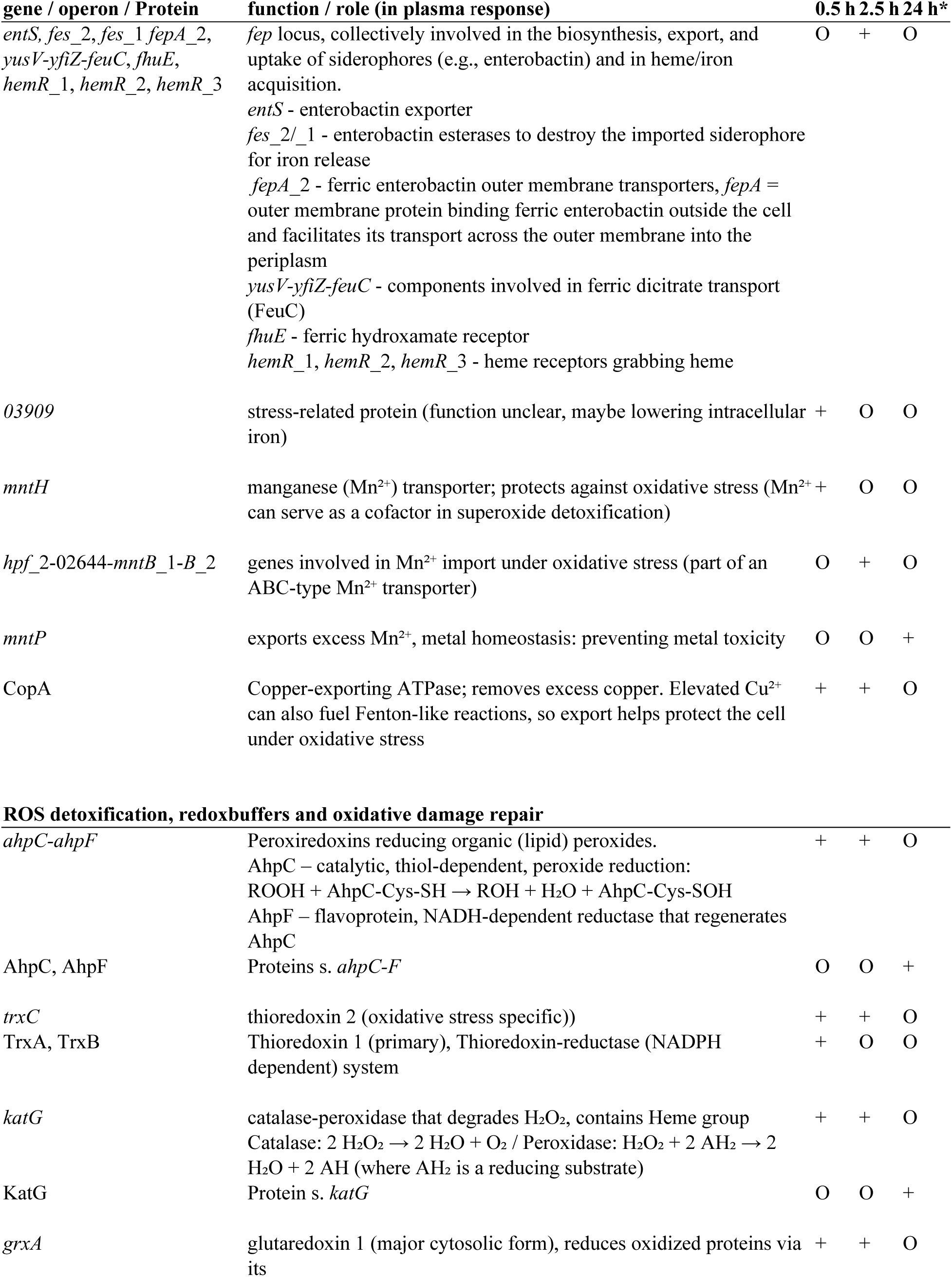
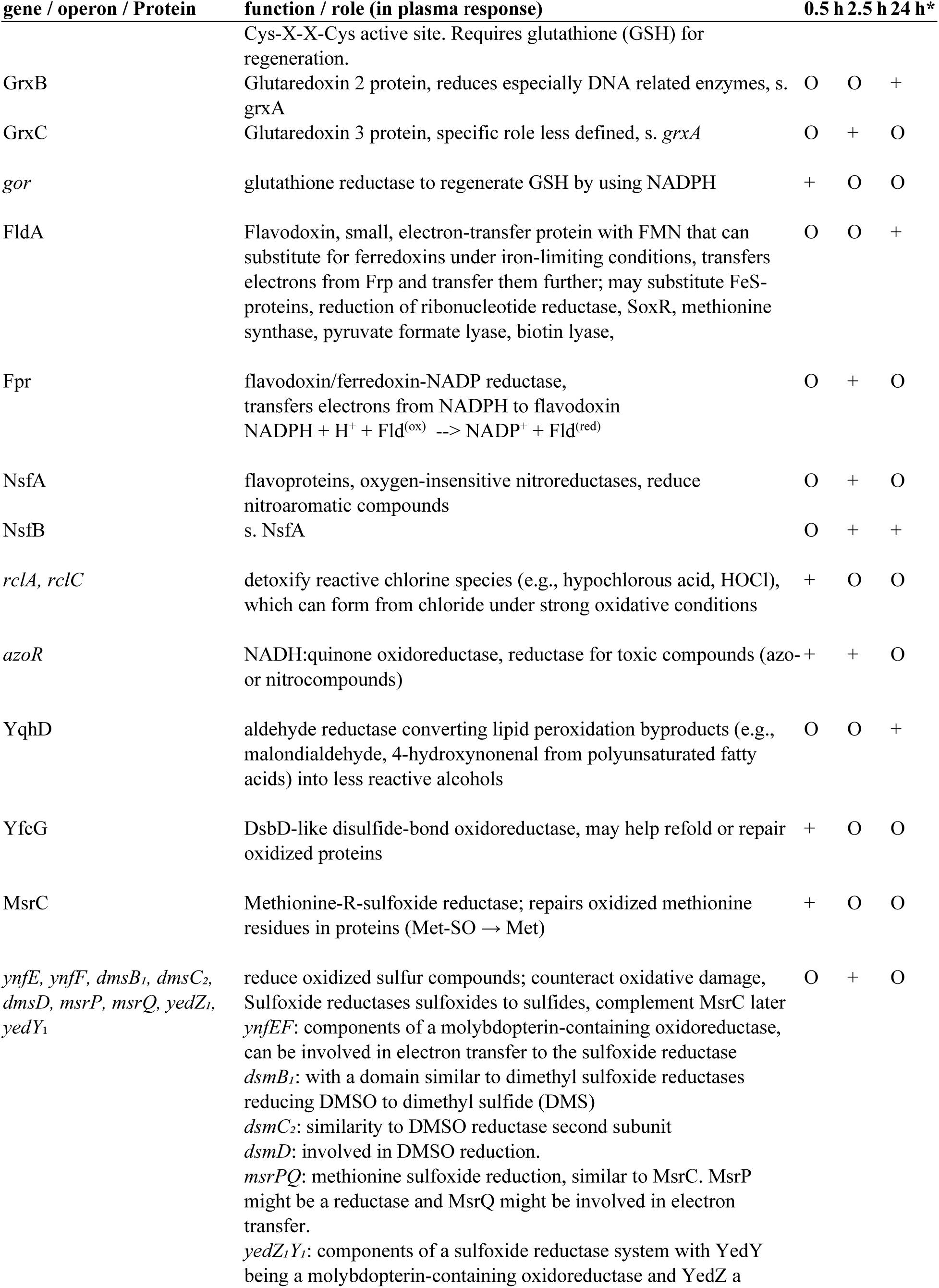
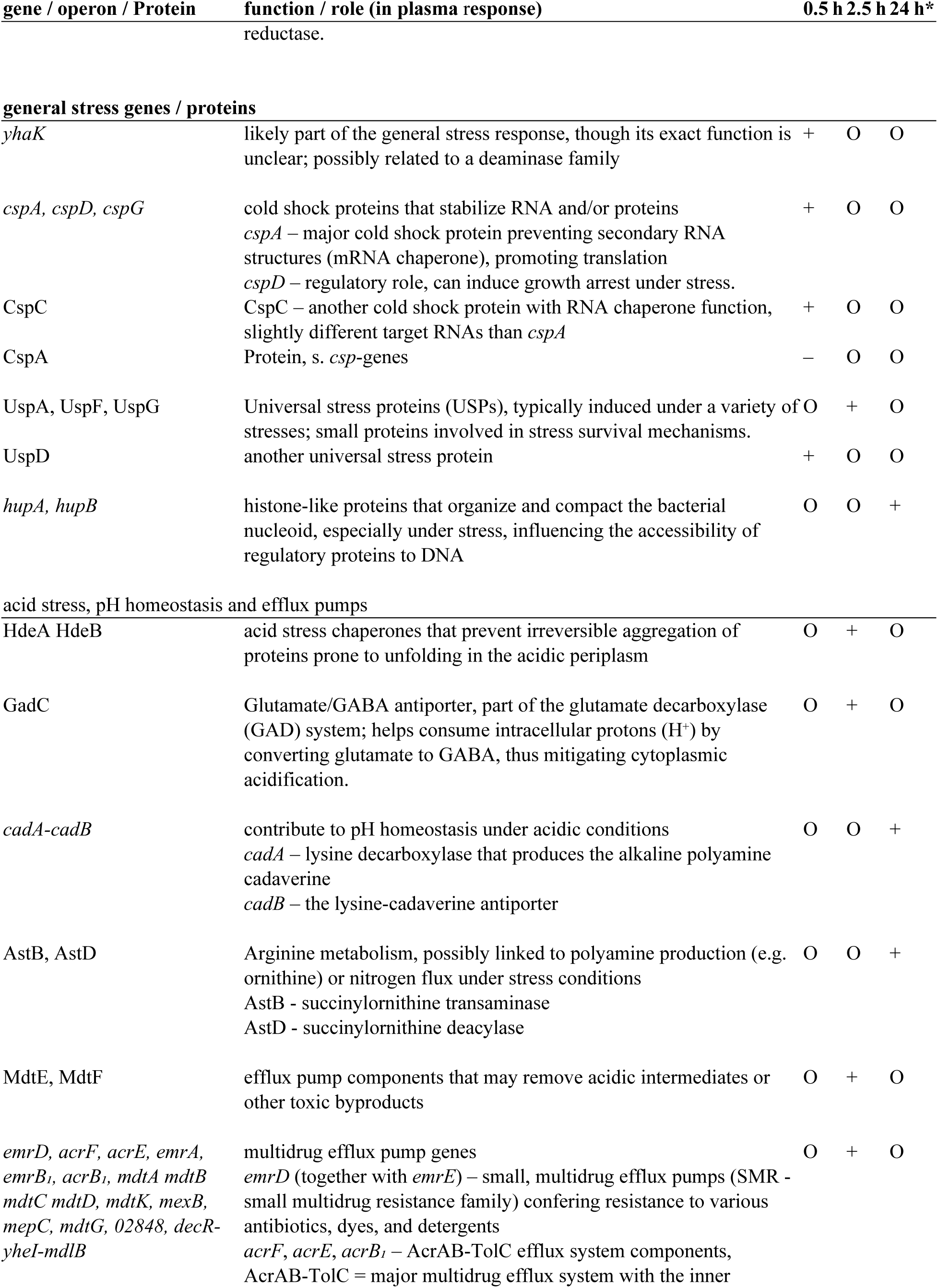
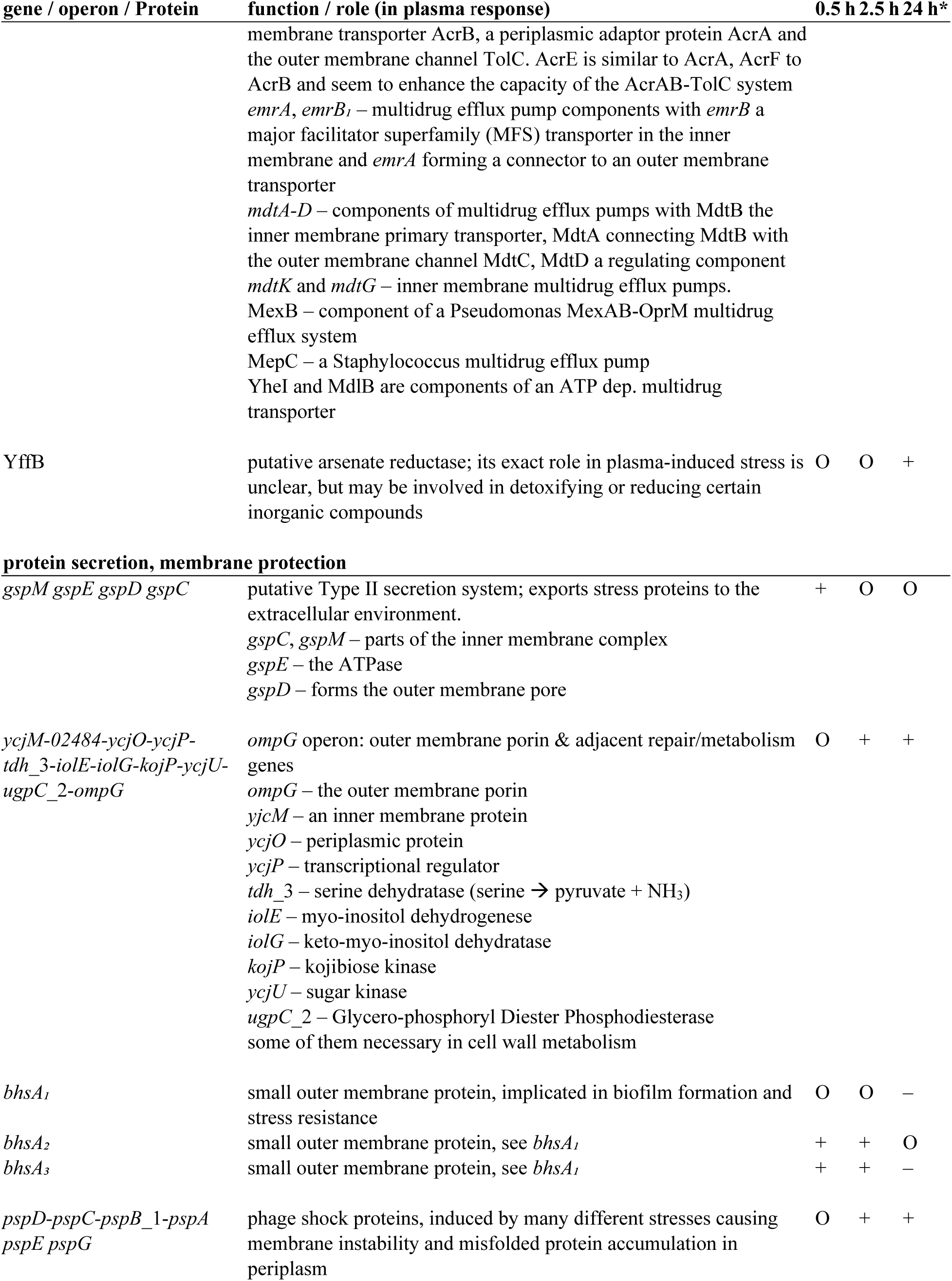
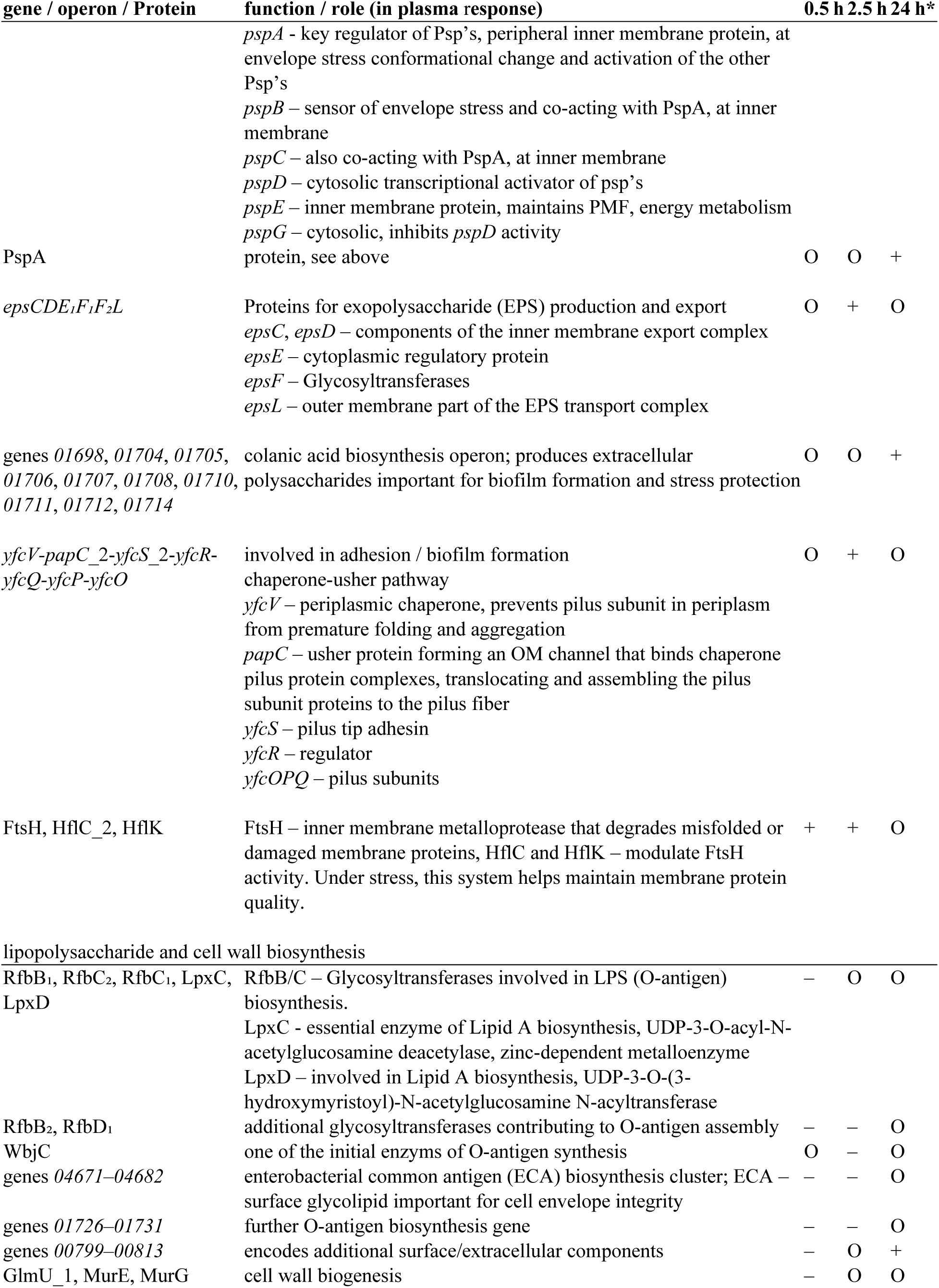
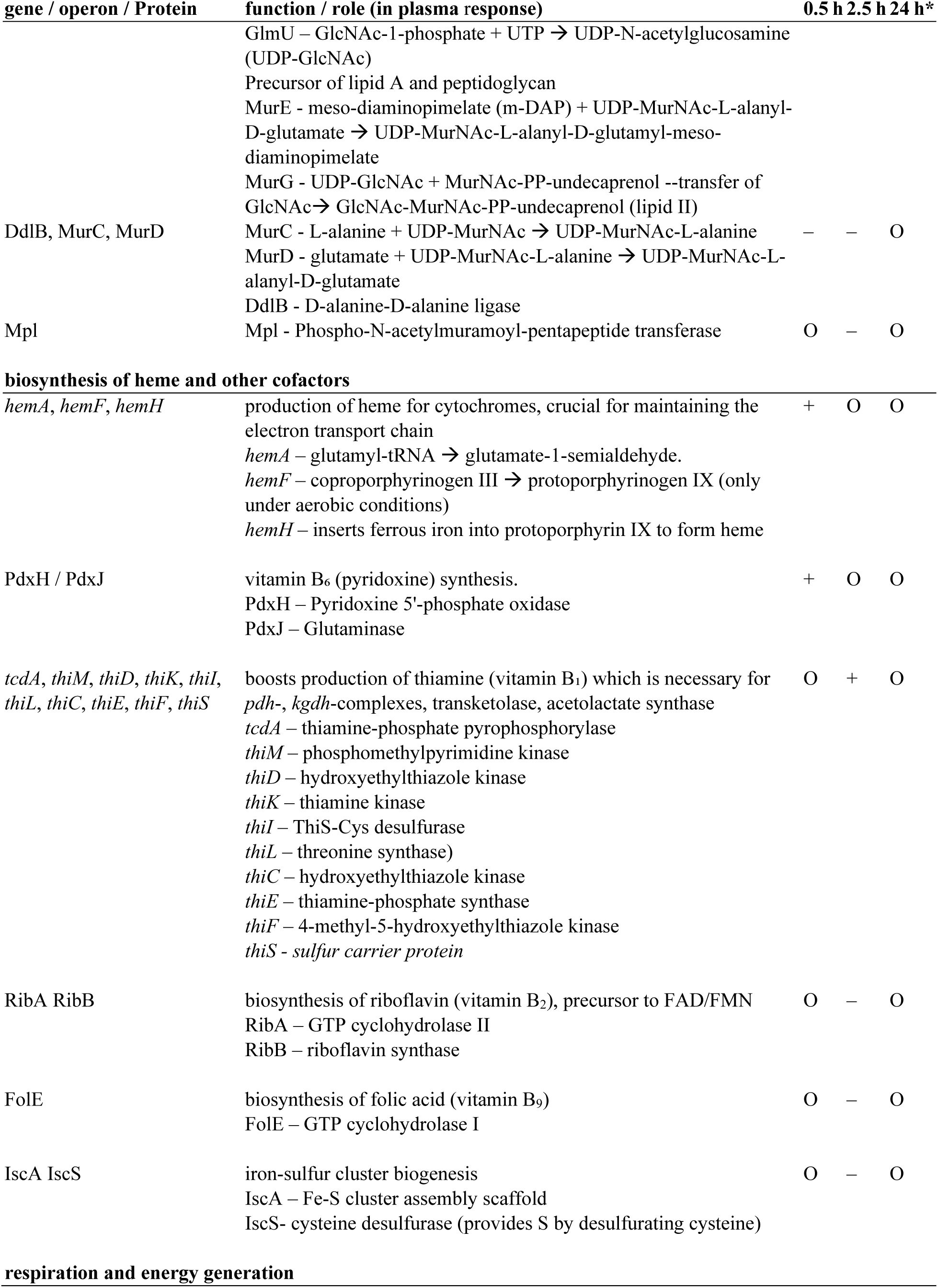
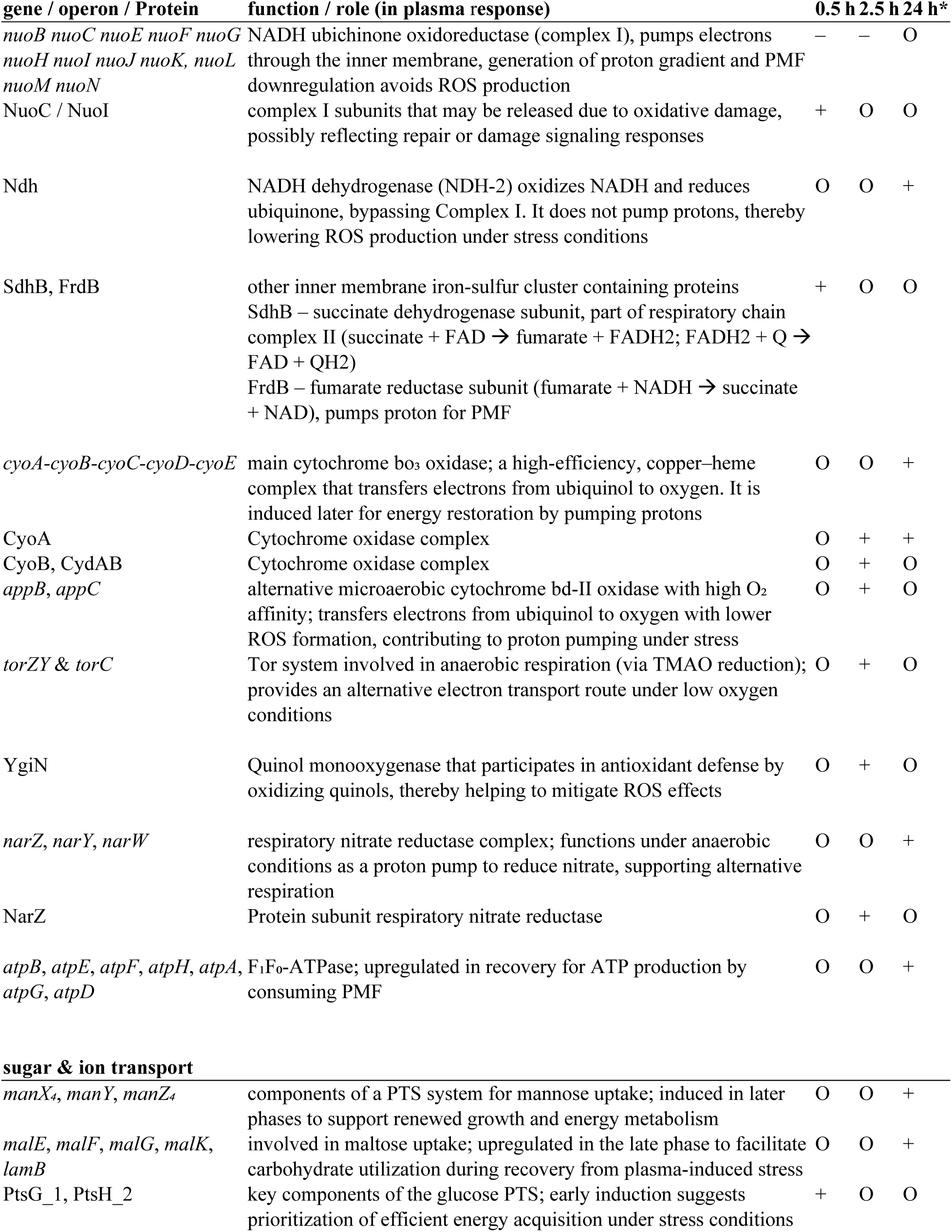
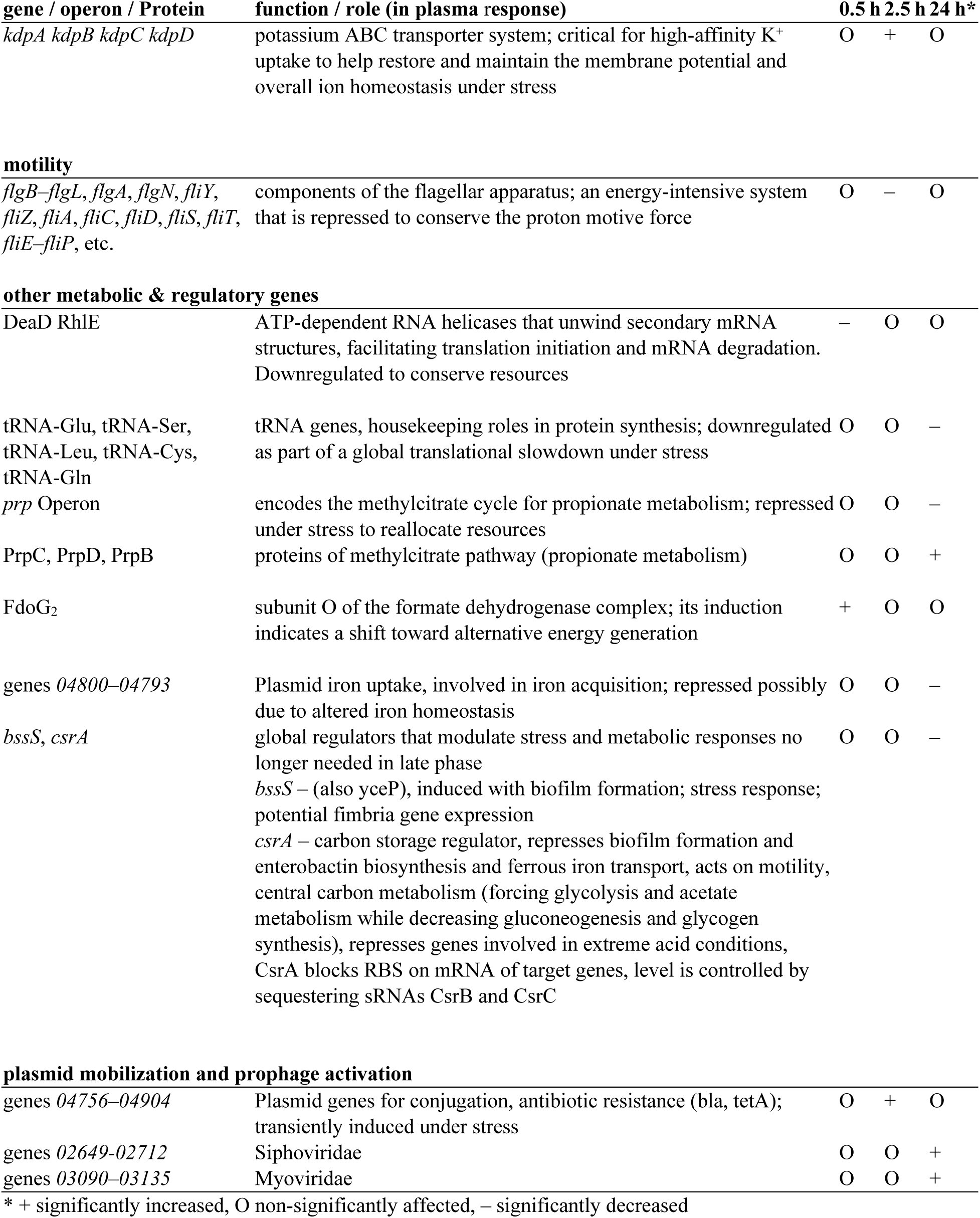
significantly behaving genes and gene products after plasma treatment.

**Figure S1: Volcano plots visualizing transcriptomic changes in *Escherichia coli* GW-AmxH19 following plasma treatment across all analyzed time points.**

Each subplot corresponds to one plasma-versus-control comparison at a specific time point, as indicated above each panel. Visualization follows the same approach as in Figure 4, using rank-based axis scaling with compressed resolution for genes within defined thresholds. Thresholds are consistent with Figure 4: log₂FC = ±1 (FC = 2 / 0.5) and adjusted p-value cutoffs at 0.05 and 0.01. Genes with log₂FC > +1 are shown in orange, and those with log₂FC < –1 in blue.

**Figure S2: Heatmap showing transcriptome-level expression profiles of all genes whose transcript or protein abundance was classified as induced at least at one time point following plasma treatment in *Escherichia coli* GW-AmxH19.**

Expression values are based on ppm-normalized RNA counts and were z-transformed per gene to highlight relative expression changes across time points and conditions.

Columns represent sampling time points, with grouped biological replicates (no gaps between replicates; visual separation between time points). Rows correspond to individual genes, annotated with gene function, gene symbol, and identifier.

Genes are grouped by functional categories, and within each category, sorted by the timing of their expression pattern in plasma-treated samples — with earlier-responding genes appearing higher in each group.

Color scale: orange indicates expression above average (positive z-score), blue indicates below-average expression (negative z-score).

The heatmap spans 7 pages to ensure readability.

**Figure S3: Heatmaps showing transcript levels of genes belonging to selected transcriptional regulons in *Escherichia coli* GW-AmxH19 following plasma treatment.**

Each row corresponds to one gene; each column to one sample. Expression values are based on ppm-normalized read abundances and were z-transformed per gene to emphasize relative expression dynamics across the time course.

Only genes that showed induction at at least one time point on either the transcriptome or proteome level (based on predefined thresholds) were included.

Regulons were selected based on coordinated expression changes in response to plasma treatment. For a visual representation of a similar subset of these genes in the context of global sample relationships, see also Figure 5.

**Figure S4: Volcano plots visualizing proteomic changes in Escherichia coli GW-AmxH19 following plasma treatment across all analyzed time points.**

Each subplot compares plasma-treated and control samples at the indicated time point. The visualization approach corresponds to Figure 6, using ranked fold change and SCA-adjusted p-values, with compressed axis scaling within defined threshold ranges. Threshold lines are set at log₂FC = +1 (FC = 2) and log₂FC = –1 (FC = 0.5), with adjusted p-value cutoffs at 0.05 and 0.01. Proteins with log₂FC > +1 are highlighted in orange; those with log₂FC < –1 in blue.

Note: The t–0.25 h time point is not shown, as no treatment had yet occurred and no proteomic differences between groups were expected. Time point t0h was not analyzed at the proteome level, as immediate post-treatment changes in protein abundance were not expected due to delayed translation and accumulation dynamics.

**Figure S5. Heatmap showing proteome-level expression profiles of all proteins whose own abundance or whose corresponding transcript levels were classified as induced at least at one time point following plasma treatment in Escherichia coli GW-AmxH19.**

Protein abundances are based on ppm-normalized values and were z-transformed per protein across all samples (including both control and plasma-treated replicates) to highlight relative expression dynamics. All biological replicates per time point are shown, except for c–0.25 hC and p–2.5 hC, which were excluded due to insufficient data/sample quality.

Proteins are grouped vertically by functional categories, and within each category, sorted by the timing of maximal expression in the plasma-treated samples — with early-responding proteins appearing higher and later-responding proteins lower in the plot.

Color scale: orange indicates above-average abundance (positive z-score), blue indicates below-average abundance (negative z-score).

**Figure S6. Predicted prophage regions in the genome of *Escherichia coli* GW-AmxH19 as identified by PHASTEST analysis.**

A) Circular genome map showing the locations of four predicted prophage regions: two classified as complete (green: Region 3 and Region 4) and two as incomplete (red: Region 1 and Region 2).

B) Detailed gene maps of each prophage region, displaying predicted open reading frames, gene orientation (arrow direction), and functional annotations. Coloring reflects the predicted completeness (green = complete, red = incomplete).

PHASTEST classifies prophages based on gene content, synteny, and presence of phage-related features such as integrases, capsid proteins, and tail fiber genes.

