## Supplementary material for "Time-resolved effects of cold atmospheric plasma on *E. coli* GW-AmxH19 transcriptome and proteome in an emulated wastewater environment": Figures and Supplementary Figures: Ecoli_plasma_allfigures-and-supplementaryfigures_biorxiv.pdf

Fig 1
Workflow

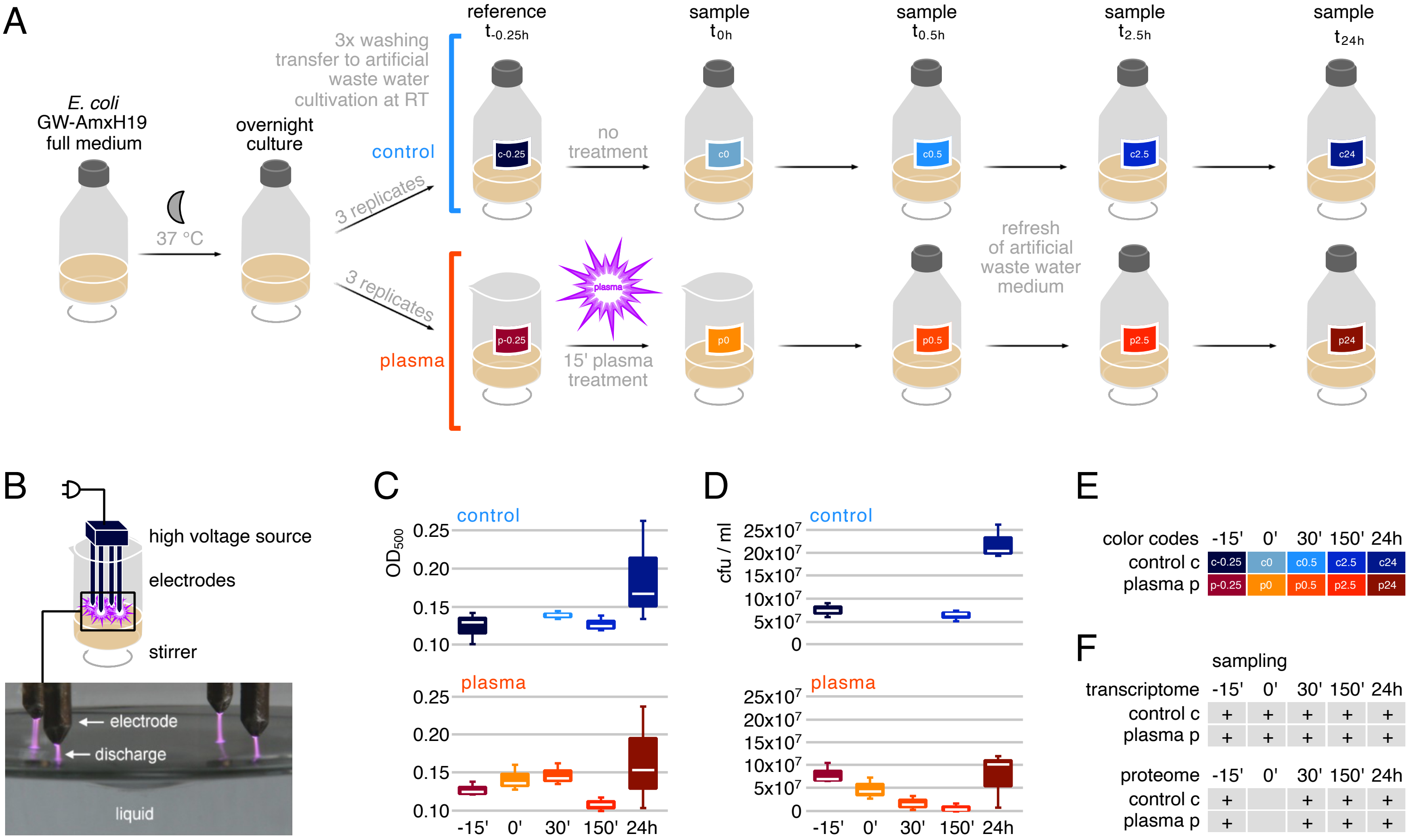

Fig 2
A MDS Plot RNASeq
B MDS Plot Protein

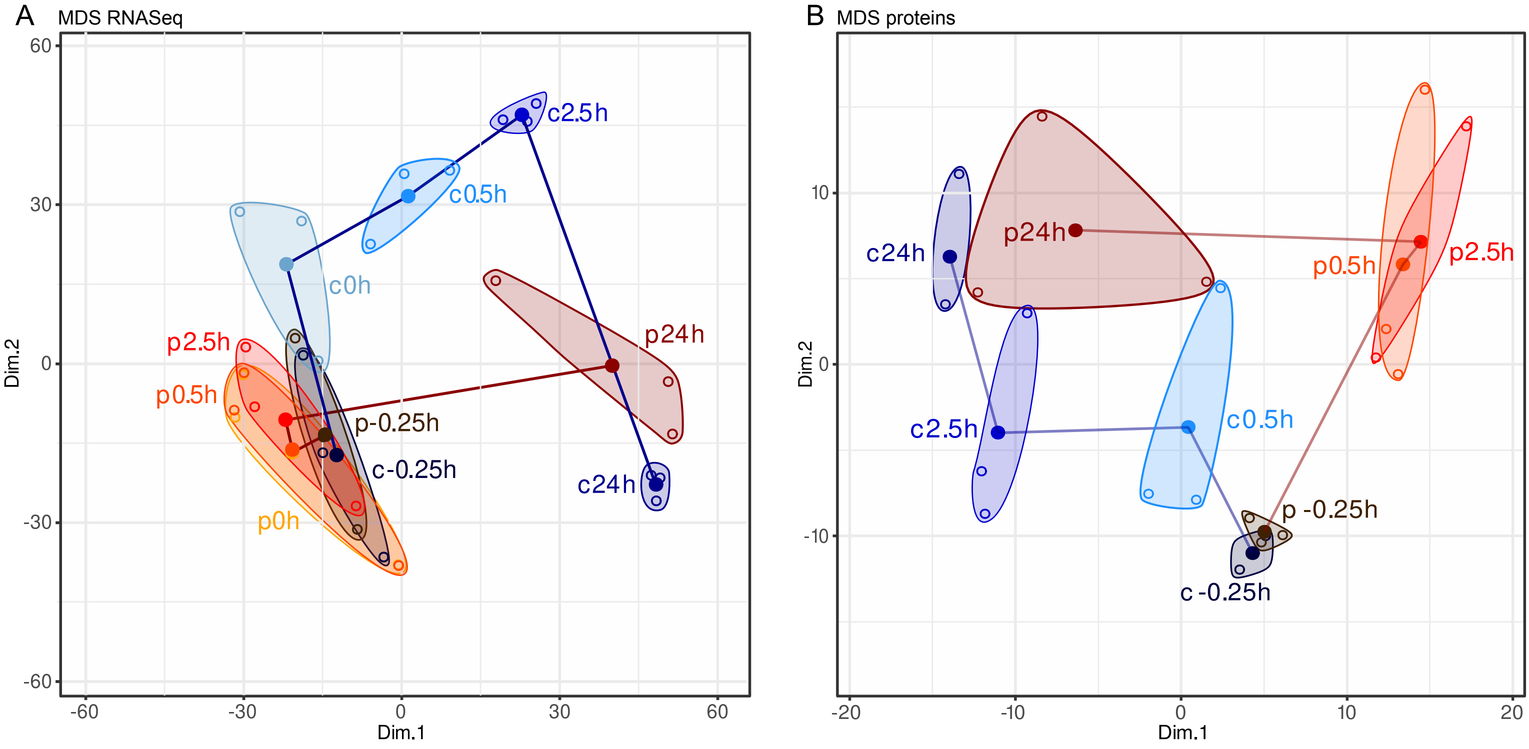

Fig 3
A Upset plot / B Euler chart RNASeq
C Upset plot / D Euler chart Proteins

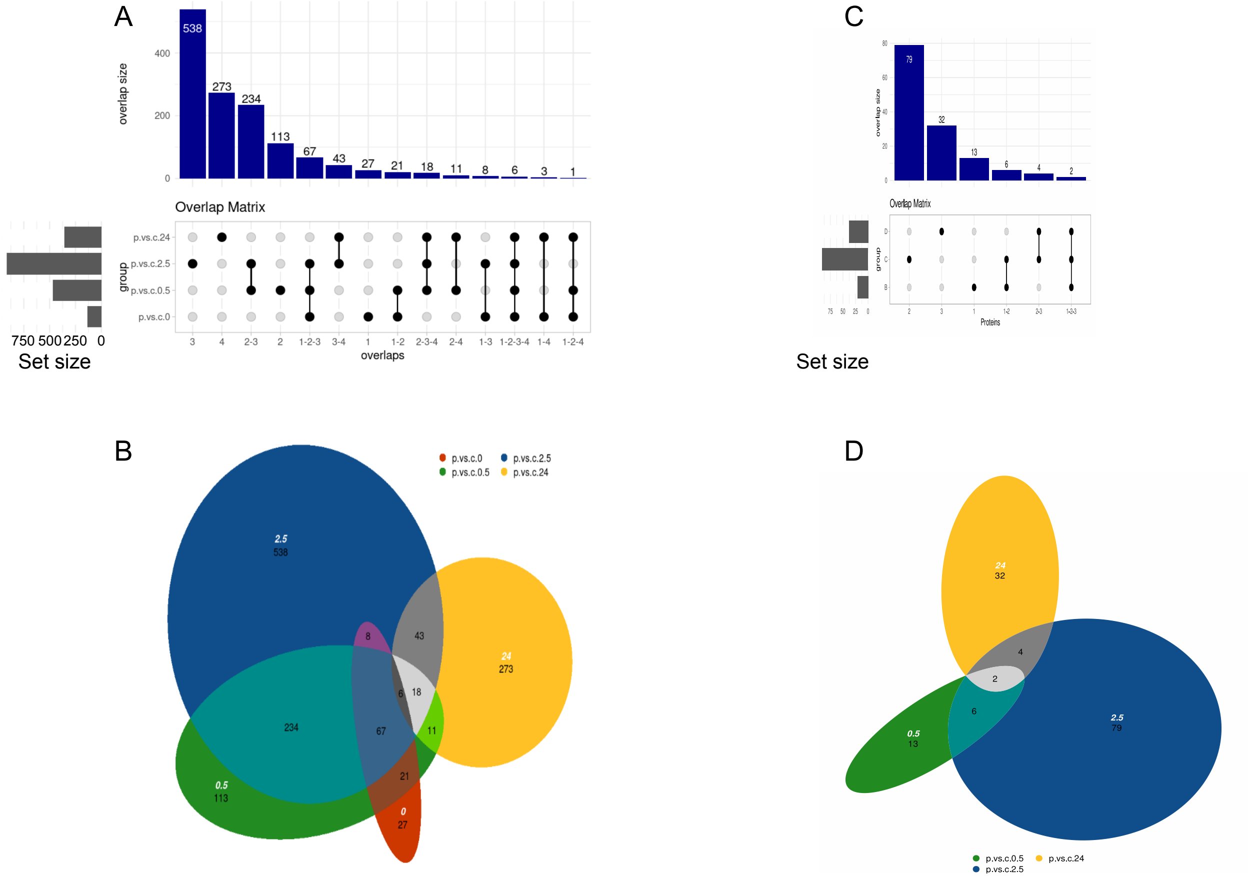

Fig 4
Volcano Plot mRNA plasma vs. control at 2.5h

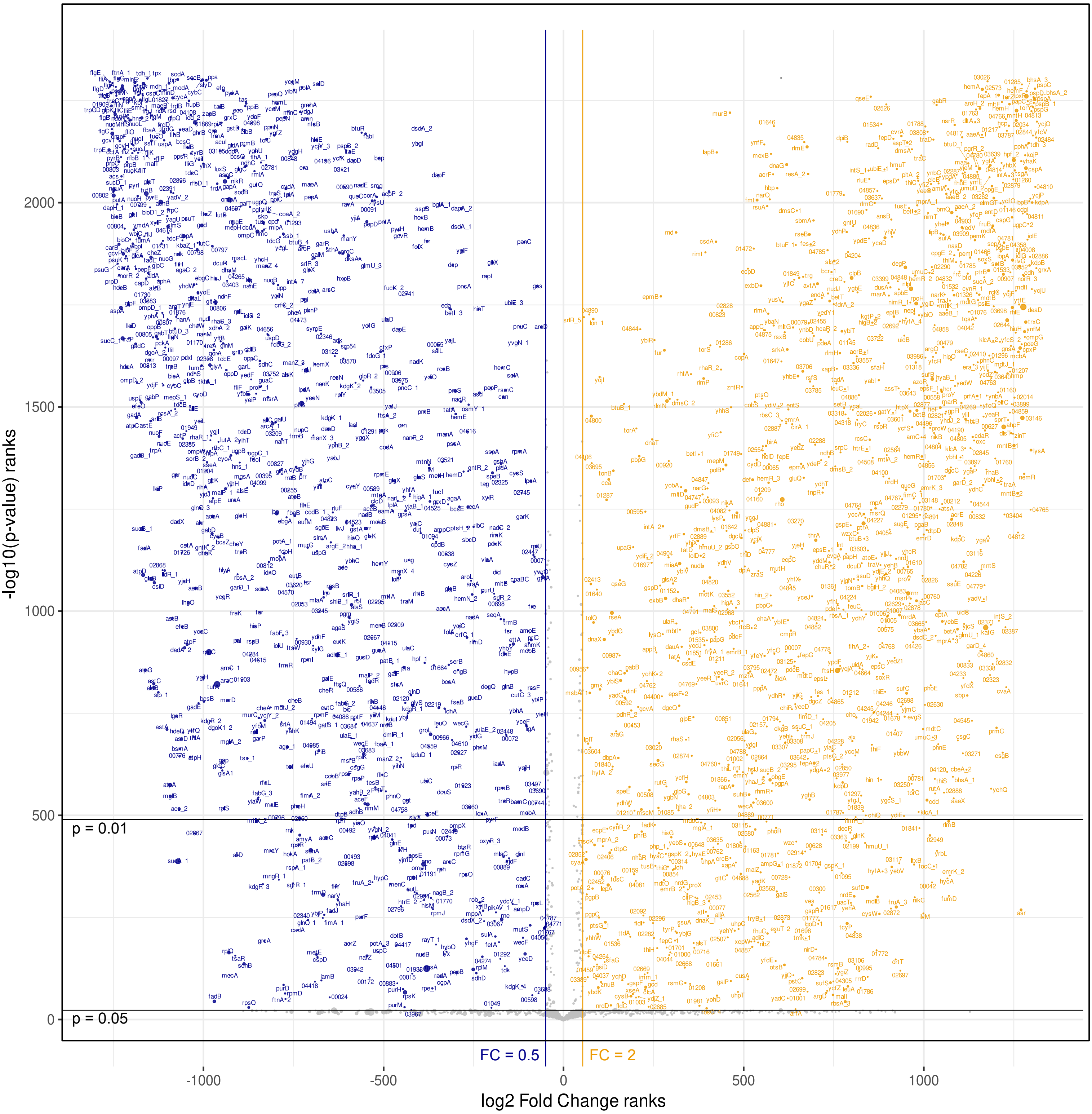

Fig 5
RNASeq density plots of regulons

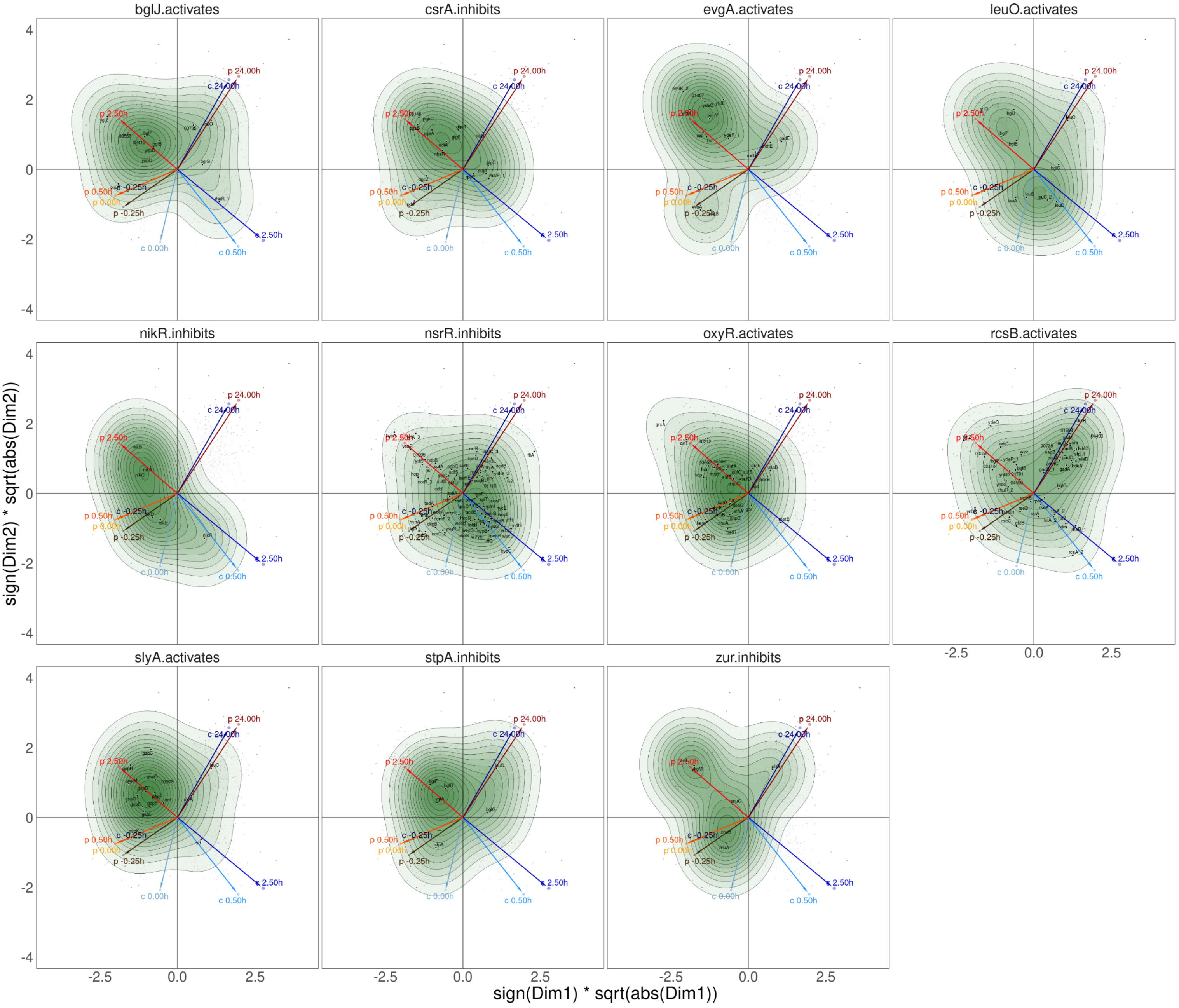

Fig 6
volcano plot proteins 2.5h

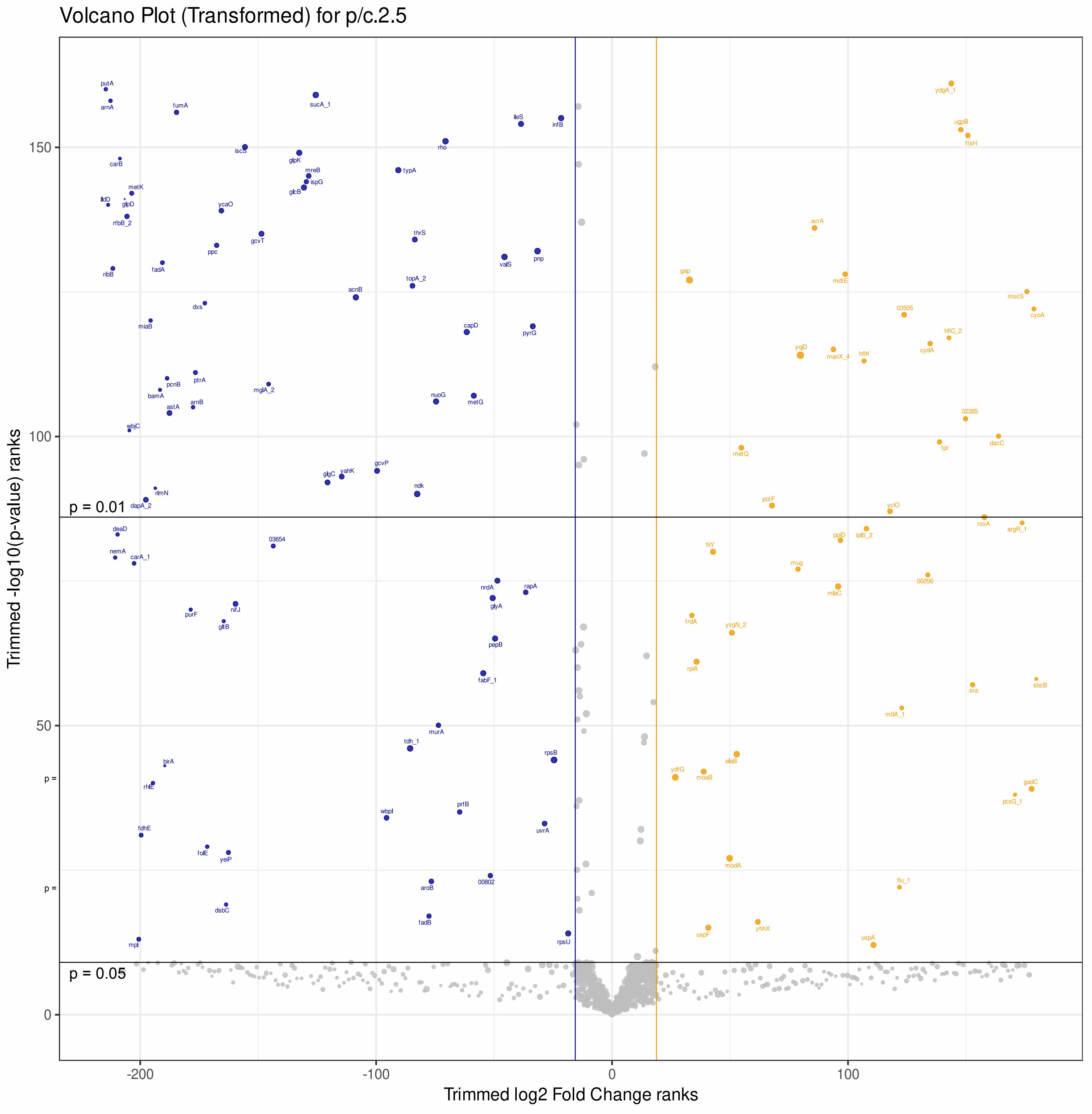

### Fig S1 - supplements Fig 4
### Volcano Plots RNASeq all p vs c panels - supp

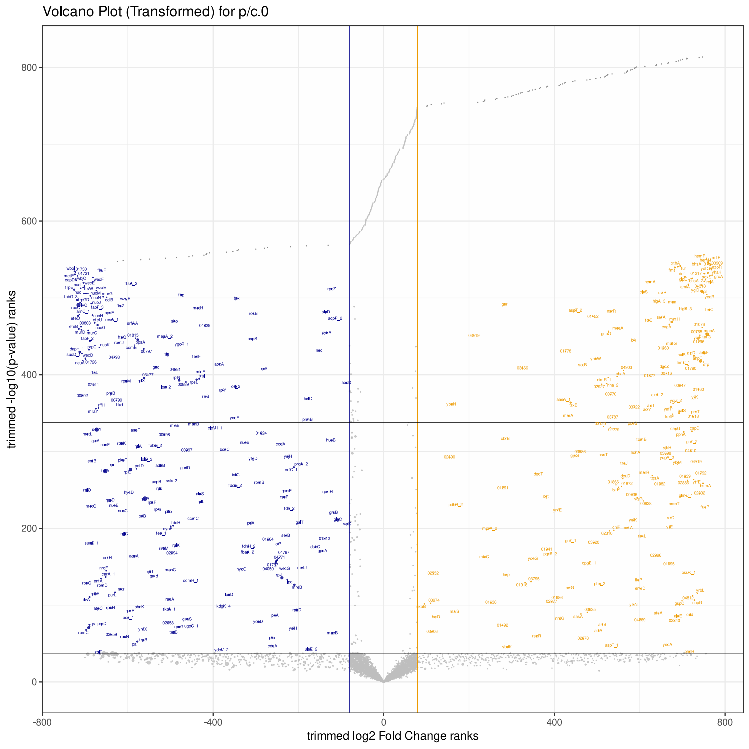


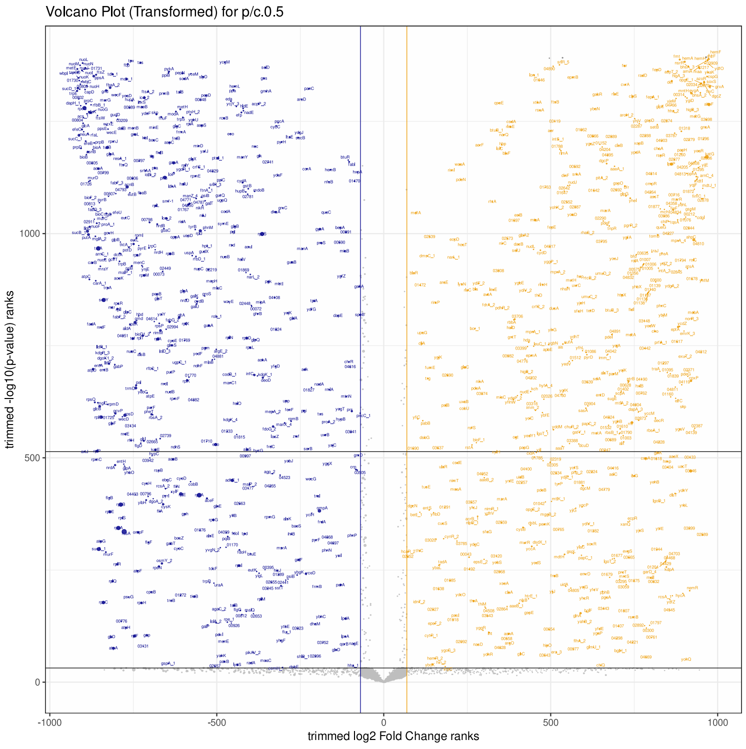


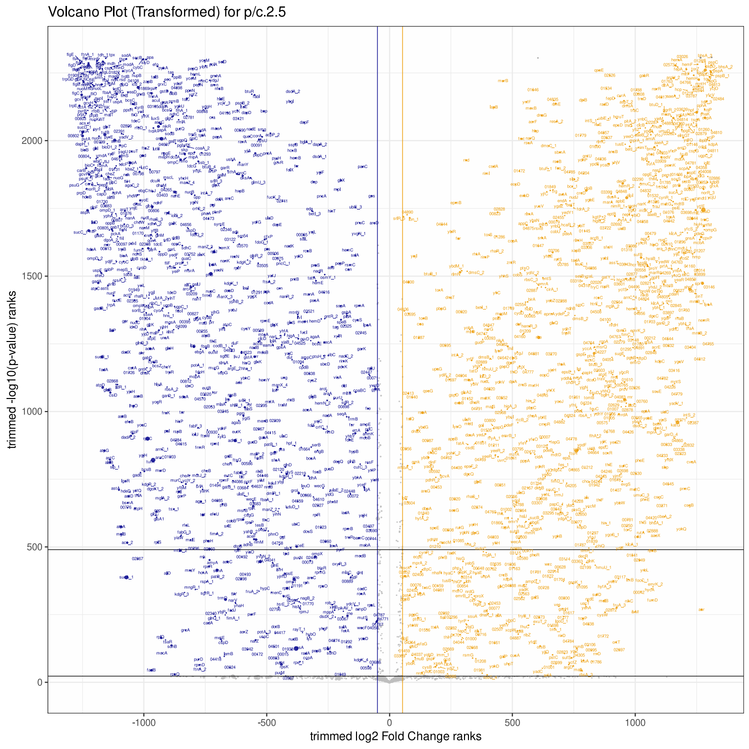


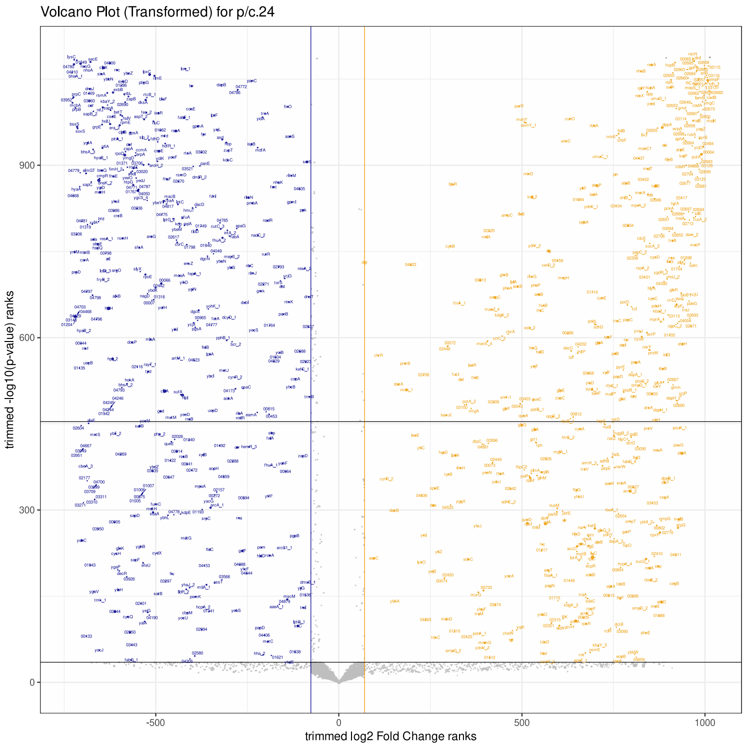

### Fig S2 - supplements RNA results
### RNASeq - induced genes - supp

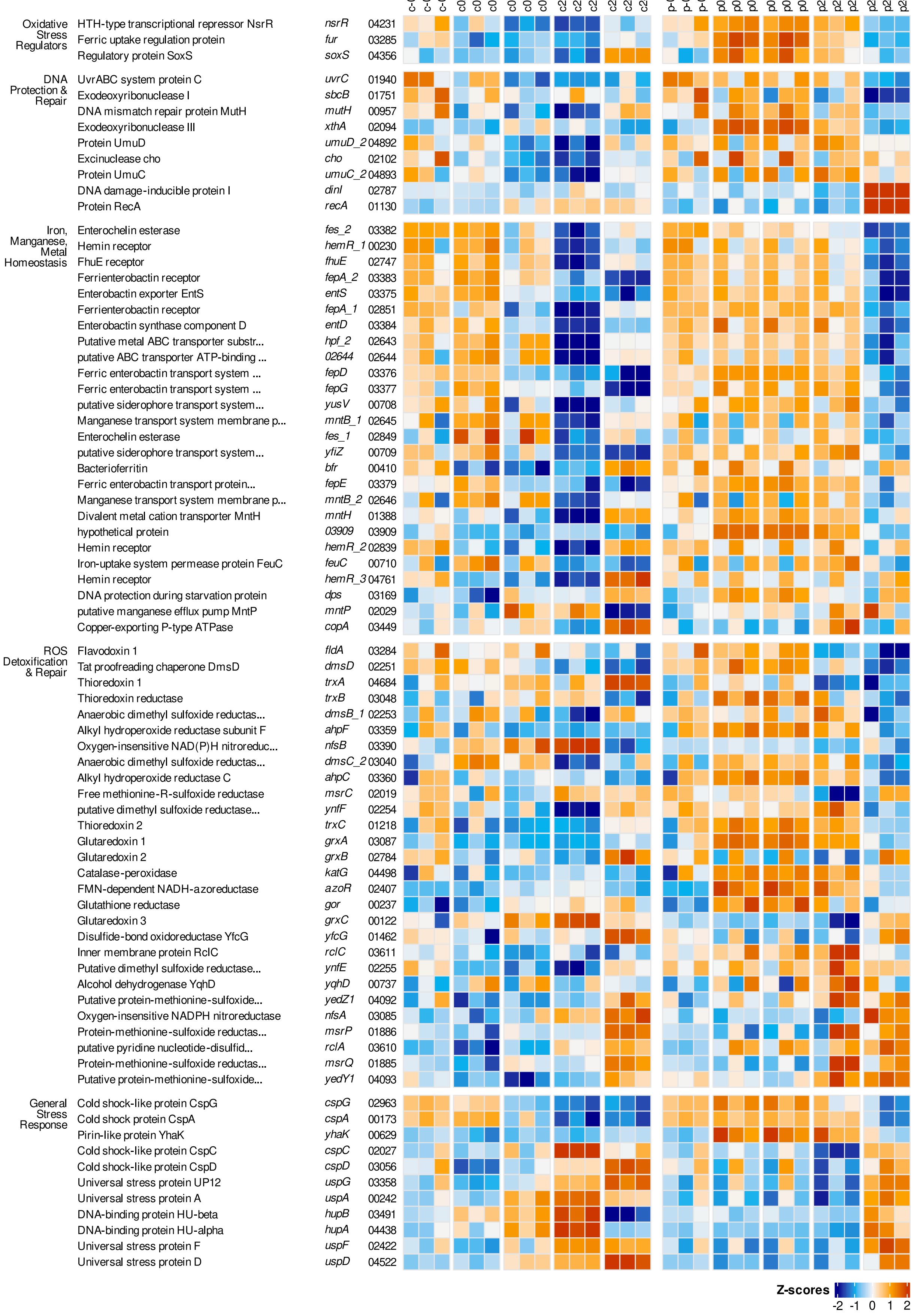

### Fig S2 cont1

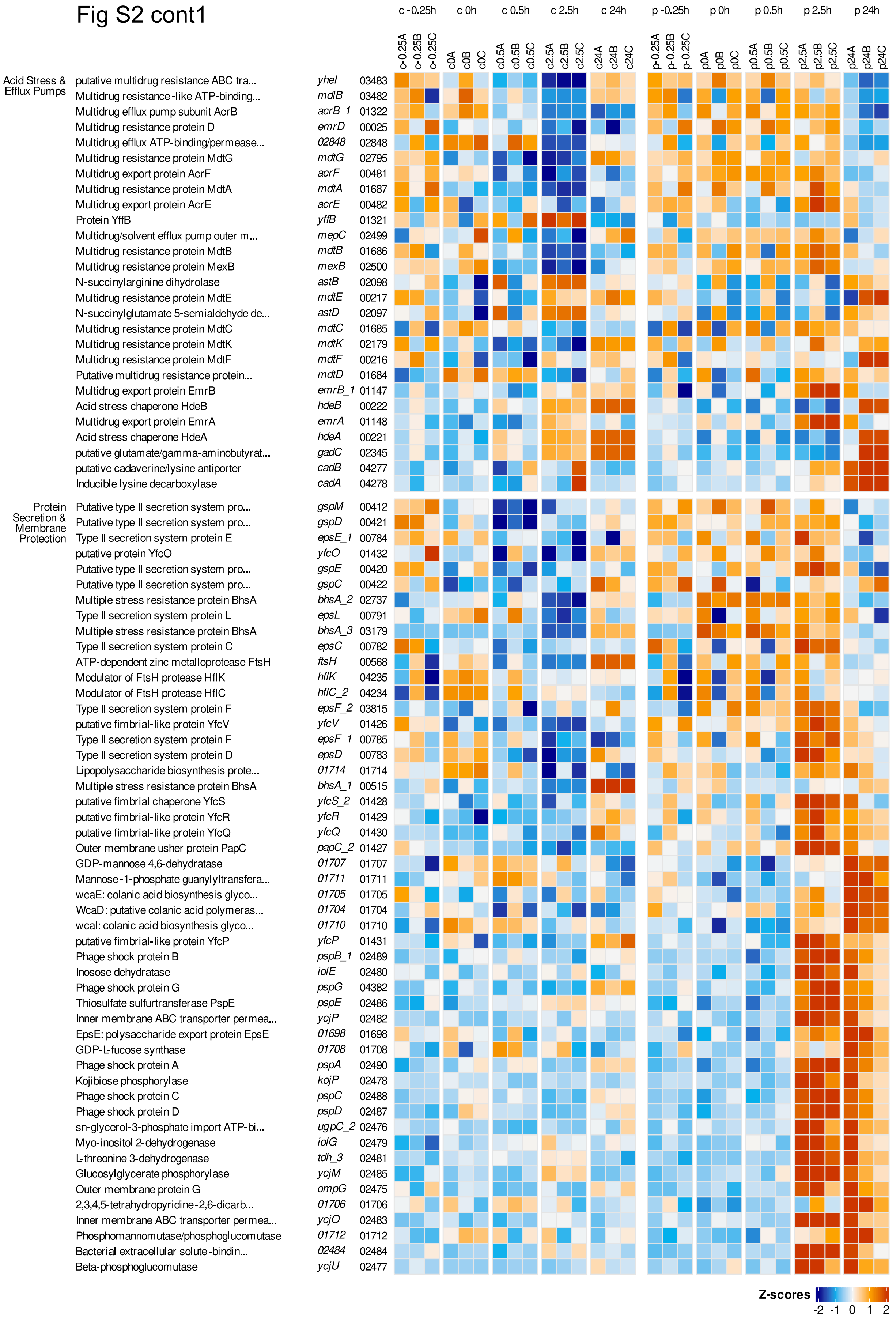
c -0.25h
c 0h
c 0.5h
c 2.5h
c 24h
p -0.25h
p 0h
p 0.5h
p 2.5h
p 24h
c-0.25A c-0.25B c-0.25C c0A c0B c0C c0.5A c0.5B c0.5C c2.5A c2.5B c2.5C c24A c24B c24C p-0.25A p-0.25B p-0.25C p0A p0B p0C p0.5A p0.5B p0.5C p2.5A p2.5B p2.5C p24A p24B p24C
Acid Stress & Efflux Pumps
putative multidrug resistance ABC tra... yheI 03483
Multidrug resistance-like ATP-binding... mdlB 03482
Multidrug efflux pump subunit AcrB acrB_1 01322
Multidrug resistance protein D emrD 00025
Multidrug efflux ATP-binding/permease... 02848 02848
Multidrug resistance protein MdtG mdtG 02795
Multidrug export protein AcrF acrF 00481
Multidrug resistance protein MdtA mdtA 01687
Multidrug export protein AcrE acrE 00482
Protein YffB yffB 01321
Multidrug/solvent efflux pump outer m... mepC 02499
Multidrug resistance protein MdtB mdtB 01686
Multidrug resistance protein MexB mexB 02500
N-succinylarginine dihydrolase astB 02098
Multidrug resistance protein MdtE mdtE 00217
N-succinylglutamate 5-semialdehyde de... astD 02097
Multidrug resistance protein MdtC mdtC 01685
Multidrug resistance protein MdtK mdtK 02179
Multidrug resistance protein MdtF mdtF 00216
Putative multidrug resistance protein... mdtD 01684
Multidrug export protein EmrB emrB_1 01147
Acid stress chaperone HdeB hdeB 00222
Multidrug export protein EmrA emrA 01148
Acid stress chaperone HdeA hdeA 00221
putative glutamate/gamma-aminobutyrat... gadC 02345
putative cadaverine/lysine antiporter cadB 04277
Inducible lysine decarboxylase cadA 04278
Protein Secretion & Membrane Protection
Putative type II secretion system pro... gspM 00412
Putative type II secretion system pro... gspD 00421
Type II secretion system protein E epsE_1 00784
putative protein YfcO yfcO 01432
Putative type II secretion system pro... gspE 00420
Putative type II secretion system pro... gspC 00422
Multiple stress resistance protein BhsA bhsA_2 02737
Type II secretion system protein L epsL 00791
Multiple stress resistance protein BhsA bhsA_3 03179
Type II secretion system protein C epsC 00782
ATP-dependent zinc metalloprotease FtsH ftsH 00568
Modulator of FtsH protease HflK hflK 04235
Modulator of FtsH protease HflC hflC_2 04234
Type II secretion system protein F epsF_2 03815
putative fimbrial-like protein YfcV yfcV 01426
Type II secretion system protein F epsF_1 00785
Type II secretion system protein D epsD 00783
Lipopolysaccharide biosynthesis prote... 01714 01714
Multiple stress resistance protein BhsA bhsA_1 00515
putative fimbrial chaperone YfcS yfcS_2 01428
putative fimbrial-like protein YfcR yfcR 01429
putative fimbrial-like protein YfcQ yfcQ 01430
Outer membrane usher protein PapC papC_2 01427
GDP-mannose 4,6-dehydratase 01707 01707
Mannose-1-phosphate guanylyltransfera... 01711 01711
wcaE: colanic acid biosynthesis glyco... 01705 01705
WcaD: putative colanic acid polymeras... 01704 01704
wcaI: colanic acid biosynthesis glyco... 01710 01710
putative fimbrial-like protein YfcP yfcP 01431
Phage shock protein B pspB_1 02489
Inosose dehydratase iolE 02480
Phage shock protein G pspG 04382
Thiosulfate sulfurtransferase PspE pspE 02486
Inner membrane ABC transporter permea... ycjP 02482
EpsE: polysaccharide export protein EpsE 01698 01698
GDP-L-fucose synthase 01708 01708
Phage shock protein A pspA 02490
Kojibiose phosphorylase kojP 02478
Phage shock protein C pspC 02488
Phage shock protein D pspD 02487
sn-glycerol-3-phosphate import ATP-bi... ugpC_2 02476
Myo-inositol 2-dehydrogenase iolG 02479
L-threonine 3-dehydrogenase tdh_3 02481
Glucosylglycerate phosphorylase ycjM 02485
Outer membrane protein G ompG 02475
2,3,4,5-tetrahydropyridine-2,6-dicarb... 01706 01706
Inner membrane ABC transporter permea... ycjO 02483
Phosphomannomutase/phosphoglucomutase 01712 01712
Bacterial extracellular solute-bindin... 02484 02484
Beta-phosphoglucomutase ycjU 02477
Z-scores
-2 -1 0 1 2

Fig S2 cont2

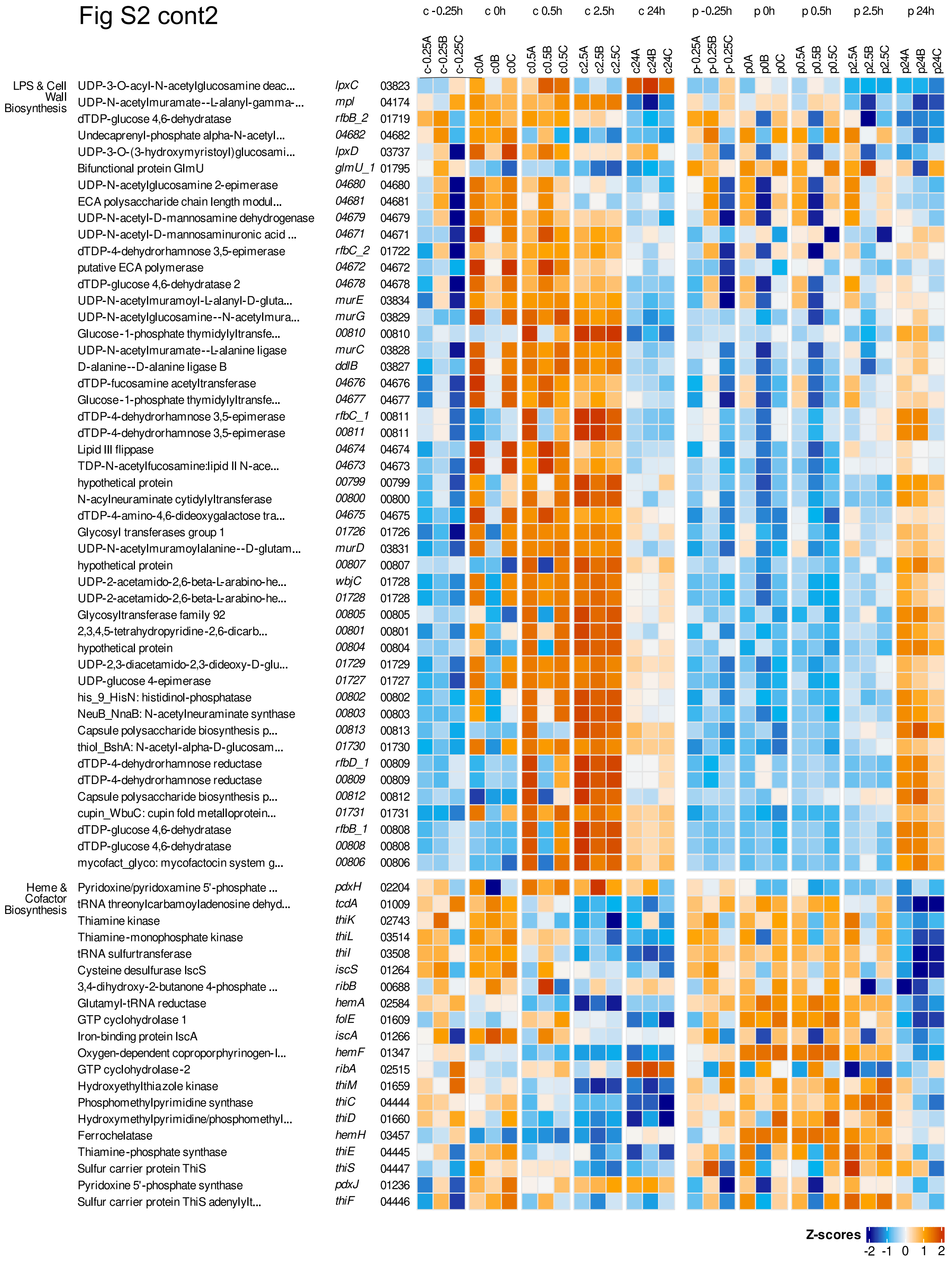
c -0.25h
c 0h
c 0.5h
c 2.5h
c 24h
p -0.25h
p 0h
p 0.5h
p 2.5h
p 24h
c-0.25A c-0.25B c-0.25C c0A c0B c0C c0.5A c0.5B c0.5C c2.5A c2.5B c2.5C c24A c24B c24C p-0.25A p-0.25B p-0.25C p0A p0B p0C p0.5A p0.5B p0.5C p2.5A p2.5B p2.5C p24A p24B p24C
LPS & Cell Wall Biosynthesis
UDP-3-O-acyl-N-acetylglucosamine deac... lpxC 03823
UDP-N-acetylmuramate--L-alanyl-gamma-... mpl 04174
dTDP-glucose 4,6-dehydratase rfbB_2 01719
Undecaprenyl-phosphate alpha-N-acetyl... 04682 04682
UDP-3-O-(3-hydroxymyristoyl)glucosami... lpxD 03737
Bifunctional protein GlmU glmU_1 01795
UDP-N-acetylglucosamine 2-epimerase 04680 04680
ECA polysaccharide chain length modul... 04681 04681
UDP-N-acetyl-D-mannosamine dehydrogenase 04679 04679
UDP-N-acetyl-D-mannosaminuronic acid ... 04671 04671
dTDP-4-dehydrorhamnose 3,5-epimerase rfbC_2 01722
putative ECA polymerase 04672 04672
dTDP-glucose 4,6-dehydratase 2 04678 04678
UDP-N-acetylmuramoyl-L-alanyl-D-gluta... murE 03834
UDP-N-acetylglucosamine--N-acetylmura... murG 03829
Glucose-1-phosphate thymidylyltransfe... 00810 00810
UDP-N-acetylmuramate--L-alanine ligase murC 03828
D-alanine--D-alanine ligase B ddlB 03827
dTDP-fucosamine acetyltransferase 04676 04676
Glucose-1-phosphate thymidylyltransfe... 04677 04677
dTDP-4-dehydrorhamnose 3,5-epimerase rfbC_1 00811
dTDP-4-dehydrorhamnose 3,5-epimerase 00811 00811
Lipid III flippase 04674 04674
TDP-N-acetylfucosamine:lipid II N-ace... 04673 04673
hypothetical protein 00799 00799
N-acylneuraminate cytidylyltransferase 00800 00800
dTDP-4-amino-4,6-dideoxygalactose tra... 04675 04675
Glycosyl transferases group 1 01726 01726
UDP-N-acetylmuramoylalanine--D-glutam... murD 03831
hypothetical protein 00807 00807
UDP-2-acetamido-2,6-beta-L-arabino-he... wbjC 01728
UDP-2-acetamido-2,6-beta-L-arabino-he... 01728 01728
Glycosyltransferase family 92 00805 00805
2,3,4,5-tetrahydropyridine-2,6-dicarb... 00801 00801
hypothetical protein 00804 00804
UDP-2,3-diacetamido-2,3-dideoxy-D-glu... 01729 01729
UDP-glucose 4-epimerase 01727 01727
his_9_HisN: histidinol-phosphatase 00802 00802
NeuB_NnaB: N-acetylneuraminate synthase 00803 00803
Capsule polysaccharide biosynthesis p... 00813 00813
thiol_BshA: N-acetyl-alpha-D-glucosam... 01730 01730
dTDP-4-dehydrorhamnose reductase rfbD_1 00809
dTDP-4-dehydrorhamnose reductase 00809 00809
Capsule polysaccharide biosynthesis p... 00812 00812
cupin_WbuC: cupin fold metalloprotein... 01731 01731
dTDP-glucose 4,6-dehydratase rfbB_1 00808
dTDP-glucose 4,6-dehydratase 00808 00808
mycofact_glyco: mycofactocin system g... 00806 00806
Heme & Cofactor Biosynthesis
Pyridoxine/pyridoxamine 5'-phosphate ... pdxH 02204
tRNA threonylcarbamoyladenosine dehyd... tcdA 01009
Thiamine kinase thiK 02743
Thiamine-monophosphate kinase thiL 03514
tRNA sulfurtransferase thiI 03508
Cysteine desulfurase IscS iscS 01264
3,4-dihydroxy-2-butanone 4-phosphate ... ribB 00688
Glutamyl-tRNA reductase hemA 02584
GTP cyclohydrolase 1 folE 01609
Iron-binding protein IscA iscA 01266
Oxygen-dependent coproporphyrinogen-I... hemF 01347
GTP cyclohydrolase-2 ribA 02515
Hydroxyethylthiazole kinase thiM 01659
Phosphomethylpyrimidine synthase thiC 04444
Hydroxymethylpyrimidine/phosphomethyl... thiD 01660
Ferrochelatase hemH 03457
Thiamine-phosphate synthase thiE 04445
Sulfur carrier protein ThiS thiS 04447
Pyridoxine 5'-phosphate synthase pdxJ 01236
Sulfur carrier protein ThiS adenylylt... thiF 04446
Z-scores
-2 -1 0 1 2

### Fig S2 cont3

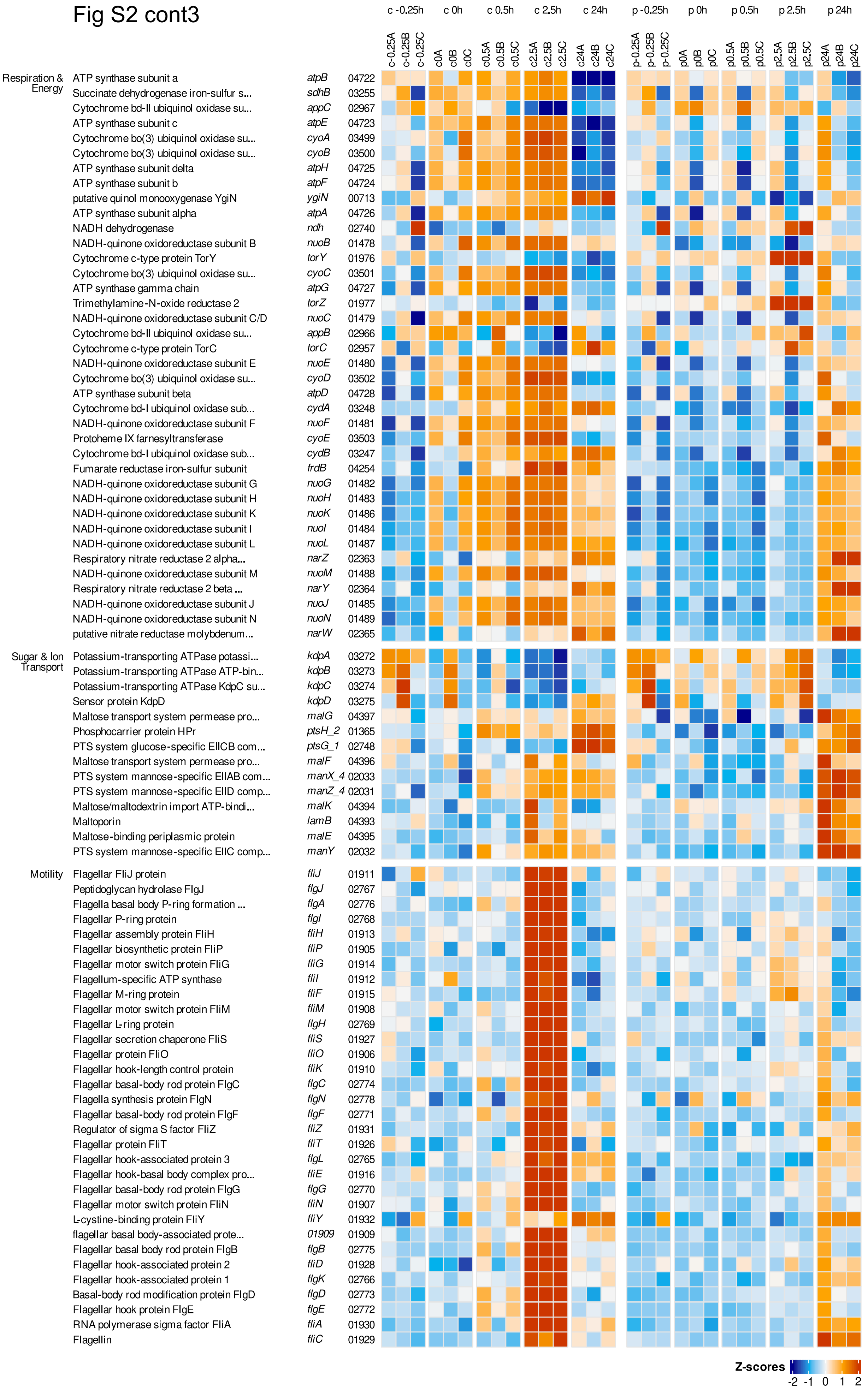

c -0.25h
c 0h
c 0.5h
c 2.5h
c 24h
p -0.25h
p 0h
p 0.5h
p 2.5h
p 24h
c-0.25A c-0.25B c-0.25C c0A c0B c0C c0.5A c0.5B c0.5C c2.5A c2.5B c2.5C c24A c24B c24C p-0.25A p-0.25B p-0.25C p0A p0B p0C p0.5A p0.5B p0.5C p2.5A p2.5B p2.5C p24A p24B p24C
Respiration & Energy
ATP synthase subunit a atpB 04722
Succinate dehydrogenase iron-sulfur s... sdhB 03255
Cytochrome bd-II ubiquinol oxidase su... appC 02967
ATP synthase subunit c atpE 04723
Cytochrome bo(3) ubiquinol oxidase su... cyoA 03499
Cytochrome bo(3) ubiquinol oxidase su... cyoB 03500
ATP synthase subunit delta atpH 04725
ATP synthase subunit b atpF 04724
putative quinol monooxygenase YgiN ygiN 00713
ATP synthase subunit alpha atpA 04726
NADH dehydrogenase ndh 02740
NADH-quinone oxidoreductase subunit B nuoB 01478
Cytochrome c-type protein TorY torY 01976
Cytochrome bo(3) ubiquinol oxidase su... cyoC 03501
ATP synthase gamma chain atpG 04727
Trimethylamine-N-oxide reductase 2 torZ 01977
NADH-quinone oxidoreductase subunit C/D nuoC 01479
Cytochrome bd-II ubiquinol oxidase su... appB 02966
Cytochrome c-type protein TorC torC 02957
NADH-quinone oxidoreductase subunit E nuoE 01480
Cytochrome bo(3) ubiquinol oxidase su... cyoD 03502
ATP synthase subunit beta atpD 04728
Cytochrome bd-I ubiquinol oxidase sub... cydA 03248
NADH-quinone oxidoreductase subunit F nuoF 01481
Protoheme IX farnesyltransferase cyoE 03503
Cytochrome bd-I ubiquinol oxidase sub... cydB 03247
Fumarate reductase iron-sulfur subunit frdB 04254
NADH-quinone oxidoreductase subunit G nuoG 01482
NADH-quinone oxidoreductase subunit H nuoH 01483
NADH-quinone oxidoreductase subunit K nuoK 01486
NADH-quinone oxidoreductase subunit I nuoI 01484
NADH-quinone oxidoreductase subunit L nuoL 01487
Respiratory nitrate reductase 2 alpha... narZ 02363
NADH-quinone oxidoreductase subunit M nuoM 01488
Respiratory nitrate reductase 2 beta ... narY 02364
NADH-quinone oxidoreductase subunit J nuoJ 01485
NADH-quinone oxidoreductase subunit N nuoN 01489
putative nitrate reductase molybdenum... narW 02365
Sugar & Ion Transport
Potassium-transporting ATPase potassi... kdpA 03272
Potassium-transporting ATPase ATP-bin... kdpB 03273
Potassium-transporting ATPase KdpC su... kdpC 03274
Sensor protein KdpD kdpD 03275
Maltose transport system permease pro... malG 04397
Phosphocarrier protein HPr ptsH_2 01365
PTS system glucose-specific EIICB com... ptsG_1 02748
Maltose transport system permease pro... malF 04396
PTS system mannose-specific EIIAB com... manX_4 02033
PTS system mannose-specific EIID comp... manZ_4 02031
Maltose/maltodextrin import ATP-bindi... malK 04394
Maltoporin lamB 04393
Maltose-binding periplasmic protein malE 04395
PTS system mannose-specific EIIC comp... manY 02032
Motility
Flagellar FliJ protein fliJ 01911
Peptidoglycan hydrolase FlgJ flgJ 02767
Flagella basal body P-ring formation ... flgA 02776
Flagellar P-ring protein flgI 02768
Flagellar assembly protein FliH fliH 01913
Flagellar biosynthetic protein FliP fliP 01905
Flagellar motor switch protein FliG fliG 01914
Flagellum-specific ATP synthase fliI 01912
Flagellar M-ring protein fliF 01915
Flagellar motor switch protein FliM fliM 01908
Flagellar L-ring protein flgH 02769
Flagellar secretion chaperone FliS fliS 01927
Flagellar protein FliO fliO 01906
Flagellar hook-length control protein fliK 01910
Flagellar basal-body rod protein FlgC flgC 02774
Flagella synthesis protein FlgN flgN 02778
Flagellar basal-body rod protein FlgF flgF 02771
Regulator of sigma S factor FliZ fliZ 01931
Flagellar protein FliT fliT 01926
Flagellar hook-associated protein 3 flgL 02765
Flagellar hook-basal body complex pro... fliE 01916
Flagellar basal-body rod protein FlgG flgG 02770
Flagellar motor switch protein FliN fliN 01907
L-cystine-binding protein FliY fliY 01932
flagellar basal body-associated prote... 01909 01909
Flagellar basal body rod protein FlgB flgB 02775
Flagellar hook-associated protein 2 fliD 01928
Flagellar hook-associated protein 1 flgK 02766
Basal-body rod modification protein FlgD flgD 02773
Flagellar hook protein FlgE flgE 02772
RNA polymerase sigma factor FliA fliA 01930
Flagellin fliC 01929
Z-scores
-2 -1 0 1 2

Fig S2 cont4

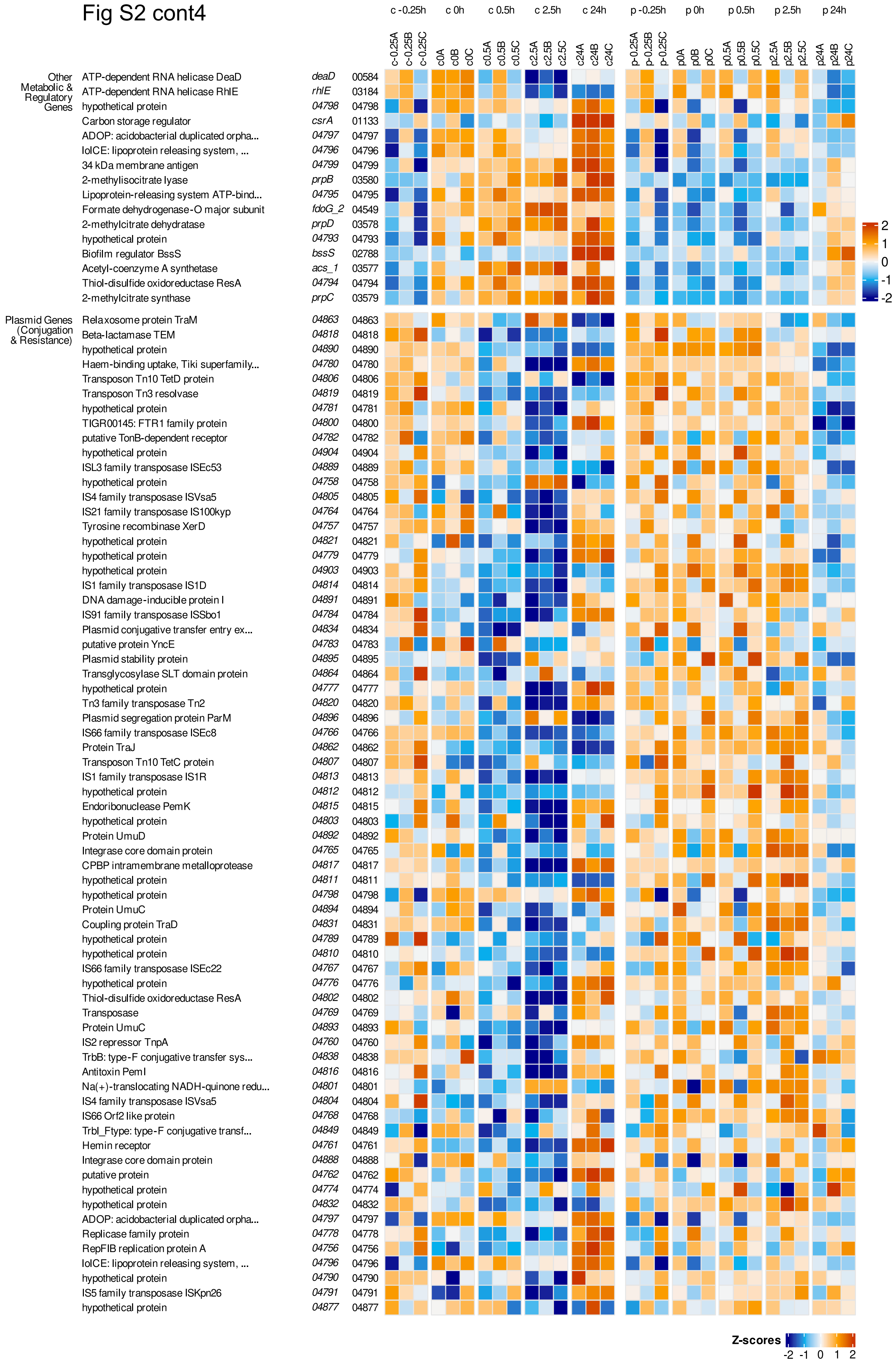

c -0.25h
c 0h
c 0.5h
c 2.5h
c 24h
p -0.25h
p 0h
p 0.5h
p 2.5h
p 24h
c-0.25A c-0.25B c-0.25C c0A c0B c0C c0.5A c0.5B c0.5C c2.5A c2.5B c2.5C c24A c24B c24C p-0.25A p-0.25B p-0.25C p0A p0B p0C p0.5A p0.5B p0.5C p2.5A p2.5B p2.5C p24A p24B p24C
Other Metabolic & Regulatory Genes
ATP-dependent RNA helicase DeaD deaD 00584
ATP-dependent RNA helicase RhlE rhlE 03184
hypothetical protein 04798 04798
Carbon storage regulator csrA 01133
ADOP: acidobacterial duplicated orpha... 04797 04797
lolCE: lipoprotein releasing system, ... 04796 04796
34 kDa membrane antigen 04799 04799
2-methylisocitrate lyase prpB 03580
Lipoprotein-releasing system ATP-bind... 04795 04795
Formate dehydrogenase-O major subunit fdoG_2 04549
2-methylcitrate dehydratase prpD 03578
hypothetical protein 04793 04793
Biofilm regulator BssS bssS 02788
Acetyl-coenzyme A synthetase acs_1 03577
Thiol-disulfide oxidoreductase ResA 04794 04794
2-methylcitrate synthase prpC 03579
Plasmid Genes (Conjugation & Resistance)
Relaxosome protein TraM 04863 04863
Beta-lactamase TEM 04818 04818
hypothetical protein 04890 04890
Haem-binding uptake, Tiki superfamily... 04780 04780
Transposon Tn10 TetD protein 04806 04806
Transposon Tn3 resolvase 04819 04819
hypothetical protein 04781 04781
TIGR00145: FTR1 family protein 04800 04800
putative TonB-dependent receptor 04782 04782
hypothetical protein 04904 04904
ISL3 family transposase ISEc53 04889 04889
hypothetical protein 04758 04758
IS4 family transposase ISVsa5 04805 04805
IS21 family transposase IS100kyp 04764 04764
Tyrosine recombinase XerD 04757 04757
hypothetical protein 04821 04821
hypothetical protein 04779 04779
hypothetical protein 04903 04903
IS1 family transposase IS1D 04814 04814
DNA damage-inducible protein I 04891 04891
IS91 family transposase ISSbo1 04784 04784
Plasmid conjugative transfer entry ex... 04834 04834
putative protein YncE 04783 04783
Plasmid stability protein 04895 04895
Transglycosylase SLT domain protein 04864 04864
hypothetical protein 04777 04777
Tn3 family transposase Tn2 04820 04820
Plasmid segregation protein ParM 04896 04896
IS66 family transposase ISEc8 04766 04766
Protein TraJ 04862 04862
Transposon Tn10 TetC protein 04807 04807
IS1 family transposase IS1R 04813 04813
hypothetical protein 04812 04812
Endoribonuclease PemK 04815 04815
hypothetical protein 04803 04803
Protein UmuD 04892 04892
Integrase core domain protein 04765 04765
CPBP intramembrane metalloprotease 04817 04817
hypothetical protein 04811 04811
hypothetical protein 04798 04798
Protein UmuC 04894 04894
Coupling protein TraD 04831 04831
hypothetical protein 04789 04789
hypothetical protein 04810 04810
IS66 family transposase ISEc22 04767 04767
hypothetical protein 04776 04776
Thiol-disulfide oxidoreductase ResA 04802 04802
Transposase 04769 04769
Protein UmuC 04893 04893
IS2 repressor TnpA 04760 04760
TrbB: type-F conjugative transfer sys... 04838 04838
Antitoxin PemI 04816 04816
Na(+)-translocating NADH-quinone redu... 04801 04801
IS4 family transposase ISVsa5 04804 04804
IS66 Orf2 like protein 04768 04768
TrbI_Ftype: type-F conjugative transf... 04849 04849
Hemin receptor 04761 04761
Integrase core domain protein 04888 04888
putative protein 04762 04762
hypothetical protein 04774 04774
hypothetical protein 04832 04832
ADOP: acidobacterial duplicated orpha... 04797 04797
Replicase family protein 04778 04778
RepFIB replication protein A 04756 04756
lolCE: lipoprotein releasing system, ... 04796 04796
hypothetical protein 04790 04790
IS5 family transposase ISKpn26 04791 04791
hypothetical protein 04877 04877
2
1
0
-1
-2
Z-scores
-2 -1 0 1 2

### Fig S2 cont5

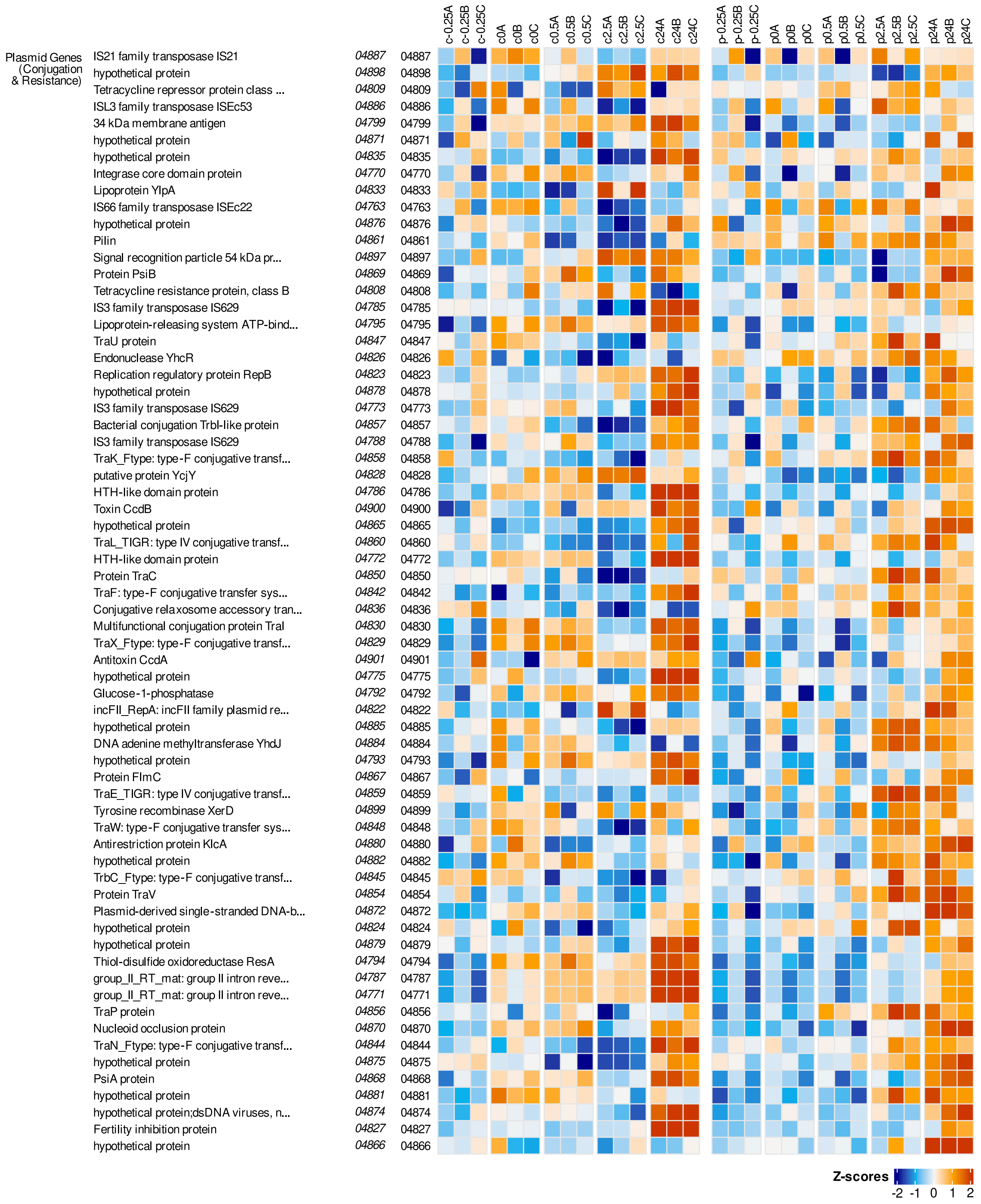
c -0.25h
c 0h
c 0.5h
c 2.5h
c 24h
p -0.25h
p 0h
p 0.5h
p 2.5h
p 24h
c-0.25A c-0.25B c-0.25C
c0A c0B c0C
c0.5A c0.5B c0.5C
c2.5A c2.5B c2.5C
c24A c24B c24C
p-0.25A p-0.25B p-0.25C
p0A p0B p0C
p0.5A p0.5B p0.5C
p2.5A p2.5B p2.5C
p24A p24B p24C
Plasmid Genes (Conjugation & Resistance)
IS21 family transposase IS21 04887 04887
hypothetical protein 04898 04898
Tetracycline repressor protein class ... 04809 04809
ISL3 family transposase ISEc53 04886 04886
34 kDa membrane antigen 04799 04799
hypothetical protein 04871 04871
hypothetical protein 04835 04835
Integrase core domain protein 04770 04770
Lipoprotein YlpA 04833 04833
IS66 family transposase ISEc22 04763 04763
hypothetical protein 04876 04876
Pilin 04861 04861
Signal recognition particle 54 kDa pr... 04897 04897
Protein PsiB 04869 04869
Tetracycline resistance protein, class B 04808 04808
IS3 family transposase IS629 04785 04785
Lipoprotein-releasing system ATP-bind... 04795 04795
TraU protein 04847 04847
Endonuclease YhcR 04826 04826
Replication regulatory protein RepB 04823 04823
hypothetical protein 04878 04878
IS3 family transposase IS629 04773 04773
Bacterial conjugation TrbI-like protein 04857 04857
IS3 family transposase IS629 04788 04788
TraK_Ftype: type-F conjugative transf... 04858 04858
putative protein YcjY 04828 04828
HTH-like domain protein 04786 04786
Toxin CcdB 04900 04900
hypothetical protein 04865 04865
TraL_TIGR: type IV conjugative transf... 04860 04860
HTH-like domain protein 04772 04772
Protein TraC 04850 04850
TraF: type-F conjugative transfer sys... 04842 04842
Conjugative relaxosome accessory tran... 04836 04836
Multifunctional conjugation protein TraI 04830 04830
TraX_Ftype: type-F conjugative transf... 04829 04829
Antitoxin CcdA 04901 04901
hypothetical protein 04775 04775
Glucose-1-phosphatase 04792 04792
incFII_RepA: incFII family plasmid re... 04822 04822
hypothetical protein 04885 04885
DNA adenine methyltransferase YhdJ 04884 04884
hypothetical protein 04793 04793
Protein FlmC 04867 04867
TraE_TIGR: type IV conjugative transf... 04859 04859
Tyrosine recombinase XerD 04899 04899
TraW: type-F conjugative transfer sys... 04848 04848
Antirestriction protein KlcA 04880 04880
hypothetical protein 04882 04882
TrbC_Ftype: type-F conjugative transf... 04845 04845
Protein TraV 04854 04854
Plasmid-derived single-stranded DNA-b... 04872 04872
hypothetical protein 04824 04824
hypothetical protein 04879 04879
Thiol-disulfide oxidoreductase ResA 04794 04794
group_II_RT_mat: group II intron reve... 04787 04787
group_II_RT_mat: group II intron reve... 04771 04771
TraP protein 04856 04856
Nucleoid occlusion protein 04870 04870
TraN_Ftype: type-F conjugative transf... 04844 04844
hypothetical protein 04875 04875
PsiA protein 04868 04868
hypothetical protein 04881 04881
hypothetical protein;dsDNA viruses, n... 04874 04874
Fertility inhibition protein 04827 04827
hypothetical protein 04866 04866
Z-scores
-2 -1 0 1 2

### Fig S2 cont6

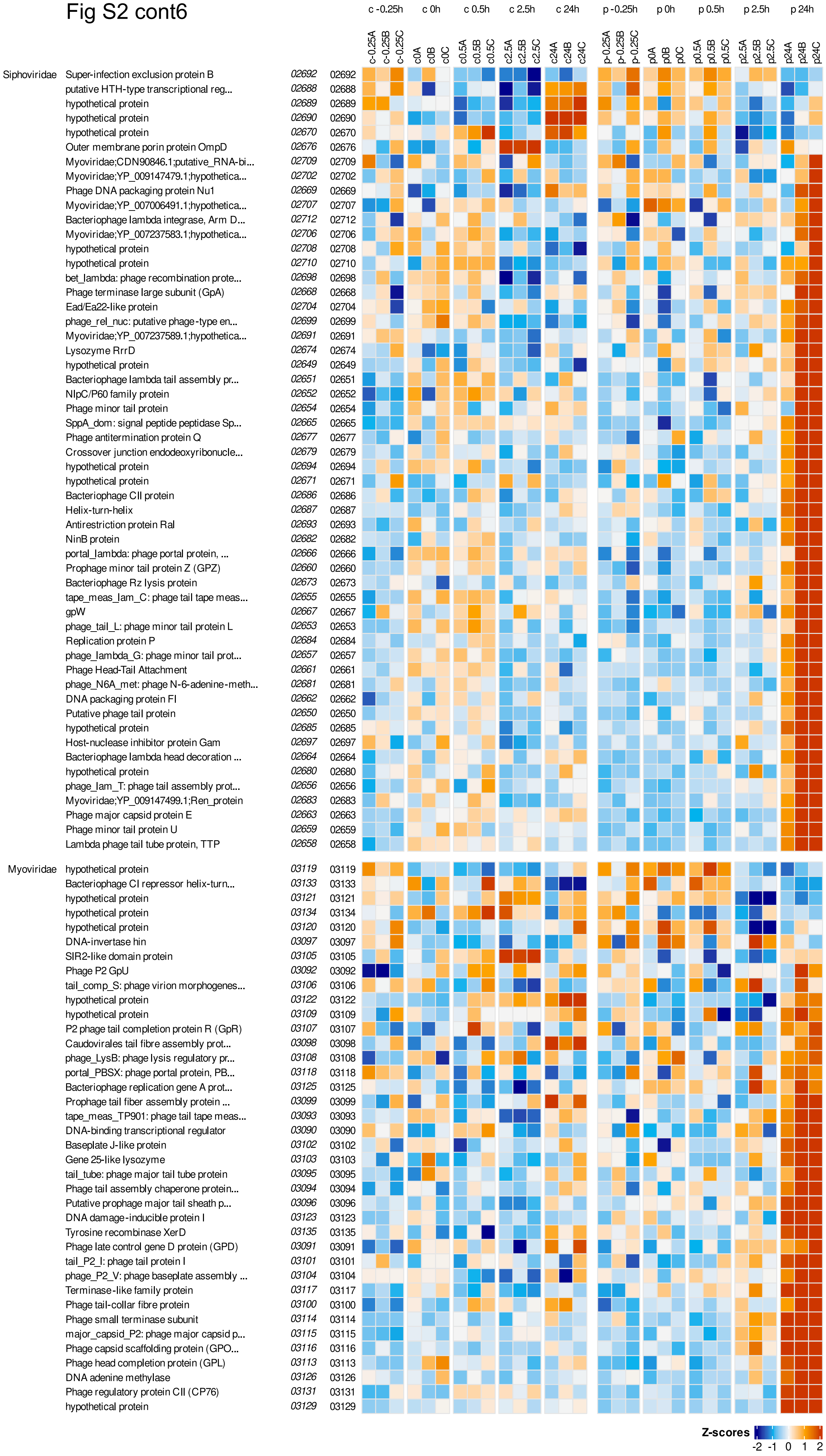
c -0.25h
c 0h
c 0.5h
c 2.5h
c 24h
p -0.25h
p 0h
p 0.5h
p 2.5h
p 24h
c-0.25A
c-0.25B
c-0.25C
c0A
c0B
c0C
c0.5A
c0.5B
c0.5C
c2.5A
c2.5B
c2.5C
c24A
c24B
c24C
p-0.25A
p-0.25B
p-0.25C
p0A
p0B
p0C
p0.5A
p0.5B
p0.5C
p2.5A
p2.5B
p2.5C
p24A
p24B
p24C
Siphoviridae
Super-infection exclusion protein B 02692 02692
putative HTH-type transcriptional reg... 02688 02688
hypothetical protein 02689 02689
hypothetical protein 02690 02690
hypothetical protein 02670 02670
Outer membrane porin protein OmpD 02676 02676
Myoviridae;CDN90846.1;putative_RNA-bi... 02709 02709
Myoviridae;YP_009147479.1;hypothetica... 02702 02702
Phage DNA packaging protein Nu1 02669 02669
Myoviridae;YP_007006491.1;hypothetica... 02707 02707
Bacteriophage lambda integrase, Arm D... 02712 02712
Myoviridae;YP_007237583.1;hypothetica... 02706 02706
hypothetical protein 02708 02708
hypothetical protein 02710 02710
bet_lambda: phage recombination prote... 02698 02698
Phage terminase large subunit (GpA) 02668 02668
Ead/Ea22-like protein 02704 02704
phage_rel_nuc: putative phage-type en... 02699 02699
Myoviridae;YP_007237589.1;hypothetica... 02691 02691
Lysozyme RrrD 02674 02674
hypothetical protein 02649 02649
Bacteriophage lambda tail assembly pr... 02651 02651
NlpC/P60 family protein 02652 02652
Phage minor tail protein 02654 02654
SppA_dom: signal peptide peptidase Sp... 02665 02665
Phage antitermination protein Q 02677 02677
Crossover junction endodeoxyribonucle... 02679 02679
hypothetical protein 02694 02694
hypothetical protein 02671 02671
Bacteriophage CII protein 02686 02686
Helix-turn-helix 02687 02687
Antirestriction protein Ral 02693 02693
NinB protein 02682 02682
portal_lambda: phage portal protein, ... 02666 02666
Prophage minor tail protein Z (GPZ) 02660 02660
Bacteriophage Rz lysis protein 02673 02673
tape_meas_lam_C: phage tail tape meas... 02655 02655
gpW 02667 02667
phage_tail_L: phage minor tail protein L 02653 02653
Replication protein P 02684 02684
phage_lambda_G: phage minor tail prot... 02657 02657
Phage Head-Tail Attachment 02661 02661
phage_N6A_met: phage N-6-adenine-meth... 02681 02681
DNA packaging protein FI 02662 02662
Putative phage tail protein 02650 02650
hypothetical protein 02685 02685
Host-nuclease inhibitor protein Gam 02697 02697
Bacteriophage lambda head decoration ... 02664 02664
hypothetical protein 02680 02680
phage_lam_T: phage tail assembly prot... 02656 02656
Myoviridae;YP_009147499.1;Ren_protein 02683 02683
Phage major capsid protein E 02663 02663
Phage minor tail protein U 02659 02659
Lambda phage tail tube protein, TTP 02658 02658
Myoviridae
hypothetical protein 03119 03119
Bacteriophage CI repressor helix-turn... 03133 03133
hypothetical protein 03121 03121
hypothetical protein 03134 03134
hypothetical protein 03120 03120
DNA-invertase hin 03097 03097
SIR2-like domain protein 03105 03105
Phage P2 GpU 03092 03092
tail_comp_S: phage virion morphogenes... 03106 03106
hypothetical protein 03122 03122
hypothetical protein 03109 03109
P2 phage tail completion protein R (GpR) 03107 03107
Caudovirales tail fibre assembly prot... 03098 03098
phage_LysB: phage lysis regulatory pr... 03108 03108
portal_PBSX: phage portal protein, PB... 03118 03118
Bacteriophage replication gene A prot... 03125 03125
Prophage tail fiber assembly protein ... 03099 03099
tape_meas_TP901: phage tail tape meas... 03093 03093
DNA-binding transcriptional regulator 03090 03090
Baseplate J-like protein 03102 03102
Gene 25-like lysozyme 03103 03103
tail_tube: phage major tail tube protein 03095 03095
Phage tail assembly chaperone protein... 03094 03094
Putative prophage major tail sheath p... 03096 03096
DNA damage-inducible protein I 03123 03123
Tyrosine recombinase XerD 03135 03135
Phage late control gene D protein (GPD) 03091 03091
tail_P2_I: phage tail protein I 03101 03101
phage_P2_V: phage baseplate assembly ... 03104 03104
Terminase-like family protein 03117 03117
Phage tail-collar fibre protein 03100 03100
Phage small terminase subunit 03114 03114
major_capsid_P2: phage major capsid p... 03115 03115
Phage capsid scaffolding protein (GPO... 03116 03116
Phage head completion protein (GPL) 03113 03113
DNA adenine methylase 03126 03126
Phage regulatory protein CII (CP76) 03131 03131
hypothetical protein 03129 03129
Z-scores
-2 -1 0 1 2

### Fig S3 - supplements Fig 5
### RNASeq heatmap regulons - supp

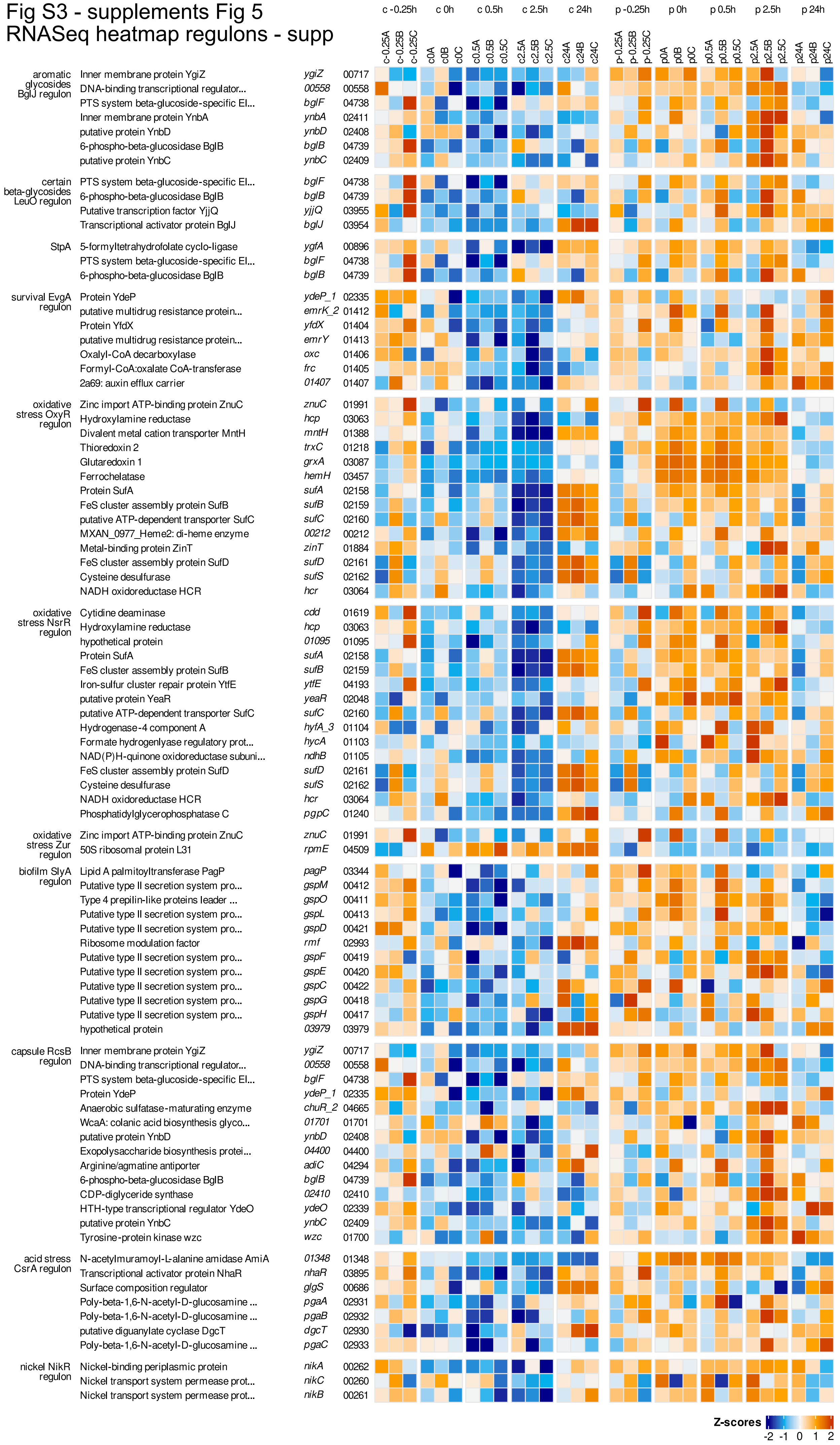

Fig S4 - supplements Fig 6
volcano plots proteins three panels - supp

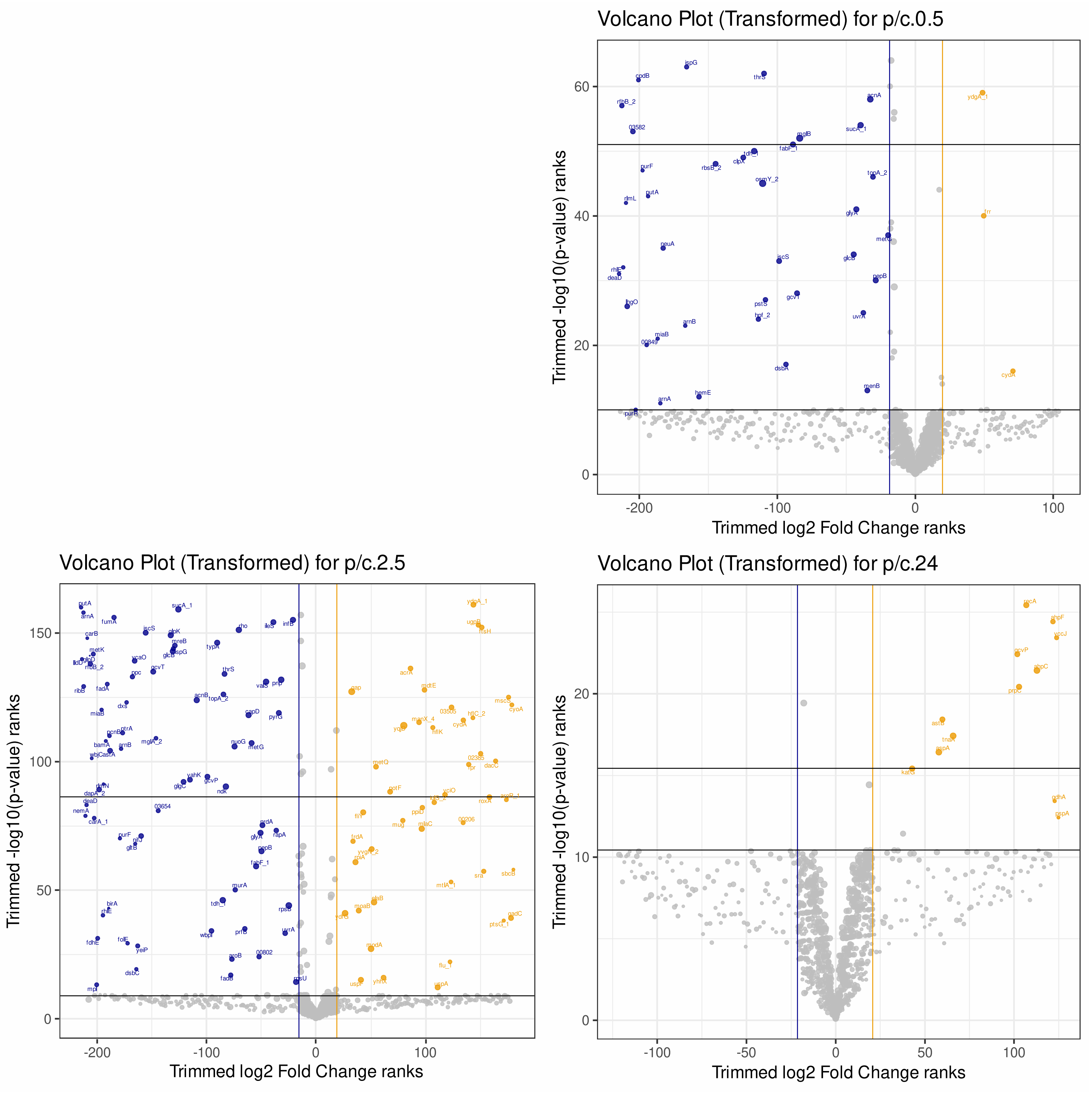

### Fig S5 - supplements protein results
proteins - induced - supp

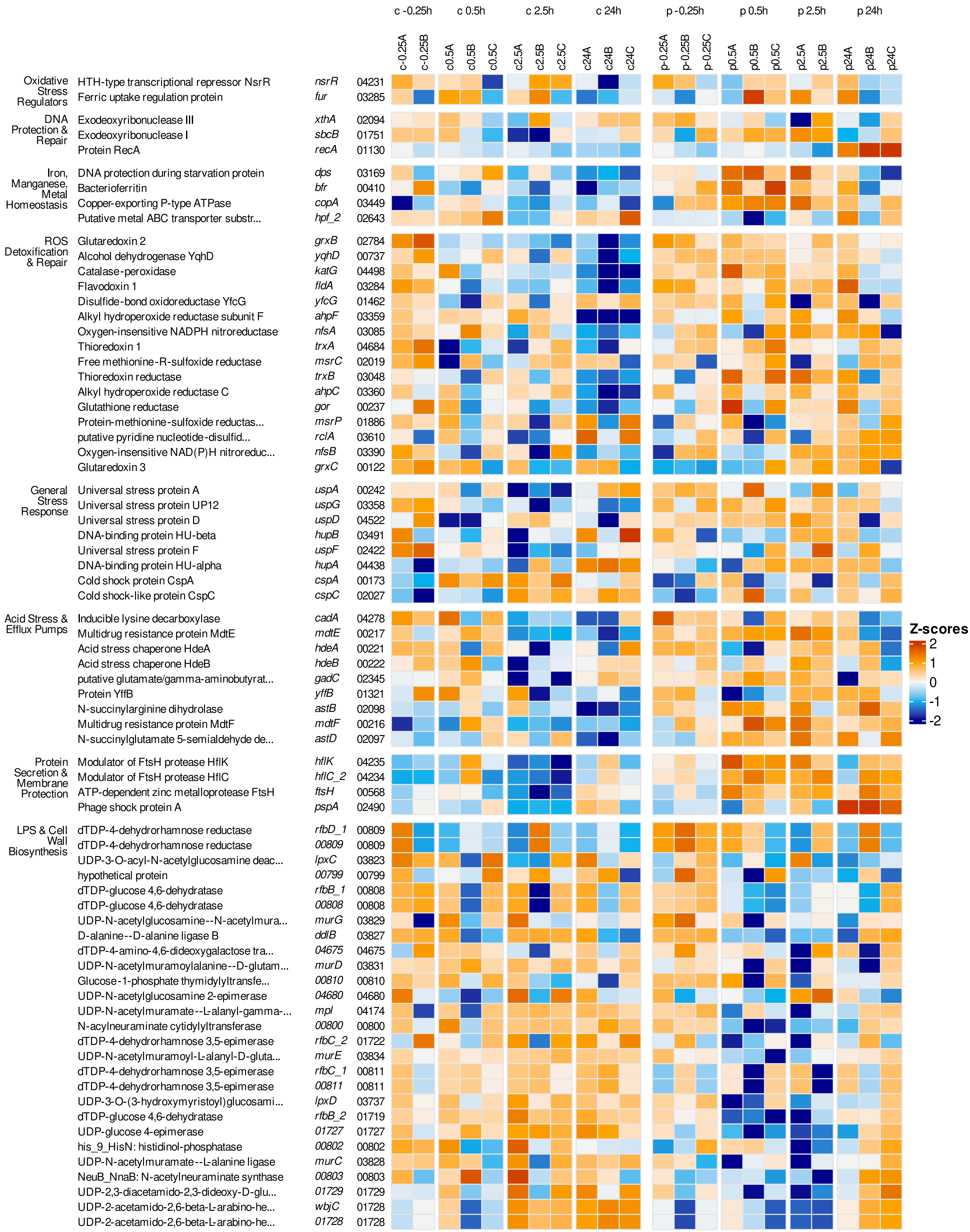

### Fig S5 - cont1

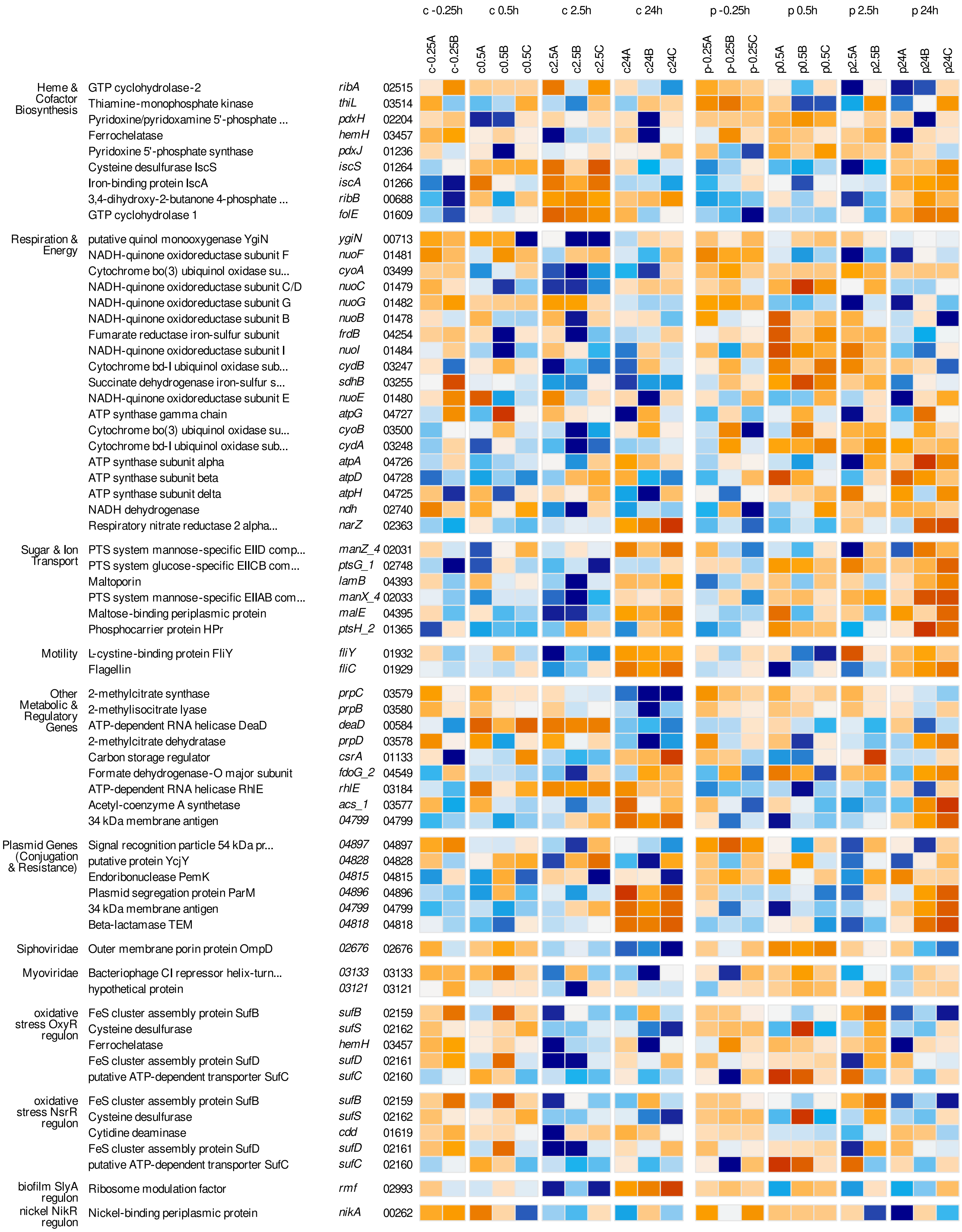
c -0.25h
c 0.5h
c 2.5h
c 24h
p -0.25h
p 0.5h
p 2.5h
p 24h
c-0.25A
c-0.25B
c0.5A
c0.5B
c0.5C
c2.5A
c2.5B
c2.5C
c24A
c24B
c24C
p-0.25A
p-0.25B
p-0.25C
p0.5A
p0.5B
p0.5C
p2.5A
p2.5B
p24A
p24B
p24C
Heme & Cofactor Biosynthesis
GTP cyclohydrolase-2 ribA 02515
Thiamine-monophosphate kinase thiL 03514
Pyridoxine/pyridoxamine 5'-phosphate ... pdxH 02204
Ferrochelatase hemH 03457
Pyridoxine 5'-phosphate synthase pdxJ 01236
Cysteine desulfurase IscS iscS 01264
Iron-binding protein IscA iscA 01266
3,4-dihydroxy-2-butanone 4-phosphate ... ribB 00688
GTP cyclohydrolase 1 folE 01609
Respiration & Energy
putative quinol monooxygenase YgiN ygiN 00713
NADH-quinone oxidoreductase subunit F nuoF 01481
Cytochrome bo(3) ubiquinol oxidase su... cyoA 03499
NADH-quinone oxidoreductase subunit C/D nuoC 01479
NADH-quinone oxidoreductase subunit G nuoG 01482
NADH-quinone oxidoreductase subunit B nuoB 01478
Fumarate reductase iron-sulfur subunit frdB 04254
NADH-quinone oxidoreductase subunit I nuoI 01484
Cytochrome bd-I ubiquinol oxidase sub... cydB 03247
Succinate dehydrogenase iron-sulfur s... sdhB 03255
NADH-quinone oxidoreductase subunit E nuoE 01480
ATP synthase gamma chain atpG 04727
Cytochrome bo(3) ubiquinol oxidase su... cyoB 03500
Cytochrome bd-I ubiquinol oxidase sub... cydA 03248
ATP synthase subunit alpha atpA 04726
ATP synthase subunit beta atpD 04728
ATP synthase subunit delta atpH 04725
NADH dehydrogenase ndh 02740
Respiratory nitrate reductase 2 alpha... narZ 02363
Sugar & Ion Transport
PTS system mannose-specific EIID comp... manZ_4 02031
PTS system glucose-specific EIICB com... ptsG_1 02748
Maltoporin lamB 04393
PTS system mannose-specific EIIAB com... manX_4 02033
Maltose-binding periplasmic protein malE 04395
Phosphocarrier protein HPr ptsH_2 01365
Motility
L-cystine-binding protein FliY fliY 01932
Flagellin fliC 01929
Other Metabolic & Regulatory Genes
2-methylcitrate synthase prpC 03579
2-methylisocitrate lyase prpB 03580
ATP-dependent RNA helicase DeaD deaD 00584
2-methylcitrate dehydratase prpD 03578
Carbon storage regulator csrA 01133
Formate dehydrogenase-O major subunit fdoG_2 04549
ATP-dependent RNA helicase RhlE rhlE 03184
Acetyl-coenzyme A synthetase acs_1 03577
34 kDa membrane antigen 04799 04799
Plasmid Genes (Conjugation & Resistance)
Signal recognition particle 54 kDa pr... 04897 04897
putative protein YcjY 04828 04828
Endoribonuclease PemK 04815 04815
Plasmid segregation protein ParM 04896 04896
34 kDa membrane antigen 04799 04799
Beta-lactamase TEM 04818 04818
Siphoviridae
Outer membrane porin protein OmpD 02676 02676
Myoviridae
Bacteriophage CI repressor helix-turn... 03133 03133
hypothetical protein 03121 03121
oxidative stress OxyR regulon
FeS cluster assembly protein SufB sufB 02159
Cysteine desulfurase sufS 02162
Ferrochelatase hemH 03457
FeS cluster assembly protein SufD sufD 02161
putative ATP-dependent transporter SufC sufC 02160
oxidative stress NsrR regulon
FeS cluster assembly protein SufB sufB 02159
Cysteine desulfurase sufS 02162
Cytidine deaminase cdd 01619
FeS cluster assembly protein SufD sufD 02161
putative ATP-dependent transporter SufC sufC 02160
biofilm SlyA regulon
Ribosome modulation factor rmf 02993
nickel NikR regulon
Nickel-binding periplasmic protein nikA 00262

Fig S6
Phage genomes - supp

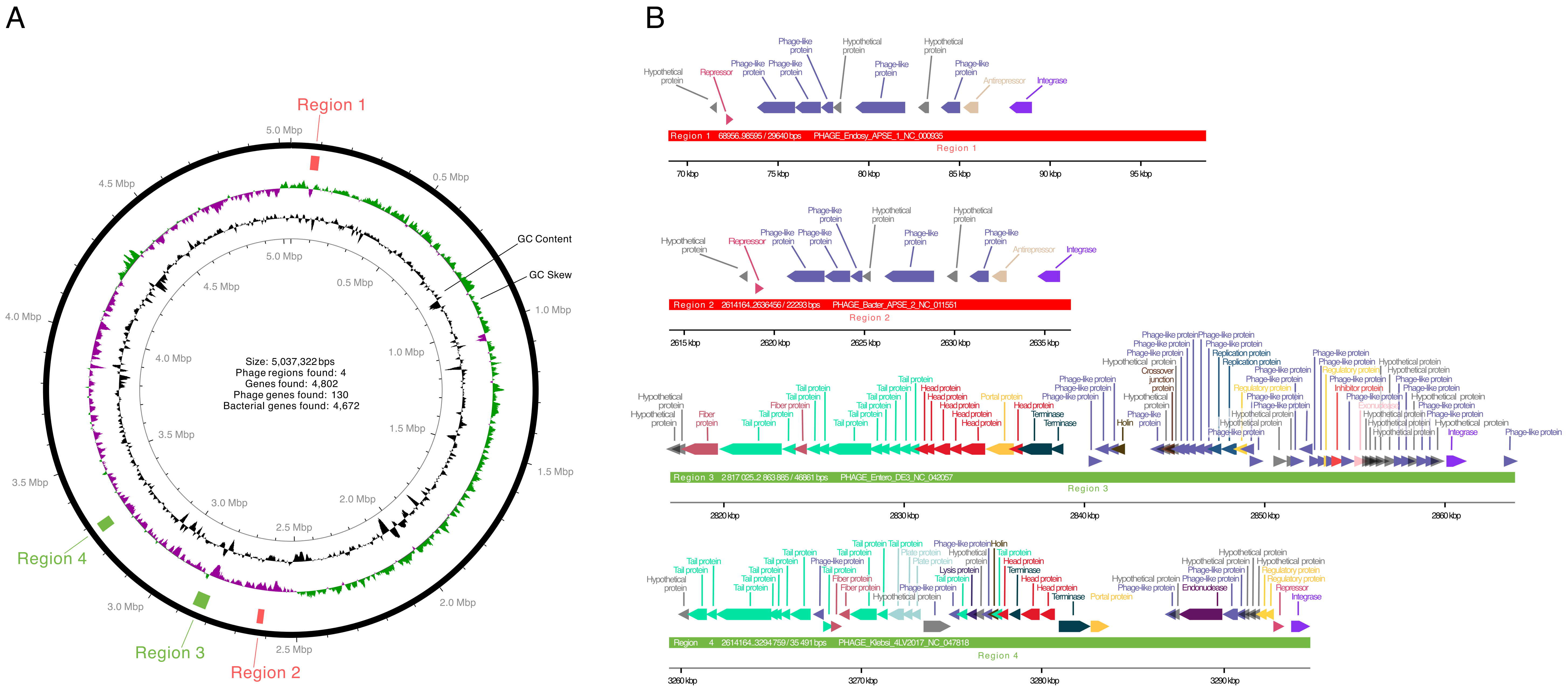
